# The compact genome of the sponge *Oopsacas minuta* (Hexactinellida) is lacking key metazoan core genes

**DOI:** 10.1101/2022.07.26.501511

**Authors:** Sébastien Santini, Quentin Schenkelaars, Cyril Jourda, Marc Duschene, Hassiba Belahbib, Caroline Rocher, Marjorie Selva, Ana Riesgo, Michel Vervoort, Sally P. Leys, Laurent Kodjabachian, André Le Bivic, Carole Borchiellini, Jean-Michel Claverie, Emmanuelle Renard

## Abstract

**Background:** Bilaterian animals today represent 99% of animal biodiversity. Elucidating how bilaterian hallmarks emerged is a central question of animal evo-devo and evolutionary genomics. Studies of non-bilaterian genomes have suggested that the ancestral animal already possessed a diversified developmental toolkit, including some pathways required for bilaterian body plans. Comparing genomes within the early branching metazoan Porifera phylum is key to identify which changes and innovations contributed to the successful transition towards bilaterians.

**Results:** Here, we report the first whole genome comprehensive analysis of a glass sponge, *Oopsacas minuta*, a member of the *Hexactinellida*. Studying this class of sponge is evolutionary relevant because it differs from the three other *Porifera* classes in terms of development, tissue organization, ecology and physiology. Although *O. minuta* does not exhibit drastic body simplifications, its genome is among the smallest animal genomes sequenced so far, surprisingly lacking several metazoan core genes (including Wnt and several key transcription factors). Our study also provided the complete genome of the symbiotic organism dominating the associated microbial community: a new *Thaumarchaeota* species.

**Conclusions:** The genome of the glass sponge *O. minuta* differs from all other available sponge genomes by its compactness and smaller number of predicted proteins. The unexpected losses of numerous genes considered as ancestral and pivotal for metazoan morphogenetic processes most likely reflect the peculiar syncytial organization in this group. Our work further documents the importance of convergence during animal evolution, with multiple emergences of sponge skeleton, electrical signaling and multiciliated cells.

## BACKGROUND

Elucidating the early steps of animal evolution is one of the major challenges of evolutionary biology. One way towards this goal is by comparing genomic data from non-bilaterian animals (Ctenophora, Placozoa, Porifera, Cnidaria)(1-3).

Porifera (sponges) are among the best candidates as sister group of all other animals (4-10) (Fig. 1a) and of particular interest. Although the biology of these animals remains poorly documented (11), its long-lasting (>600 million years) history (12) has resulted in a highly diversified phylum (about 9500 described species (13)), distributed into four classes, with diverse ecological, embryological, cellular and morphological features (14-17) (Fig.1a, 1b). The first sponge genome sequenced (the demosponge *Amphimedon queenslandica*) revealed a size and gene content larger than expected (18-20). However, transcriptomic data from other sponges suggested that this genome was not representative of the diversity of sponges (17,21–26). The other sponge genomes currently available (27,28,29,30,31) illustrate the disparity of sponge genomes in terms of size, features of non-coding regions and gene repertoire (3, 28). However, the lack of a complete genome from the Hexactinellida (“glass sponges”)(Fig.1b) has remained a major gap in our knowledge of Porifera, although partial analyses provided a glance at their gene contents (3,23,26,32,33). A comprehensive description of the genomic features of a glass sponge is needed to precisely evaluate how much they differ from other sponges and to identify the key genomic changes that occurred during sponge diversification into four markedly different classes.

**Figure 1.**
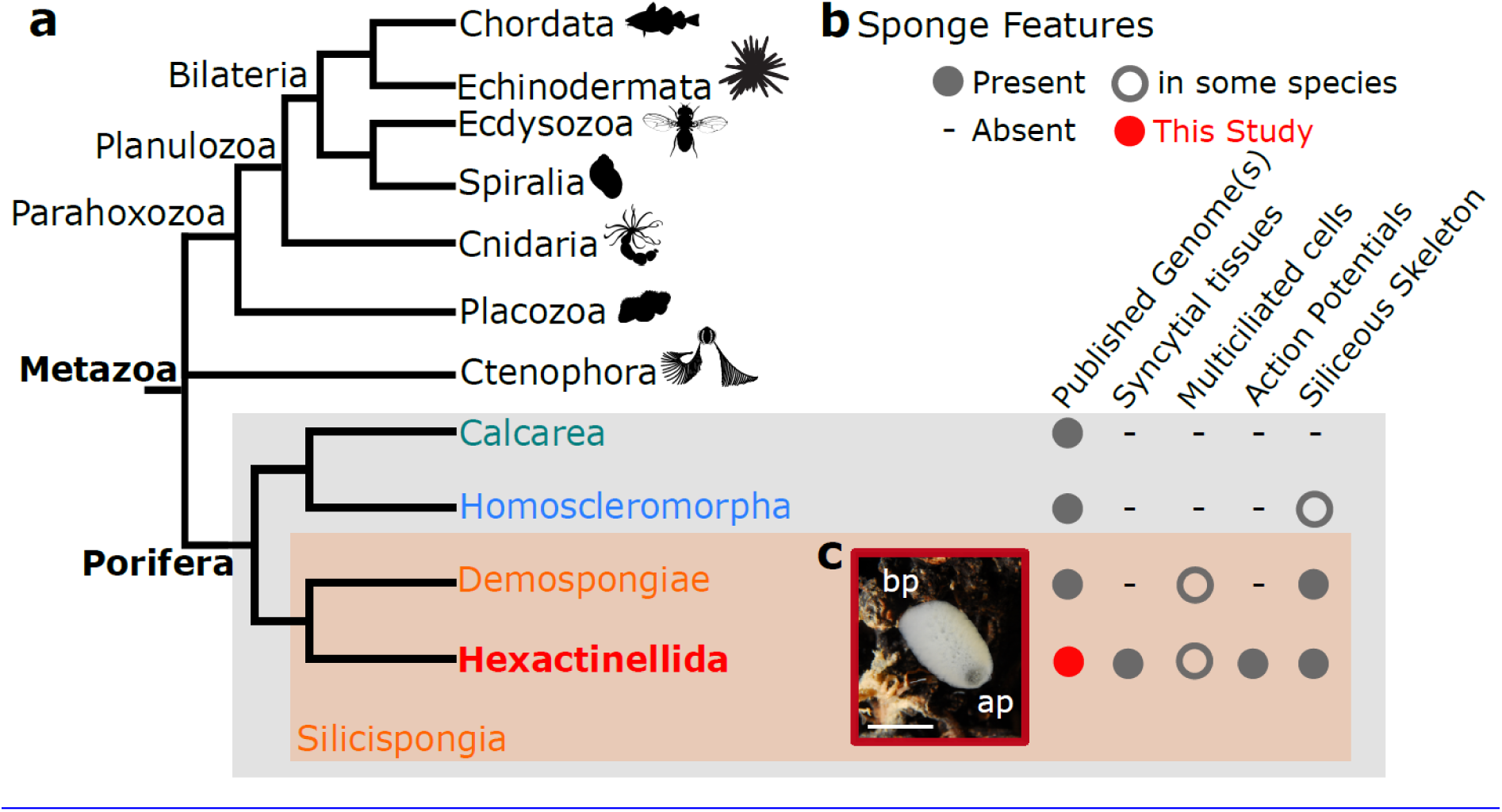
General features of the hexactinellid sponge *Oopsacas minuta*. **a:** Phylogenetic position of *O. minuta* : this species belongs to the phylum Porifera (sponges), one of the best candidates as sister-group to all other Metazoa. Free animal silhouettes were downloaded from PhyloPic (http://phylopic.org/). Among the four lineages of Porifera (> 9,490 species), *O. minuta* belongs to Hexactinellida (Glass Sponges). **b:** Hexactinellida have particular features compared to the three other sponge classes. **c**: *O. minuta* adults inhabit Mediterranean canyons and shallow caves (photo credit Dorian Guillemain), they measure about 1-7cm (scale bar = 1cm) and have a clear basal-apical polarity (ap= apical pole; bp= basal pole).

Here, we provide the first high quality genome sequence of the glass sponge *Oopsacas minuta* (Topsent 1927)(Fig.1c). This species was chosen for its clear taxonomic identification, its accessibility at shallow depths in Mediterranean caves and the availability of previous descriptions of its biological features (34-36). Given the known histological (mainly syncytial organization), physiological (action potential reported) and ecological (coming from deep seas) particularities of *Hexactinellida* (Fig.1b), we focused our bioinformatic analyses on genes involved in development, epithelia and multiciliogenesis, in silicate biogenesis and signal transduction. We also gathered data on the dominant microbiome associated with *O. minuta*.

## RESULTS

### Initial metagenomic data processing

The data obtained from the DNA mixture extracted from non-clonal specimens collected from their natural environment was initially processed as sequences from a low complexity metagenome consisting of the sponge’s nuclear genome, its mitochondrial genome, and of an undescribed population of microorganisms including non-resident (food) and resident (symbionts *sensu lato* (37)) species.

The combination of sequence data (from two complementary platforms Pacific Biosciences and Illumina) yielded an initial dataset of 1759 scaffolds larger than 1kb that were first classified as Eukaryota, Bacteria, Archaea, or virus sequences. Further steps in refining the assembly resulted in three distinctive sets of super-scaffolds associated with large coverage values: the genome of the Thaumarchaeota (coverage=1206), the *O. minuta*’s nuclear genome (coverage=186), and the mitochondrial genome (coverage=455). A number of other scaffolds with smaller coverage were attributed to the residual microbiome of the sponge (Supplementary Table 1). The fully assembled mitochondrial genome was published previously (38).

### The dominance of a Thaumarchaeota symbiont

Data on hexactinellid microbiomes are scarce, and histological observations suggest that bacterial symbionts are rare (36, 39,40).

Of the 1759 scaffolds, 107 scaffolds were assigned to Bacteria (more than 11 phyla) dominated by γ-proteobacteria (Fig. 2a, Supplementary Table 2). In addition, one of the two viral contigs was affiliated with the *Circoviridae*, a family of small single-stranded DNA viruses. Surprisingly for an animal that filters water, viruses have rarely been observed in sponges (41, 42) and few sponge viromes are available (43-46). Nevertheless, *Circoviridae*-related sequences have been reported in two demosponges (47). Known circoviruses tend to be associated with vertebrate hosts and are considered rare in marine invertebrates (48), highlighting the novelty of our finding (49).

**Figure 2.**
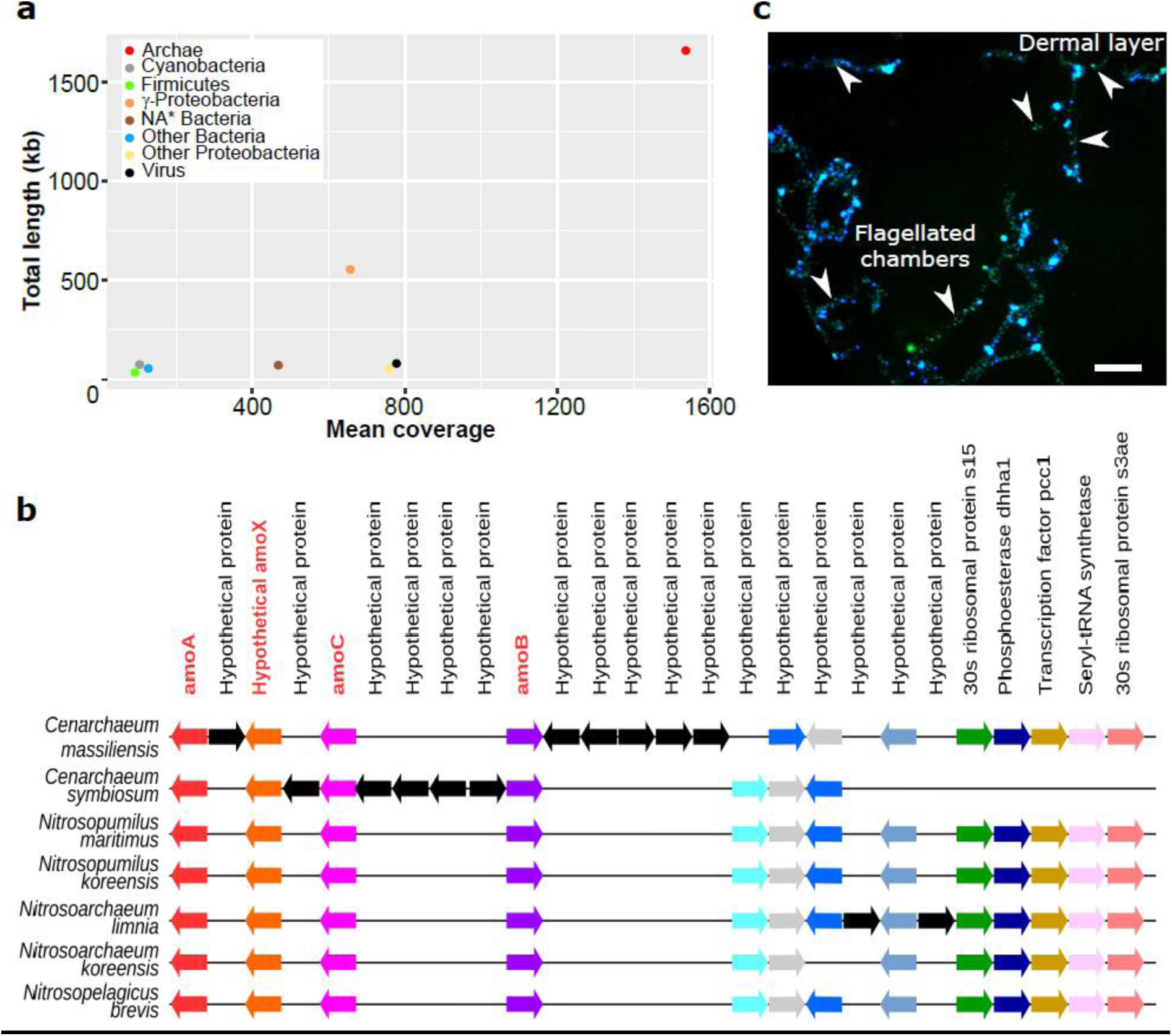
Characterization of the microbiome associated with *O. minuta*. **a:** Relative coverage and sequence size of non-eukaryotic contigs according to their taxa (Supplementary Table 1). The microbiome of *O.minuta* is dominated by a new *Thaumarchaeota* species that we propose to name *Candidatus Cenarchaeum massiliensis*, and various proteobacteria. **b**: Microsynteny analysis of genomic segments bearing an *amo* gene cluster in marine Thaumarchaeota showing that the genome *Candidatus Cenarchaeum massiliensis* presents highly conserved features of other marine Thaumarchaeota (arrows of the same color indicate orthologous genes and black arrows represent genes without orthology relationship in these regions from OrthoMCL analysis). Other data concerning *Ca. C. massiliensis* are available in Supplementary Tables 3 and 4, and Supplementary Fig. 1, 2 and 3. **c:** Localization of *Ca. C. massiliensis* in the tissues of *O. minuta* by Card-FISH using a specific probe (Blue = DAPI staining; green= labeled probe) shows the presence of this symbiont in the whole trabecular syncytium including the dermal layer and the flagellated chambers (see also Supplementary Fig. 4).

Besides a more diverse microbiome than previously reported (34-36), the most visible feature emerging from the assembly process was the presence of 11 super-scaffolds of a Thaumarchaeota-like genome associated with the highest coverage value (Fig. 2a, Supplementary Table 1). This coverage corresponds to a ratio of about 13 archaebacterial cells per sponge cell (presumed diploid according to the few sponge karyotypes available (50)). This Thaumarchaeota species represents the main microorganism associated with *O. minuta*. Its global protein sequence similarity with the NR databases and phylogenetic analyses strongly suggests that it is a new representative of the *Cenarchaeum* genus that we propose to name *Candidatus Cenarchaeum massiliensis* (Fig. 2b, Supplementary Fig. 1). The partially assembled genome sequence of *Ca. C. massiliensis* has a size of about 1.63 Mb, a G+C content of 37.3±2.5%, an estimated completeness of 99.03%(51) and a relatively low coding density (83,9%) compared to other Ammonium-Oxidizing Archaea (AOA) species (Supplementary Table 3). This genome encodes the Thaumarchaeota trademark protein set, including specific ribosomal proteins (presence of r-proteins s26e, s25e and s30e but absence of the r-proteins L14e and L34e), DNA topoisomerase IB and sub-units of ammonia monooxygenase (Fig. 2b, Supplementary Table 4). Among the 908 gene families conserved across the five genera of Thaumarchaeota, 128 are Thaumarchaeota-specific (15%). A large part of Thaumarchaeota core genes (7%) are involved in the metabolism of cofactors and vitamins, including those essential for cobalamin biosynthesis (Supplementary Fig. 2 and 3). Interestingly, as much as 147 transposase encoding genes are present in this species, this number is among the largest reported to date in a thaumarcheotal genome (52, 53).

**Figure 3.**
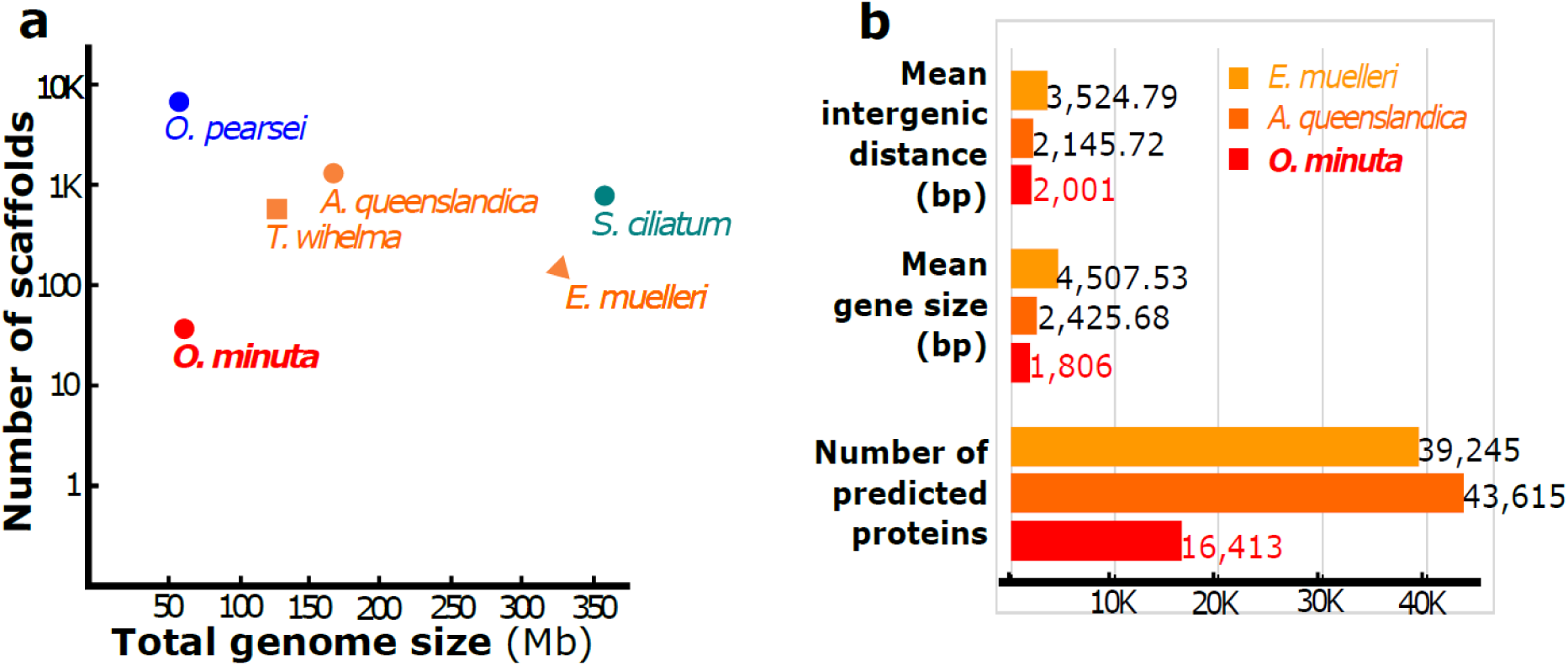
General features of the nuclear genome of *Oopsacas minuta*, compared to that of other sponges. **a:** The size of the assembled genome of *O. minuta* is one of the smallest described so far in sponges. **b:** The *O. minuta* genome has fewer predicted proteins and is more compact compared to demosponges (the sister group to hexactinellids). Other genomic features are available in Supplementary Tables 5,6 and Supplementary Fig. 5.

Using Card-FISH we localized *Ca. C. massiliensis* in the tissues of *O. minuta*. The microorganism is present in the trabecular syncytium and in embryos (Fig. 2c, Supplementary Fig. 4). Similar localization was observed in different individuals sampled at different periods. Taken together, the abundance, the durability of the association and the presence in embryos suggest that *Ca. C. massiliensis* is a symbiont of *O. minuta*, probably inherited by vertical transfer as observed in other sponges (54-56).

Our finding further supports the hypothesis that a stable Thaumarchaeota-Porifera relationships might be based on the ability of all Thaumarchaeota to oxidize ammonia in nitrite and to produce cobalamin anaerobically (52). Such capabilities are absent in animals while cobalamin is essential for their life cycle (57, 58). We propose that the symbiotic *Ca. C. massiliensis* may provide cobalamin to *O. minuta* while recycling the ammonia produced by its metabolism (59).

### *O. minuta* genomic features

The *O. minuta* genome was assembled into 365 scaffolds. Its G+C content is 36±2.1 %. The quality and low fragmentation of the assembly is attested by a N50 value of 0.67 Mb comprising 31 sequences. The estimated total genome size is about 61 Mb, thus 2 to 6 fold smaller compared to Demospongiae and Calcarea (Fig. 3a, Supplementary Table 5a). It is one of the smallest sponge genomes reported so far (*Oscarella pearsei* : 57,7 Mb)(30). Such differences in genome size are consistent with previously reported DNA contents (60).

This small genome size correlates with (Fig. 3b, Supplementary Tables 5): i) A low fragmentation of genes: although the average intron size (341bp) is similar to that of other sponges, the number of introns per gene (1.59 in average) is lower compared to demosponges. ii) A higher gene density: the size of intergenic regions (2 kb in average) is smaller than in *A. queenslandica*. *O. minuta* exhibits the most compact sponge genome assembled so far. iii) A much smaller number of encoded proteins: 16,413 for the *O. minuta* genome, compared to 39,245 for *E. muelleri* (28), 40,122 for *A. queenslandica* (19), and 37,416 for *T. wilhelma* (27). iv) Nevertheless the high number of conserved core eukaryotic genes identified in the final draft sequence attests of its completeness (91 to 93% using BUSCO (61)).

Through a first automated pass of functional annotation, 7,737 protein-coding genes were attributed a Gene Ontology (GO) annotation (Supplementary Table 6, Supplementary Fig. 5).

### The epithelial gene set of a syncytial sponge

The unique syncytial organization of Hexactinellida (39), (74-76), (65),(78) raises exciting questions regarding the genes involved in the epithelial characteristics: cell polarity, cell junctions (CJs) and basement membrane (BM) (18,66–72). A previous survey (32) showed that *Oopsacas* possesses the whole set of genes encoding proteins involved in the polarity complexes (PAR, CRUMBS, SCRIBBLE), and in Cadherin-Catenin complexes (CCC) despite the absence of adherens junctions (AJs). Here we searched for proteins involved in other types of CJs and in the BM.

Three types of cell-cell junctions are defined: adherens, gap and septate junctions (AJs, GJs and SJs) (71,73–77). No innexin-related genes, involved in GJs, were found in this genome which is to be expected as electrical signaling occurs *via* syncytia. The content of the glass sponge proteinaceous plugged junction remains to be determined (36, 78). Of SJ proteins, neither Claudin, neuroglian, neurexin IV nor contactin were found. Regarding contactin (Cont), which belongs to the Immunoglobulin superfamily (IgSF), neither our best blast hit (Supplementary Table 7) nor the sequence from *A. vastus* annotated as contactin (23),(79) exhibit the characteristic glycosylphosphatidylinositol (GPI) anchor domain, supporting the absence of Cont in glass sponges (Fig. 4a). Finally, glass sponges have septae reminiscent of SJs(80) but do not encode the proteins involved in bilaterian SJ suggesting these are convergent structures.

**Figure 4.**
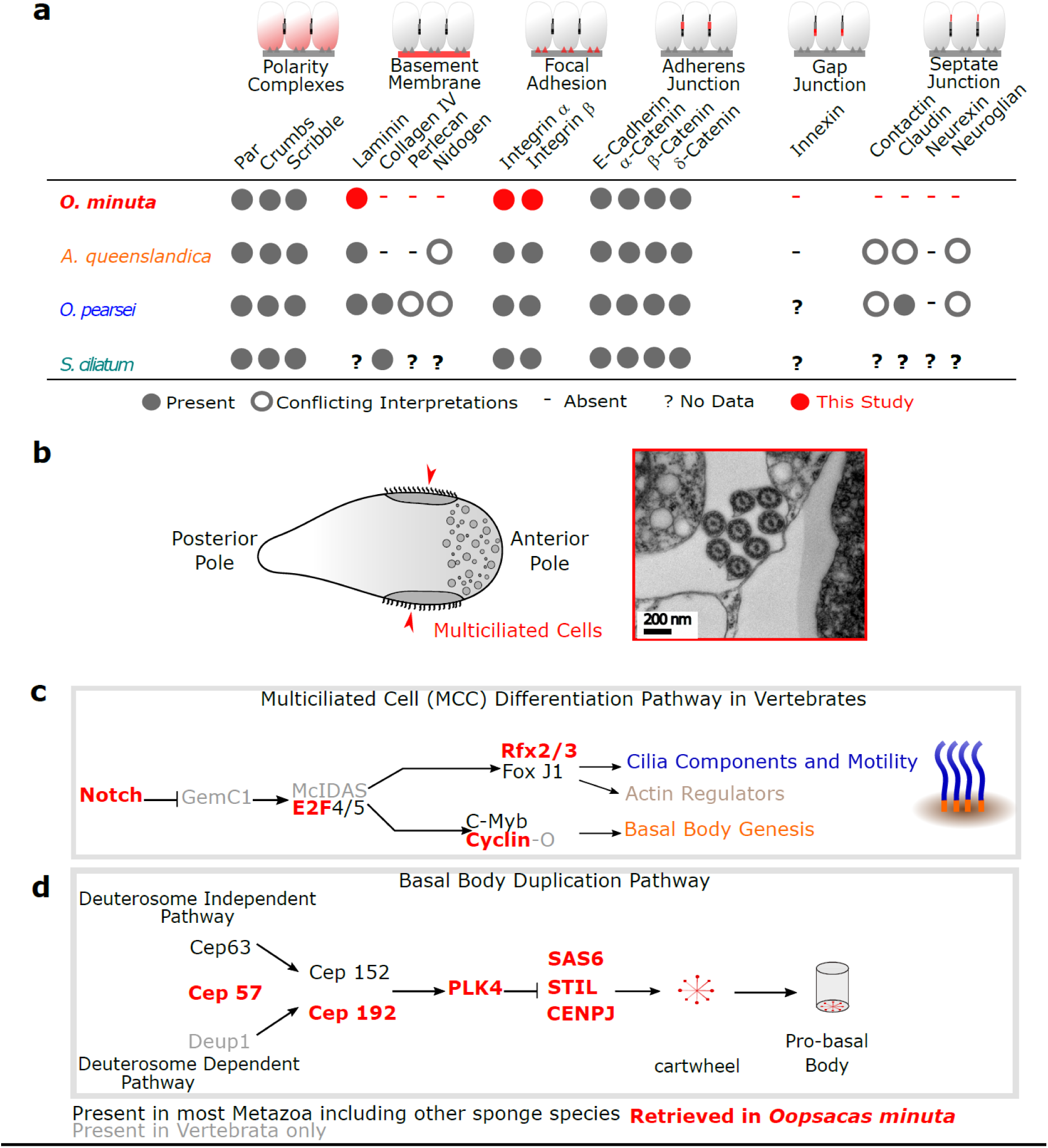
Survey of genes associated with tissue features of *O. minuta*. **a:** Presence/absence in *O. minuta* (syncytial tissue) of genes involved in epithelial functions compared to sponges pertaining to the three other classes (epithelial tissue). **b:** *O. minuta* larvae possess multiciliated cells (MCC), seen clearly in only one other group of deepwater demosponge (cladorhizids). The diagram of a larva (on the left) shows the position of MCC (several cilia are visible in cross section in the TEM picture on the right, photo credits Sally Leys). **c:** Survey of genes involved in MCC differentiation in vertebrates. **d:** Survey of genes involved in ciliogenesis and basal body duplication in bilaterians.

In cell-to-matrix junctions, the bilaterian focal adhesions (FAs) and hemidesmosomes (HDs) are based on interactions between integrins, proteins of the extracellular matrix (ECM) and the actin network (81-84). We found a diversified set of integrins as in other sponges (20,23,28,85,86): Five were assigned to the alpha chain family and three to the beta chain family. Their predicted domain structures are similar to that reported in other animals (Supplementary Fig. 6a).

Among the main components of the BM (laminins, type IV collagen, nidogen and perlecan (87-91)), only four laminins were found (Fig. 4a) with a shared characteristic domain architecture: one α-like laminin-related protein, two chimeric Lamβ/γ-like chains as in *A. queenslandica* (92, 93) and a laminin γ-like chain with a characteristic single LamIVA domain (Supplementary Fig. 6b). Both our and Fahey’s surveys (92) suggest that α, β/γ and γ-like laminin chain types were present in the last common ancestor of metazoans (LCAM). In contrast, a β-chain is identified neither in the demosponge *Amphimedon* nor in Hexactinellida. When reassessing (by domain prediction) the sequence annotated as nidogen in *Aphrocallistes vastus* (23), we predicted only the NIDO domain, which is not specific to Nidogen. The absence of all components of the BM except laminins confirms morphological observations and previous transcriptomic analyses (18,23,67,71) that a BM is absent in glass sponges (68, 69,94). Type IV collagen was probably present in the LCAM and secondarily lost several times (28,79, 87,95,96); in contrast, perlecan and nidogen probably emerged more recently.

### Multiciliogenesis

Multiciliated cells (MCCs) have been described in Vertebrata (97), Spiralia, Ctenophora and Porifera (98-102). Vertebrates and protostomes share the same core set of proteins required for centriole formation and duplication (PLK4, SAS6 and STIL) but not the upstream regulatory network (103-106). Three evolutionary scenarios are possible: 1) the upstream regulators of vertebrate MCCs are ancestral and were lost/replaced in protostomes, 2) the (unknown) upstream regulators of protostomes are ancestral and were lost/replaced in vertebrates, 3) none of the upstream regulators are ancestral and MCCs emerged several times. Data from non-bilaterians is needed to assess these scenarios. *O. minuta* is one of the few sponge species having MCCs in the larva (Fig.4b)(34,65,78,94,107,108). We therefore searched for genes involved in MCC differentiation and centriole duplication in bilaterians. Except for the Notch signaling pathway (next section) neither the vertebrate upstream regulators (Mcidas, GMNC/GEMC1 and E2F4) nor their targets (FoxJ1, c-Myb, Deup 1, Cyclin O) involved in MCC differentiation were found (Fig.4c, Supplementary Table 8). The absence of FoxJ and Myb-related genes is unexpected, because they have an ancient origin (109, 110), and the latter were reported in sponges (111). This suggests secondary losses in hexactinellids.

Among the key upstream proteins involved in bilaterian centriole duplication (Fig. 4d), we found in *Oopsacas* genes encoding CEP192 and CEP57, but not CEP63 and CEP152 (Supplementary Table 8), suggesting that centriole duplication may be controlled by a different regulatory network than in bilaterians. In contrast, we found PLK4, SAS6, STIL and CENPJ, in agreement with their conserved function in centriole duplication across Eukaryota (112).

The above findings suggest that the formation of multiciliated cells in *Oopsacas* may involve common downstream terminal effectors, but that upstream regulators are different from vertebrates and protostomes. In other words, sponge and bilaterian MCCs probably result from convergent evolution, as previously proposed on the basis of ultrastructural (106) and embryological (78) observations.

### Signaling pathways

In metazoans, conserved signaling pathways are critical transduction cascades (113-119). Surprisingly, transcriptomic analyses from glass sponges have suggested that key components of the canonical Wnt pathway are absent (23, 26,120).

Our present analysis of the *Oopsacas* genome confirms that neither *wntless*, *wnt*, nor *dishevelled* genes are present (Fig. 5a). *frizzled A* genes are absent, whereas *frizzled B* genes are present. In contrast, core members of two other key developmental pathways (Notch and TGFβ) are present (Fig. 5b, Supplementary Table 9). So far, the absence of the Wnt pathway was only reported in myxozoans (121), a group of microscopic parasitic cnidarians, whereas glass sponges do not show a highly reduced body plan (Fig. 1c). Interestingly, this finding challenges the pivotal role often attributed to the Wnt pathway in the acquisition of multicellularity and axial patterning (122).

**Figure 5.**
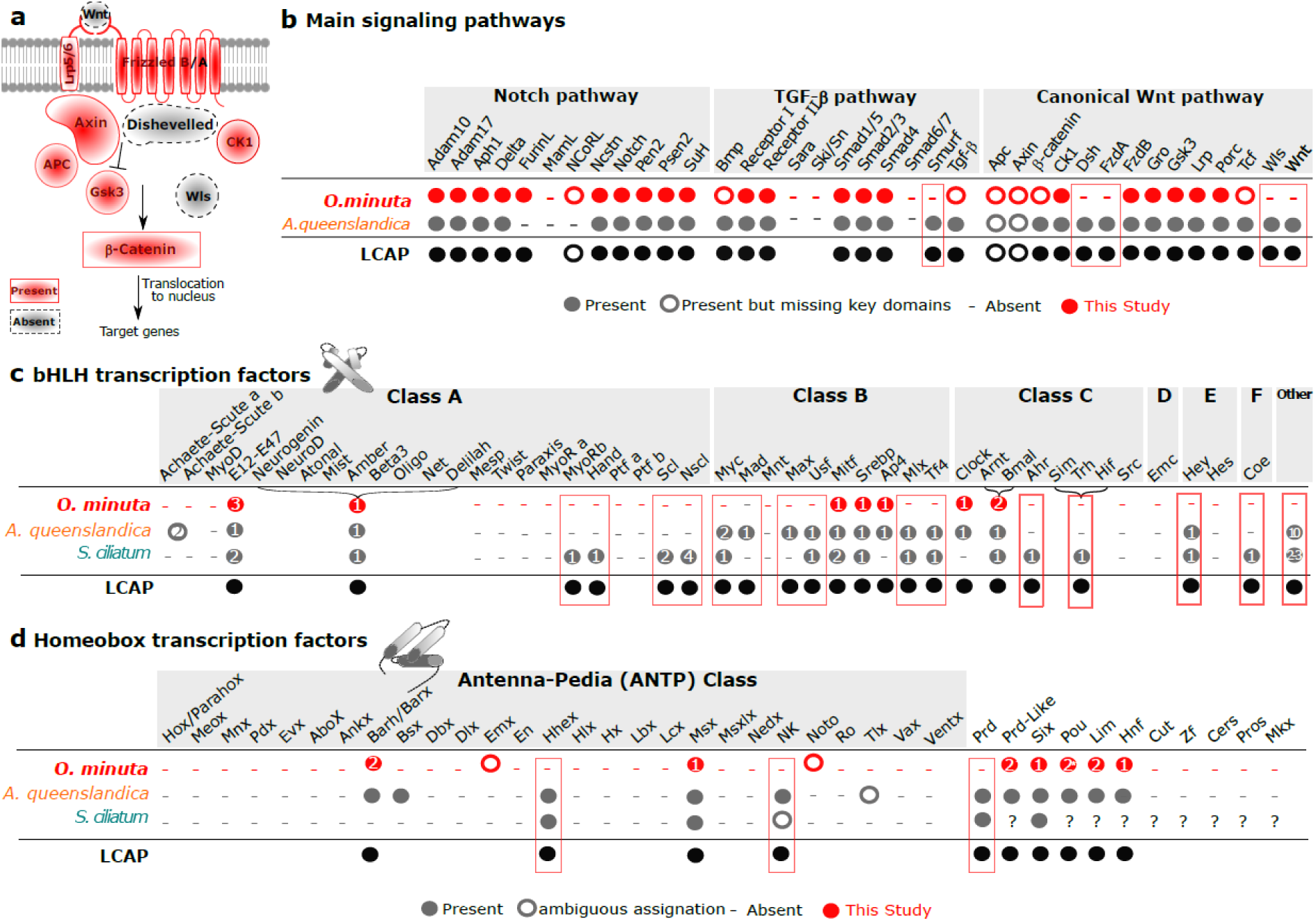
Survey of key genes involved in the metazoan developmental toolkit. **a:** Presence/absence of genes involved in the canonical Wnt pathway: the absence of Wnt and Dishevelled genes is an unusual feature Suggesting alternate pathways to the Wnt pathway. **b:** Survey of genes involved in three critical signaling pathways in *O. minuta* compared to the demosponge *Amphimedon queenslandica* : Notch and TGF-β core members are present allowing to hypothesize functional pathways in contrast to the Wnt pathway. **c:** Inventory of basic Helix loop Helix (bHLH) transcription factors present in *O. minuta* compared to two other sponges. Numerous ancestral genes are absent in *O. minuta*. **d:** Inventory of Homeobox transcription factors present in *O. minuta* compared to two other sponges. Several ancestral classes are absent in this species. Numbers in circles indicate the number of genes found,? indicates the absence of data in the literature.

In search of a pathway that may compensate for such an absence, we surveyed genes encoding heteromeric G proteins and G protein-coupled receptors (GPCRs) known to play important roles in transducing a broad range of extracellular signals. Heterodimeric G proteins have three subunits, Gα (5 classes: Gs, Gi, Gq, G12 and Gv (123)), Gβ and Gγ. Here, we found a diversified set of 8 Gα: one belonging to each of the Gs, G12 and Gv classes, two to the Gi class and three to the Gq class (Supplementary Table 10)(124). We suppose that early development must rely on a truncated Wnt pathway (no Wnt activation) and that the expanded Gq set in Porifera (and Ctenophora), compared to most bilaterians, placozoans and cnidarians (124-127) suggests a broader involvement of Gq proteins in these animals. Because recent studies have suggested the ability of Fzd5 to activate Gq (128), the FzdB copy found in glass sponges (the Fzd5-ortholog) might not be a remnant of a reduced Wnt pathway but might reflect its involvement in G protein signaling.

### Transcription Factors

Transcription factors (TFs) are also pivotal in body patterning. Here, we focused on basic Helix Loop Helix (bHLH) and Homeobox classes because 1) these TFs are significantly enriched in the animal TF repertoire (110), 2) they were exhaustively surveyed in other sponges (31).

There are 6 major groups (A, B, C, D, E, F) of bHLH TFs (129). We identified genes for 10 proteins with a bHLH domain, two of which with an additional Per-Arnt-Sim (PAS) domain (Fig. 5c). Our phylogenetic analysis showed the presence of proteins in the AP4, MITF, SREBP, and E12/E47 families (class B bHLH) (Supplementary Table 11). In addition, one *Oopsacas* protein was found to cluster with several bHLH families constituting the Atonal-related superfamily (130-132). The three remaining proteins are class C bHLH-PAS proteins: one of the Clock family, the two others clustering with ARNT and BMAL families(130). Neither D, E nor F class members were identified (Fig. 5c). There are clearly fewer bHLH proteins in *Oopsacas* than in *S. ciliatum* (30) and *A. queenslandica* (21). More than ten families that are likely ancestral to metazoans have been lost in this sponge.

Among the animal Homeobox TFs classes, the Antennapedia (ANTP) and Pair-Ruled (PRD) are the largest ones (133-135). In *Oopsacas*, we found 20 predicted proteins with a homeodomain (HD). According to both the identity of the best blast hits and domain analyzes (Supplementary Table 12, Supplementary Fig. 7), only 5 sequences pertain to the ANTP class.

Members of Msx (1), Barh (2) families are present while the relationships of the two remaining sequences is unclear (Fig. 5d, Supplementary Fig. 8). There are 6 members of the super-class TALE, 2 members in the prd-like, LIM and Pou classes. We also found an unusual truncated «Pou-like» gene (Supplementary Table 12, Supplementary Fig. 7). We identified one sequence assigned to the SIX and the HNF classes (Fig.5d). We found proteins containing a PAX domain, but none associating HD and PRD domains, suggesting that Prd class members are absent. We did not find any sequence from the ZF, CUT, PROS, CERS and MKX classes.

The number of ANTP TFs in *Oopsacas* (5) is lower than in other sponges (9-11)(31). According to previous studies some ANTP members were lost independently in sponge classes. The absence in *Oopsacas* of Hhex, NK and Prd genes usually considered ancestral is meaningful (31,134,136,137,138,139). In summary, the TF content of *Oopsacas* is different from that of other sponges and exhibits a reduced number of the core metazoan TFs. This reduced TF content may be linked to their lower complexity in terms of cell (uni or multinucleated) types (36).

### Photokinesis and signal transduction

Like unicellular eukaryotes, sponges lack neurons but respond to stimuli(140-148). Photokinesis is the best studied sensory mechanisms involved in sponge larval behavior; however, all sponge genomes examined so far lack opsins, as is also the case here. In contrast, *O. minuta* possesses two photolyase genes as *A. queenslandica* (Supplementary Table 13). Experiments would be needed to document if the involved mechanisms in *Oopsacas* are similar to that described in *Amphimedon* (149, 150).

Hexactinellids are unusual among Porifera in terms of signal transduction because they coordinate arrests of their feeding current using action potentials that travel through syncytial tissues (143, 151,152). Although glass sponge syncytia are quite different than neurons, as they do propagate electrical signals, we searched for genes typically involved in chemical signalling in bilaterians. We found no evidence for conventional monoamine signaling receptors or biosynthesis pathway components (serotonin or dopamine). In contrast, components for glutamate and GABA synthesis and signaling were identified. Nitric oxide signaling is likely present as well as acetylcholinesterase (Supplementary Table 14, Supplementary Fig. 9).

In terms of voltage gated ion channels (Supplementary Table 15), we found genes for voltage gated calcium channels (Two Pore channels, sperm-associated calcium channels and voltage gated Hydrogen channels) (Supplementary Fig. 10) but no genes for voltage gated sodium channels (Nav) nor for obvious voltage gated potassium channels (Kv) (Supplementary Fig. 11). We did not find ENaC (epithelial sodium channels) nor LEAK channels. Ionotropic glutamate receptors (iGluR) were also absent. In contrast, four genes encoding for anoctamins (voltage sensitive calcium activated chloride channels) and six chloride channels (H+/Cl-transporters) were identified. *Oopsacas* also has cyclic gated nucleotide (CGN/HCN) channels as well as purinergic and ryanodine receptors. In addition, there were many hits for Transient Receptor Potential (TRP) family proteins (including one TRP-ML and one TRP-A family protein).

The study of glass sponge conduction system performed on *Rhabdocalyptus dawsoni*(152) suggested that action potential could be driven by calcium because it was blocked by calcium channel blockers. This may be consistent with the diversity of calcium channels identified here. While the return to resting potential is sensitive to a potassium channel blocker, *Oopsacas* does not appear to possess the voltage sensor region of K channels, so it is not known which channels are responsible for resetting the membrane potential.

Finally, we also surveyed genes involved in synapses. Postsynaptic proteins include a range of scaffolding and vesicle fusion/transport proteins, and many are present in *Oopsacas*, among them synaptobrevin, syntaxin, Homer and members of the SNARE family SNAP 25, with the exception of neurexin and neuroligin. In contrast, most of conventional presynaptic proteins such as profilin, synaphin, synaptoporin and synaptogamin were missing (Supplementary Tables 16 and 17). In summary, a complete synaptic machinery is absent in *O. minuta*. Though proteins involved in vesicle transport and ion channels are identified, they may play a variety of other roles.

### The biosilicification toolkit

The typical 6-rayed spicules of glass sponges, hexactines, are made of silica. Although the spiculogenesis has similarities in Silicispongia (Fig. 1a), there are fundamental differences (153-157) between hexactinellids and demosponges. We therefore searched in the genome of *O. minuta* homologs of genes known to be involved in spicule formation in demosponges and hexactinellids (Fig. 6a, 6b, Supplementary Table 18). We found no evidence of *silicatein*, *silintaphin* or *galectin* genes, which are common in demosponges, but a single *glassin* gene, thirteen *cathepsin L* genes (Figure 6c, Supplementary Table 19), one *ferretin*, four *silicases*, and one *chitin synthase* were identified. Shimizu and collaborators (2018)(155) tried unsuccessfully to isolate silicatein proteins from the spicules of the glass sponge *Euplectella aspergillum*, and failed to recover any sequence belonging to the silicatein family in this species. Similarly, we did not find any *silicatein* ortholog (except cathepsin L). Also, transcripts for silicatein were not recovered in the hexactinellid *Aphrocallistes vastus*, or any transcriptome of hexactinellids published to date, raising serious doubts about the existence of silicatein outside demosponges (23, 158). The silicateins purified directly from the skeleton of hexactinellids might therefore have been contaminations (159-163). The most plausible explanation is that hexactinellids and demosponges use different enzymes to build their skeleton, which is also supported by the fact that Homoscleromorpha (the third class with siliceous spicules) lack both silicatein and glassin (158). This has profound implications since it suggests a convergent evolution of sponge spicules.

**Figure 6:**
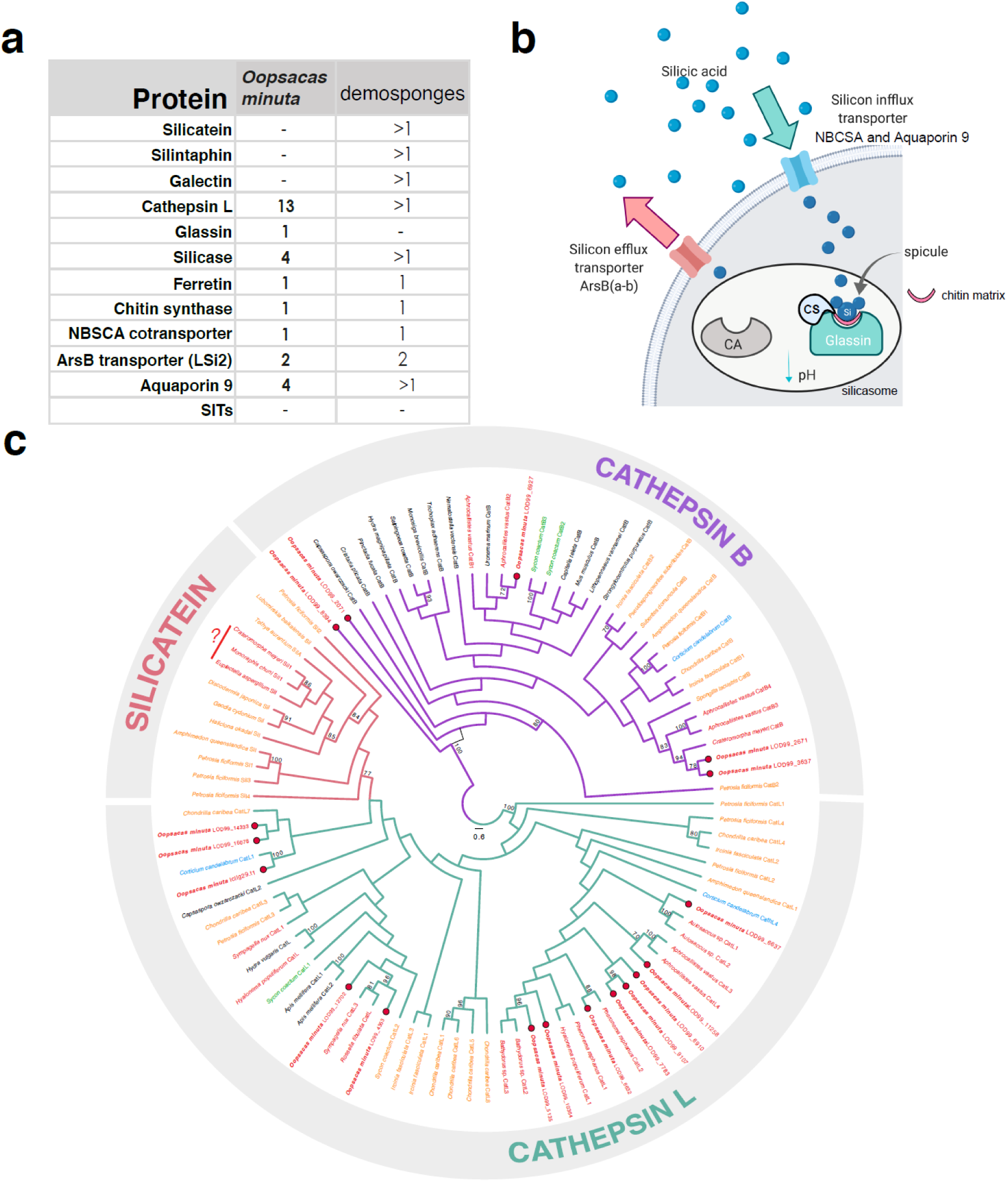
Biosilicification toolkits in *O. minuta*. **a:** Summary of major biosilicification proteins involved in demosponge siliceous skeleton production. **b:** Schematic view of the biomineralization process in the sclerocytes of Hexactinellida. Abbreviations: CA, carbonic anhydrase; CS, chitin synthase; NBSCA, NBC (Na^+^/HCO3^−^ [Si(OH)4]) transporter; Si, biosilica. **c:** Phylogenetic hypothesis obtained by maximum likelihood of cathepsin evolution of the major sponge lineages. The sequences obtained from the genome of *O. minuta* are shown in blue. Bootstrap values less than 70 are not shown. Accession numbers and sequence alignments are available in Supplementary Tables 18 and 19.

For the silicon transport into the sponge cells (164, 165), the *O. minuta* genome encodes a NBCSA cotransporter, four *aquaporin 9* genes, and two arsB transporters. The silicon transporters (SITs) used by other silicifier organisms (166) were not found. The exact mechanisms of interaction between the different enzymes are still far from being understood, but the identification of the largest complement for biosilicification in *Oopsacas* is definitely a fundamental step towards the full characterization of the process in Hexactinellida.

## DISCUSSION

The genome of the glass sponge *O. minuta* differs from all other available sponge genomes by its compactness and smaller number of predicted proteins. It is among the smallest non parasitic metazoan genomes reported so far (121,167–169).

The most striking and unexpected feature is the absence of numerous genes usually considered as ancestral and pivotal for metazoan morphogenetic processes (Supplementary Table 20). Because these genes are present in the three other sponge lineages, these absences most likely represent secondary losses. Importantly, these losses are not associated with a highly reduced body size and/or complexity or a parasitic lifestyle but may reflect the peculiar syncytial organization typical of this group. However, given the large differences documented between demosponge species (28), it would be premature to infer that these characteristics might be shared by the other 696 hexactinellid species (13) at this point. The ongoing sequencing of two other glass sponges *Aphrocallistes vastus* and the “tulip sponge” (Sally Leys and Darrin Schultz, personal communications) are expected to soon allow highly valuable comparisons.

We also report for the first time the association of a new *Thaumarchaeota* species with a hexactinellid sponge, previous reports only mention associations with Demospongiae (52). The unusual enrichment of transposases in its genome is worth noting because it may suggest a particularly high potential for lateral gene transfer (52).

Interestingly, our work further documents the important role of convergence during animal evolution, by suggesting the independent emergences of various mineral skeleton among sponges, of electrical signaling, and of multiciliated cells at an early stage of animal evolution.

## CONCLUSION

Porifera represent a significant part of diversity in aquatic ecosystems first because of their number of species (nearly 9500) but also by the shelter they offer to other ones. They emerged and diversified more than 600 Myrs ago, resulting into four distinct lineages, each harboring particular features. Here we reported the first whole genome analysis of a glass sponge (Hexactinellida), providing new insights on sponge biology and diversity. Despite a general body plan very similar to that of other sponges, Hexactinellida exhibit unexpected genome features and a reduced gene content. More generally because sponges are part of one of the most ancient still extant animal lineages, the study of their genomes is precious to reconstitute the evolution of animals and of their body features. Our finding reinforces the idea that gene losses, convergent evolution of similar body features and tight relationships with microorganisms are processes that crucially contributed to the diversification of animals.

## METHODS

### Sample collections

Adult individuals containing embryos and prelarvae were collected in the 3PP cave, La Ciotat (France, 43° 9.797’N, 5° 36.000’E). Samples were brought back to the laboratory in a cooler, then immediately cut and cleaned of superficial sediment and organisms under a stereomicroscope with brush and forceps and rinsed in 0.2 µm filtered sea water to limit contamination risks. Samples were either used freshly or stored at -80°C for future use.

### Genomic DNA and RNA extraction and sequencing

For Illumina sequencing, gDNA extractions were performed using the QIAamp® DNA Mini kit from QIAGEN according to manufacturer instructions using 22 mg of sample. DNA quality and quantity were checked by electrophoresis, NanoDrop and Qubit fluorometer. RNA extractions were performed (Qiagen kits) by the ProfileXpert IBISA platform. Sequencing was performed using Illumina technology with RNA-seq pair-ends, DNA-seq paired-ends and Nextera Mate Pair protocols on a HiSeq2500 sequencer with a read length of 150 bp (ProfileXpert-LCMT, Lyon, France, http://profilexpert.fr).

For PacBio sequencing, tissues (1.34 g) were first dissociated according to the protocol described for *Oscarella lobularis*(170) but with calcium-magnesium free sea water complemented with 20mM EDTA (instead of 10mM). To remove the spicules the suspension was passed through a 40 µm cell strainer. The cell suspension was concentrated by centrifugation (400g-3min). The DNA extraction was performed on the cell suspension with the MasterPure^TM^ Complete DNA kit according to the manufacturer instructions. DNA quantity and quality were checked by gel electrophoresis, NanoDrop and Qubit. Long read sequencing was performed on a Pacific Biosciences platform (University of Lausanne, Swiss, https://wp.unil.ch/gtf/).

### Genome Assembly

#### -Raw assembly

Genome sequencing was first performed using the Illumina platform as previously described(38). In addition, complementary sequencing was performed on a PacBio platform for a total of 751,460 long reads. These reads were filtered based on their length and quality with Pacific Bioscience tools (SMRT Portal) then self-corrected with Canu(171). All Illumina reads were mapped on the corrected PacBio reads with Bowtie 2(172). Selected Illumina reads and corrected PacBio reads where then assembled together with spades using options careful and Kmers 21,41,61,81 and 99(173).

#### -Scaffolds selection

Only scaffolds longer than 1kb were retained and submitted to MetaGeneMark(174) followed by a blastp(175) strategy against NR (best hits with evalue < 10^-5^) and a LCA-like method computed over all annotated genes for each scaffold. Scaffolds were then attributed to 4 groups: Eukaryota, Bacteria, Archaea and Not Annotated.

#### -Scaffolds polishing

Along with Illumina reads, the Eukaryota group (936 scaffolds) and the Archaea group (50 scaffolds) were individually submitted to SSPACE (176) with parameters -x 0 -z 0 -k 5 -g 0 -a 0.7 -n 15 -p 0. All super-scaffolds where then submitted to pilon (177) with default parameters and Gapfiller (178) with parameters -m 30 -o 2 -r 0.7 -n 10 -d 50 -t 10 -g 0 -i 1 to obtain 11 Archaea and 365 Eukaryota polished super-scaffolds regarded as the two draft genomes.

### Assembly metrics and comparisons

The 11 Archaea scaffolds were submitted to Checkm(51) and presented a completeness of 99,03% without contamination.

To compare the genome quality of *O. minuta* to that of other sponges, the 16,413 genes of *O. minuta* CDS were submitted to BUSCO(61) through the gVolante server (https://gvolante.riken.jp/analysis.html) using the 303 Eukaryota core genes set. The absence of potential contamination by human DNA was checked by blast against NR, yielding no hit with similarities higher than 64%.

### Gene prediction and annotation

#### -Thaumarchaeota

We predicted 1990 archaea genes using GeneMarkS and the --prok option. Each sequence was submitted to multiple annotation strategies. A blastp against NR with an e-value < 10^-5^ and using the 10 best hits was used as the main information for functional annotation. A domain search was performed against Pfam 28.0, TIGRFAM 15.0, SMART 6.2, ProDom 2006.1, PANTHER 9.0, Prosite 20.113, Hamap 201502.04 using interproscan 5.14-53(179). A CD-search(180) was done against the conserved domain database at NCBI. Potential (trans)membrane proteins were predicted using Phobius(181). Specific repeat domains were assessed using hmmsearch(182) on different hmm profiles (ankyrin repeat, BTB/POZ domain, CASC3/Barentsz eIF4AIII binding, collagen triple helix repeat, DUF3420, DUF3447, F-box, MORN repeat, pentapeptide repeat). All predicted proteins shorter than 100 amino acids without match with at least one of the above methods were discarded. These results were manually integrated to improve the functional annotation of the 1675 remaining Thaumarchaeota proteins. All 20 standard transfer RNA except for tryptophan were predicted using tRNA-scan-SE(183, 184). Each 23S, 16S and 5S ribosomal RNAs were predicted using barrnap (https://github.com/tseemann/barrnap) for the archaeal kingdom.

#### -Oopsacas minuta

Transcriptomic reads were mapped to all eukaryota scaffolds using TopHat 2(185) and the genes predicted using Braker(186). All alternative transcripts not supported by Augustus(187) were removed for a total of 16,413 remaining genes. All proteins were annotated on the basis of the best hit using blastp against NR (evalue < 10^-5^). To infer Gene Ontology for each protein, a domain search was performed using InterProScan on the same databases as for *Thaumarchaeota* and the GO identifiers were used on WEGO (http://wego.genomics.org.cn/). Results about the first two levels are presented in Supplementary Fig. S5 for the 7,737 genes that receive a GO annotation (Supplementary Table S6).

All 20 standard tRNAs as well as initiator methionyl-tRNA, Selenocystein tRNA and supressor tRNA were predicted with at least one copy using tRNA-scan-SE for the eukaryotic model. 28S, 18S, 5.8S and 5S ribosomal RNAs were identified using barrnap for the eukaryotes.

More detailed annotations were performed for a number of candidate genes (see result subsections). For this purpose, proteins of interest were specifically searched for using blastP against the predicted proteome of *O. minuta*. The returned hit-blast with evalue < 10^-2^ were then checked by a reciprocal best-hit approach against NR database (NCBI). When needed, additional analyses were performed (phylogenetic analyzes and/or protein domain analyses) on candidates with evalue < 10^-2^, the list of query sequences and of best-hits obtained at the reciprocal best-hit step are provided in Supplementary Tables (7 to 18). Supporting phylogenetic analyses or domain prediction are provided in Supplementary Fig. 8 to 11.

The Whole Genome Shotgun project of *Oopsacas minuta* has been deposited at DDBJ/ENA/GenBank under the accession JAKMXF000000000.

The version described in this paper is version JAKMXF010000000.

This Whole Genome Shotgun project of the *Thaumarchaeota* species *Candidatus Cenarchaeum massiliensis* has been deposited at DDBJ/ENA/GenBank under the accession JAJIZT000000000.

The version described in this paper is version JAJIZT010000000.

### Card-FISH localization of the main Thaumarchaeota species associated to O. minuta

*O. minuta* adults were sampled in the 3PP cave in December 2014 (N=3) and in August 2015 (N=2) and preserved in PFA 4%. After dehydration using ethanol, samples were included in paraffin wax. To obtain 8 µm sections, it was necessary to get rid of abundant silicious spicules for cutting: 5% hydrofluoric acid was applied directly to paraffin blocks for 8 minutes at room temperature; the block was rinsed with distilled water before sectioning. After dewaxing with Neoclear®, endogenenous peroxydase was inhibited by adding 0.3% hydrogen peroxide in the first, of 2, ethanol baths for 15 min, then sections were rehydrated and permeabilized with K proteinase 2µg ml^-1^ for 45 min. Sections were post-fixed with 4% PFA for 15 min and card-FISH was carried out following protocols from(188, 189) (detailed protocols provided upon request).

The most abundant thaumarcheotal species was localized using a specifically designed probe: THAUMOOPS840: CATTAGTACCGCTTCAGACC-HRP.

Negative controls consisted of the absence of probe (with or without TSA) or the use of a random sequence not matching at 100% with any sequence of the metagenome: NONOOPS02: GGTTCCTTAGTCACGCAGAA-HRP. We observed that sponge cells have an endogenous peroxidase activity (green background) incompletely abolished by 0.3% H2O2. Unfortunately, higher concentrations of hydrogen peroxide damaged the tissues to a point precluding localization.

## DECLARATIONS

### Ethics approval and consent to participate

Invertebrates used in this study are not concerned by experimental agreement

### Consent for publication

Not aplicable

### Availability of data and materials

The datasets generated and/or analysed during the current study are available at DDBJ/ENA/GenBank under the accessions JAKMXF000000000 and JAJIZT000000000.

All data generated or analysed during this study are included in this published article and its supplementary information files.

### Authors’ contributions

ALB, CB, ER, JMC designed the project

JMC, SS, CJ chose sequencing and assembling strategy

AR, CB, CJ, CR, ER, HB, MD, MV, MS, QS, SL, SS performed experiments and/or sequence analyses ALB, AR, CB, CJ, ER, JMC, LK, MV, QS, SL, SS interpreted the results, wrote and edited the text

## Acknowledgements

The authors are grateful to the diving staff of the OSU Pytheas (F. Zuberer, L. Vanbostal, D. Guillemain) and to the divers of the IMBE (P. Chevaldonné, T. Perez, C. Marschal, S. Chenesseau, J. Vacelet) for collecting samples. The molecular biology staff of the IMBE is acknowledged for providing reagents and equipment for high quality DNA extraction. We thank the ProfileXpert (Lyon) and Pacific biosciences platforms for genome sequencing. We thank PACA Bioinfo for the computing support during assembly and analysis. We also thanks Marc Garel (MIO, Marseille, France) for kind advice on card-FISH experiments, IBDM imaging and IMBE morphology services for providing technical facilities and help. We also thank A. Ereskovsky and J. Vacelet for helpful discussions on sponge anatomy and embryological features.

## Supplementary Files Index

### Characterization of the microbiome of *Oopsacas minuta*

**Supplementary Table 1:**
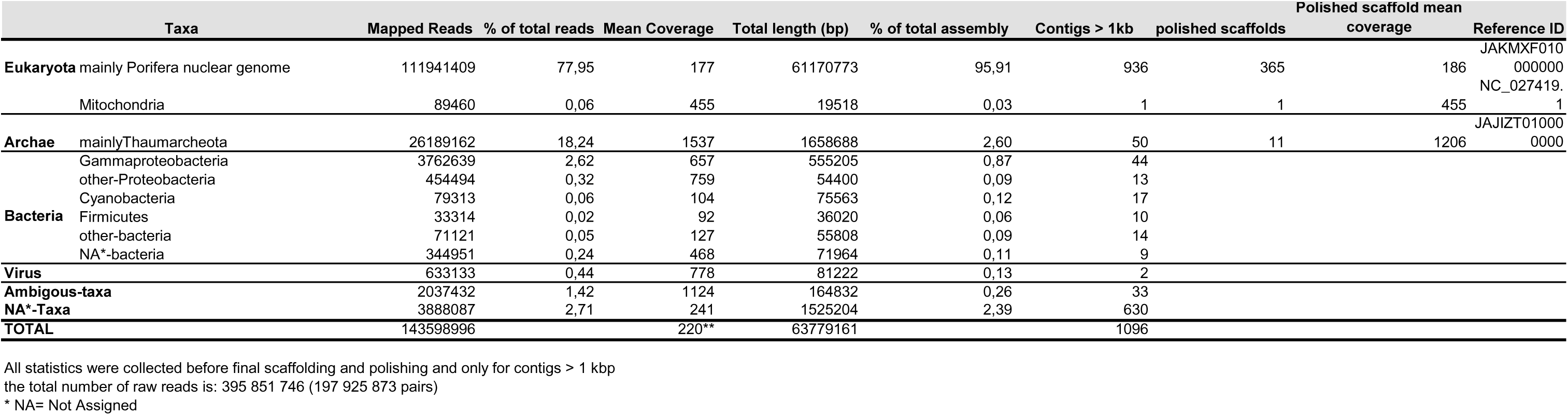
Main features and taxonomic assignation of *Oopsacas minuta*’s metagenome assembly.

**Supplementary Table 2:**
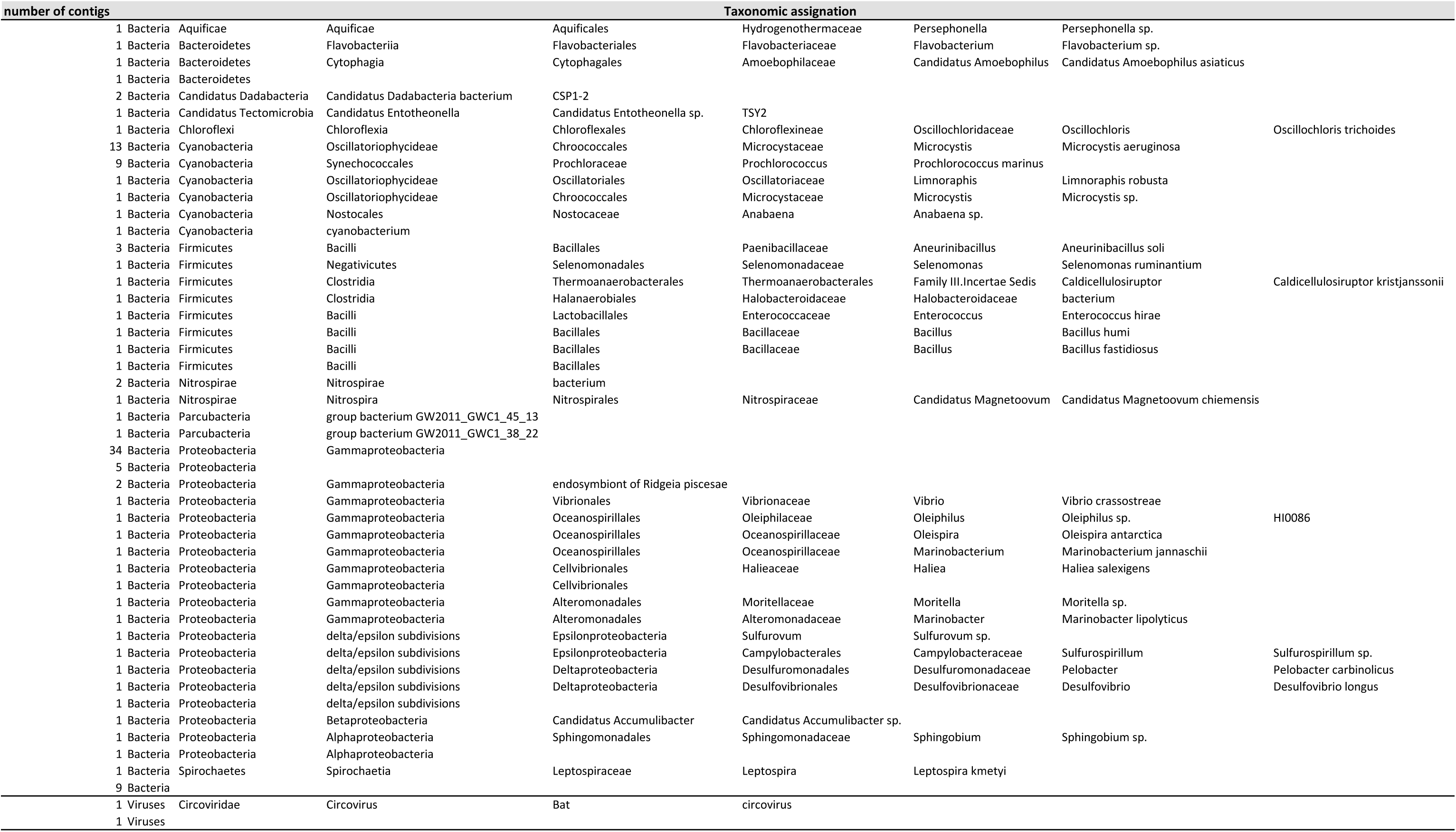
Taxonomic assignation of non-Eukaryotic and non-Archaeal Taxa.

**Supplementary Figure 1:**
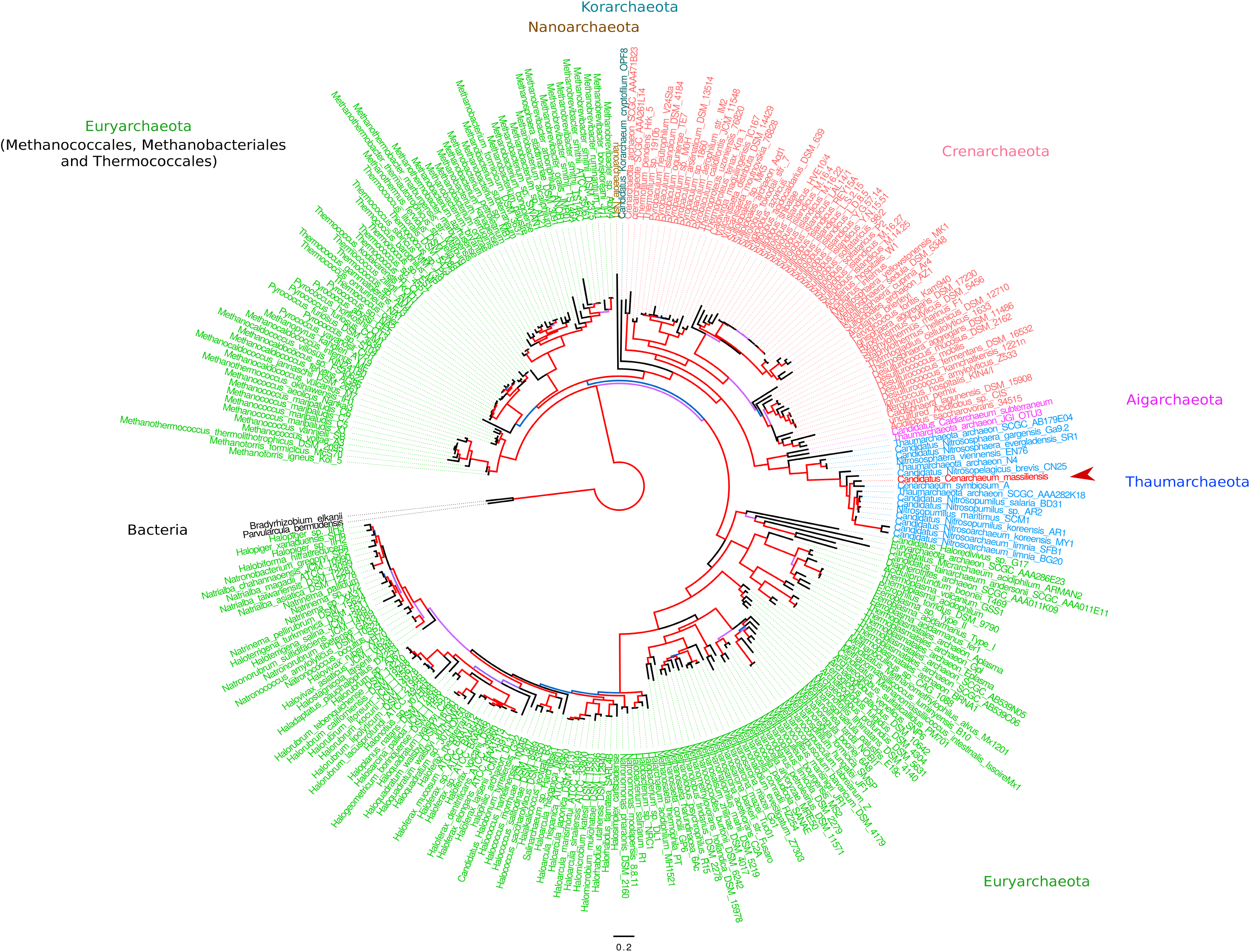
**Phylogenetic position of the main microbial species (in red) associated to *O. minuta*** (infered by maximum-likelihood on ribosomal RNA genes, see material and method section for details). Main archaeal phyla are indicated and branch supports (approximate likelihood ratio teststatistics with a Shimodaira–Hasegawa-like procedure) are indicated with a color code (>0.9, red; >0.7, purple; >0.5, blue and <0.5, black). Scale bar represents the number of substitutions per site.

**Supplementary Table 3:** Genome features of *Candidatus Cenarchaeum massiliensis* compared to other Thaumarchaeota genomes.

| Species | Size (Mb) | GC content | CDS content | Coding genes | tRNA number |
| --- | --- | --- | --- | --- | --- |
| ***Cenarchaeum massiliensis*** | 1.63 | 37.3% | 83.9% | 1675 | 48 |
| *Cenarchaeum symbiosum* | 2.05 | 57.7% | 91.4% | 2017 | 45 |
| *Nitrosopumilus maritimus* | 1.65 | 34.2% | 90.3% | 1795 | 44 |
| *Nitrosopumilus koreensis* | 1.64 | 34.2% | 87.2% | 1890 | 41 |
| *Nitrosoarchaeum koreensis* | 1.61 | 32.7% | 89.0% | 1945 | 42 |
| *Nitrosoarchaeum limnia* | 1.77 | 32.5% | 83.4% | 2038 | 45 |
| *Nitrosopelagicus brevis* | 1.23 | 33.2% | 94.6% | 1445 | 42 |
| *Nitrososphaera viennensis* | 2.53 | 52.7% | 87.0% | 3123 | 39 |
| *Nitrososphaera gargensis* | 2.83 | 48.3% | 81.6% | 3565 | 34 |
| *Nitrososphaera evergladensis* | 2.95 | 50.1% | 83.4% | 3499 | 39 |

Mb, megabase; CDS, coding sequence.

**Supplementary Table 4:**
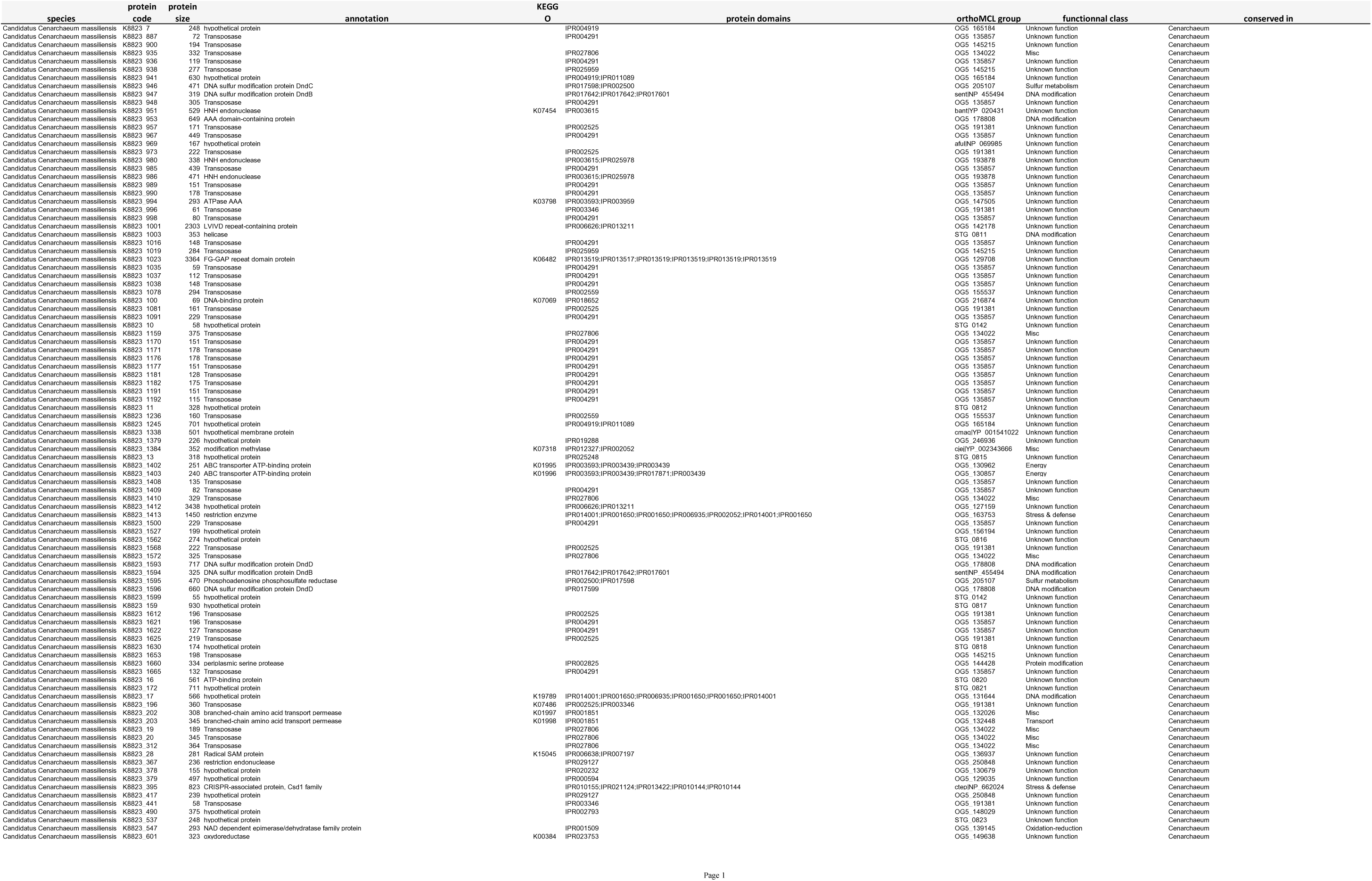
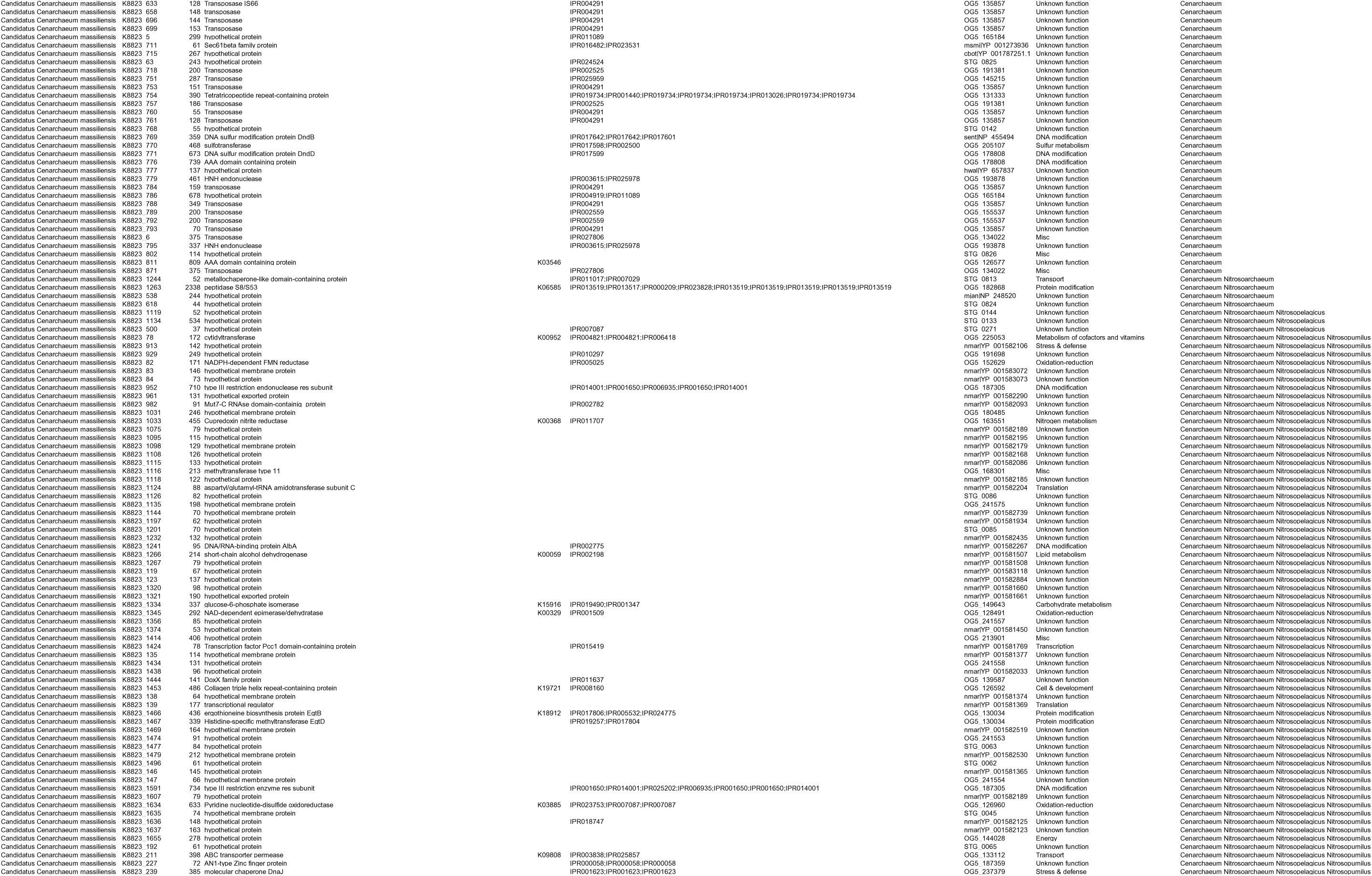
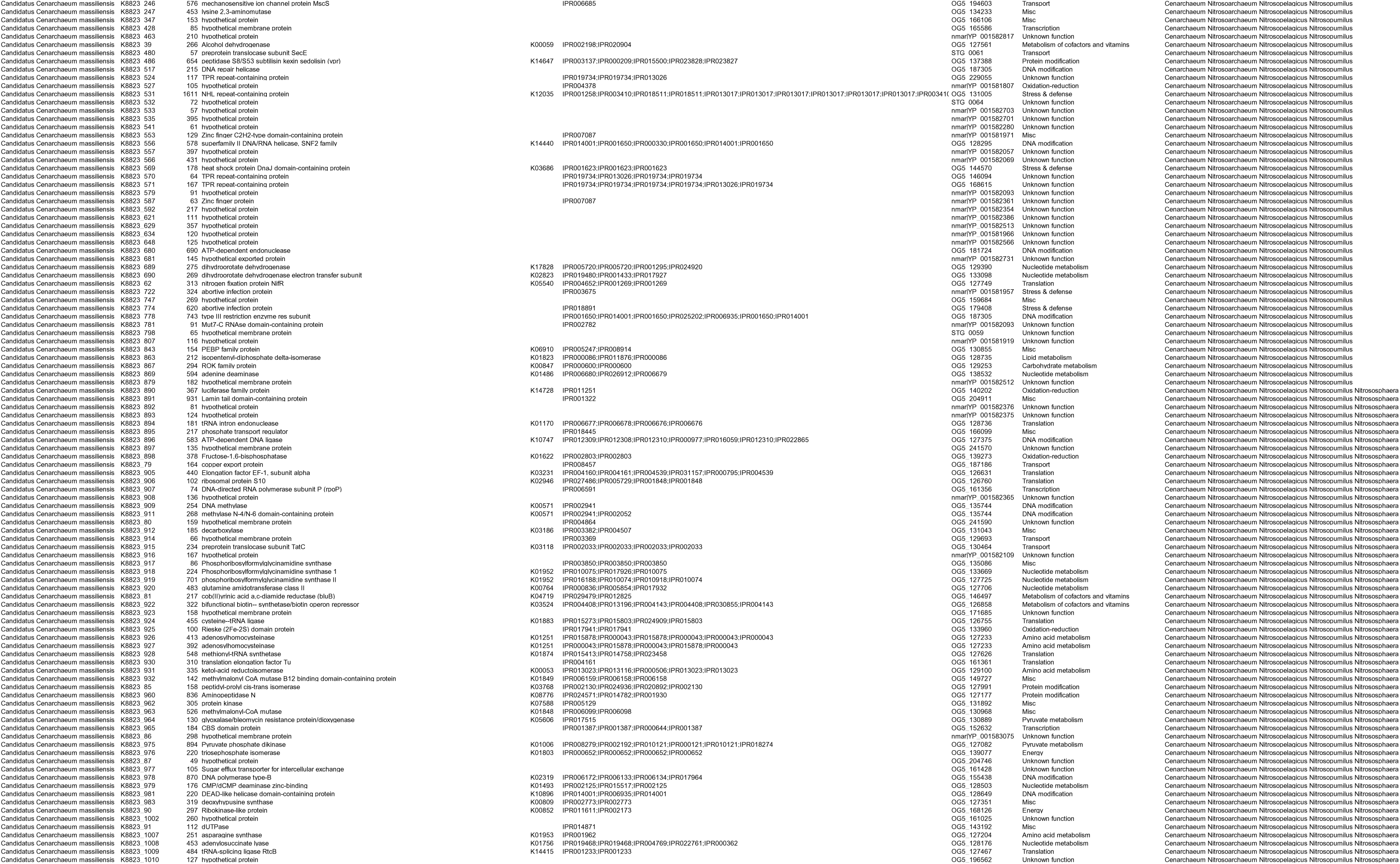
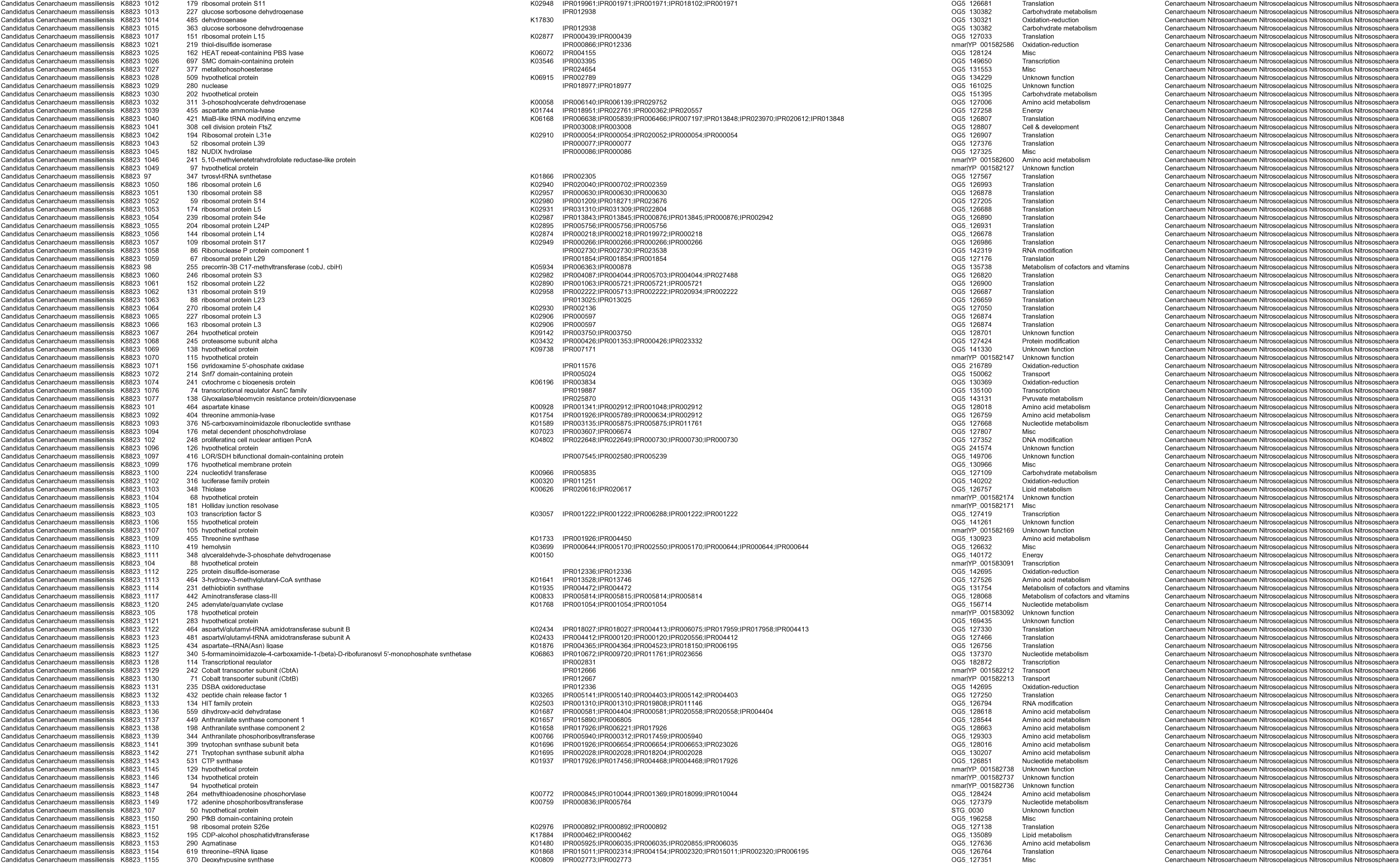
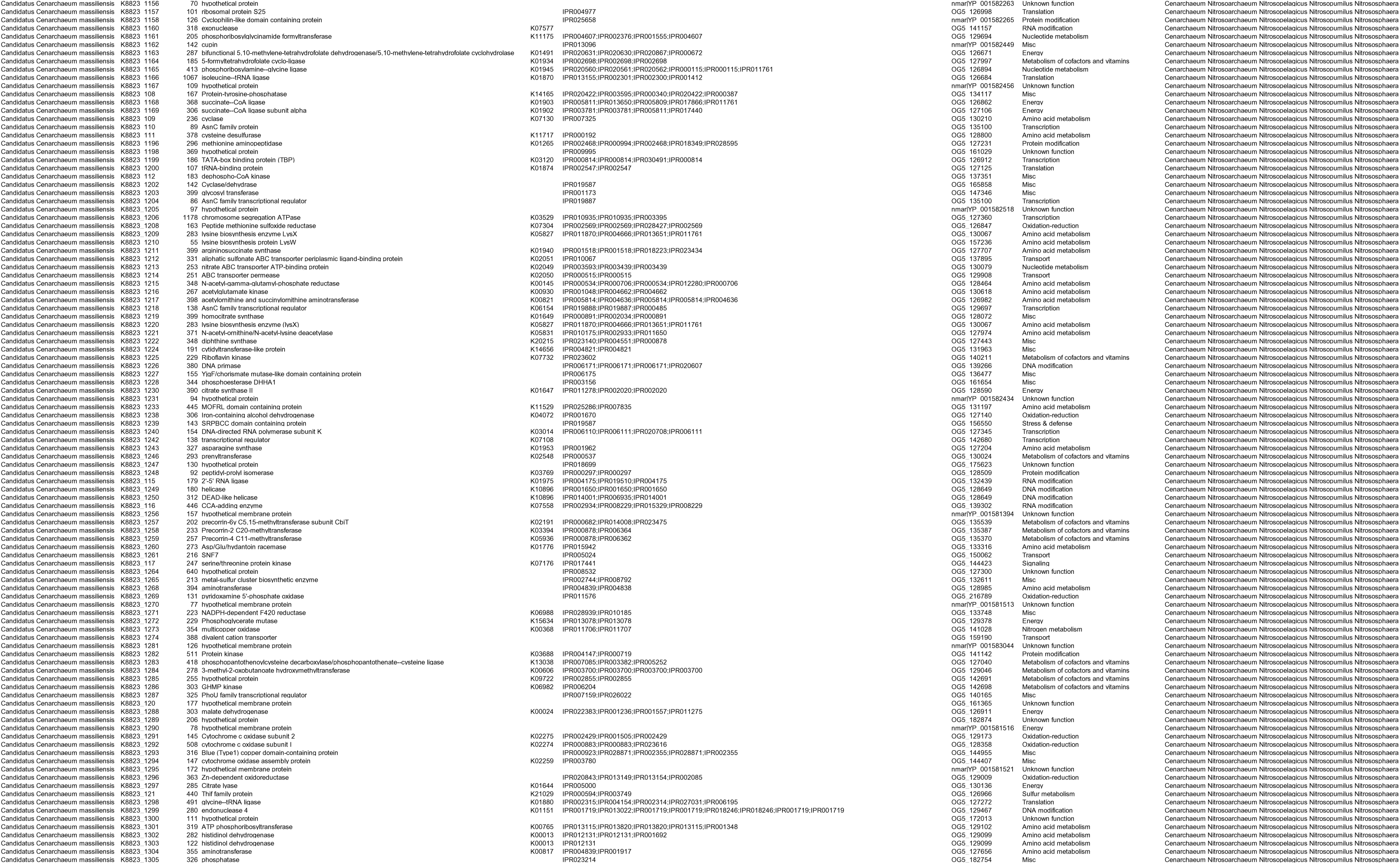
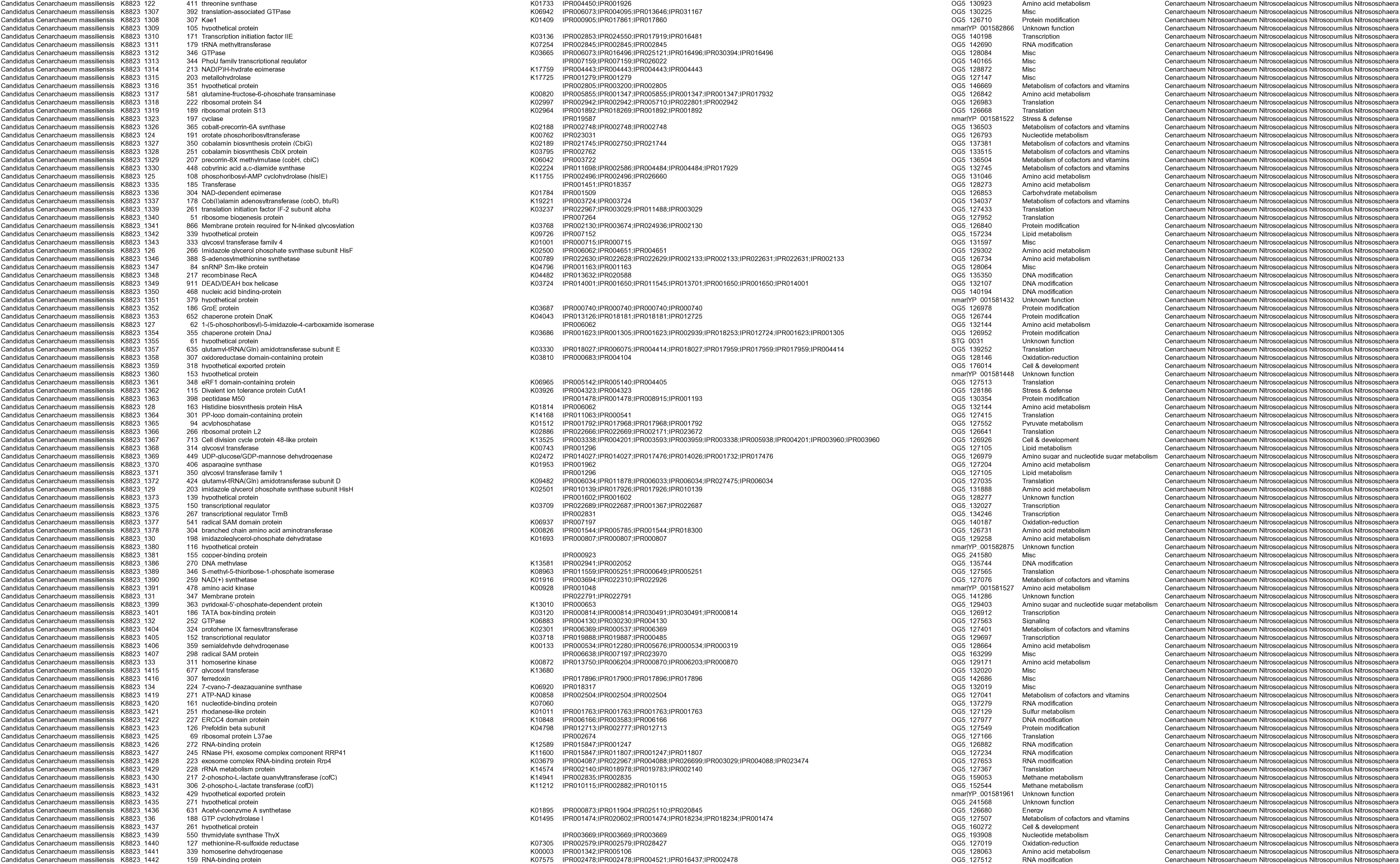
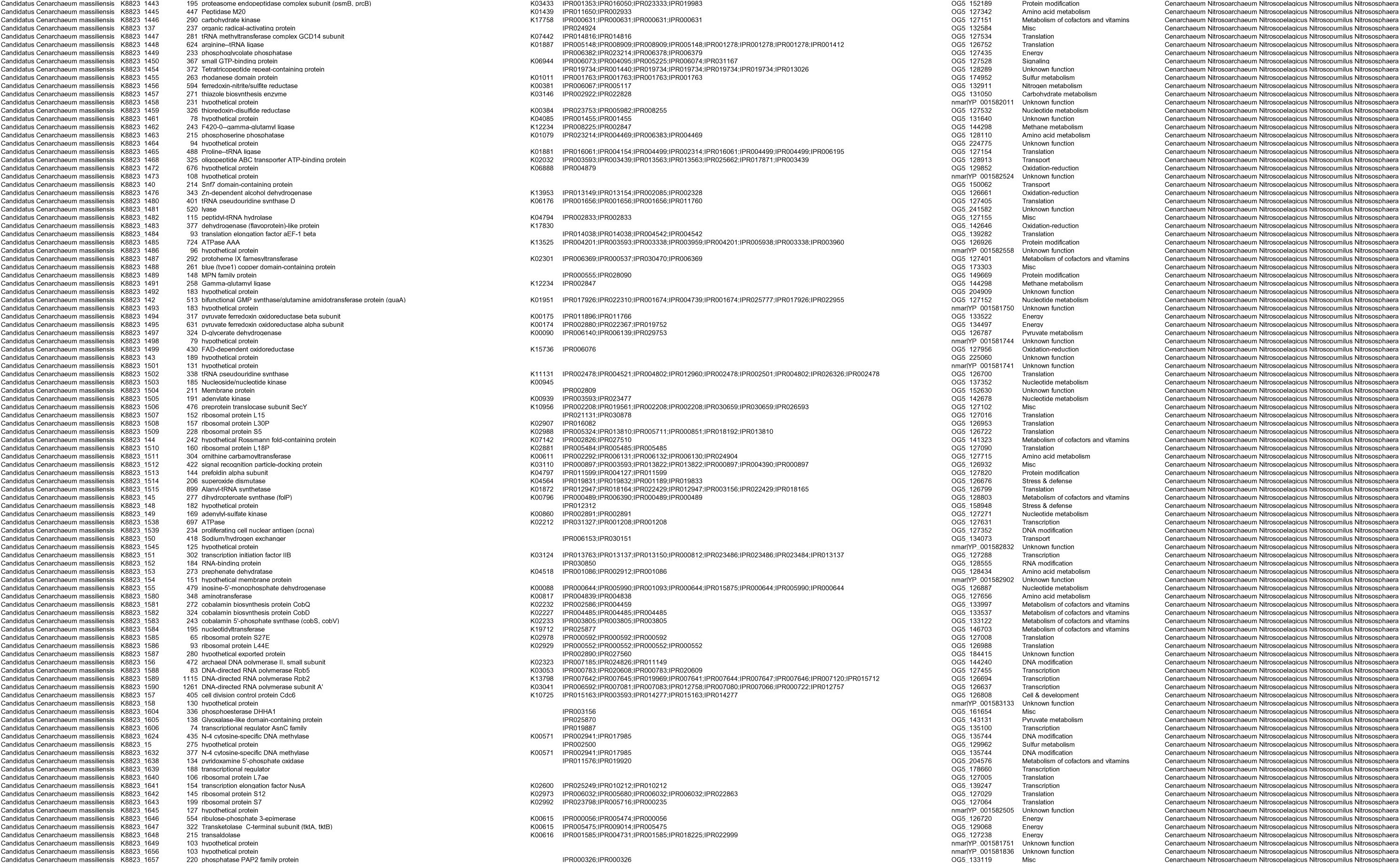
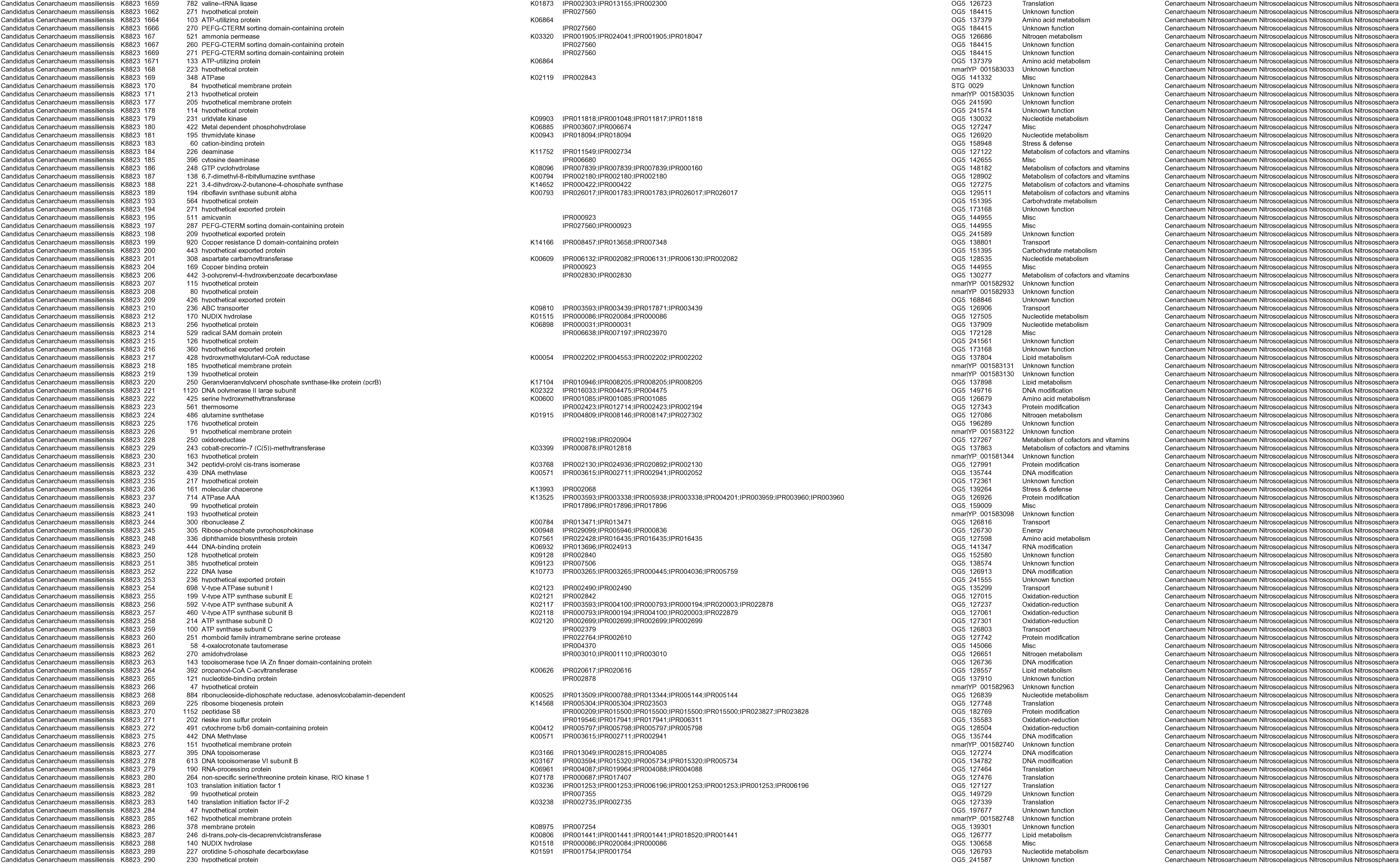
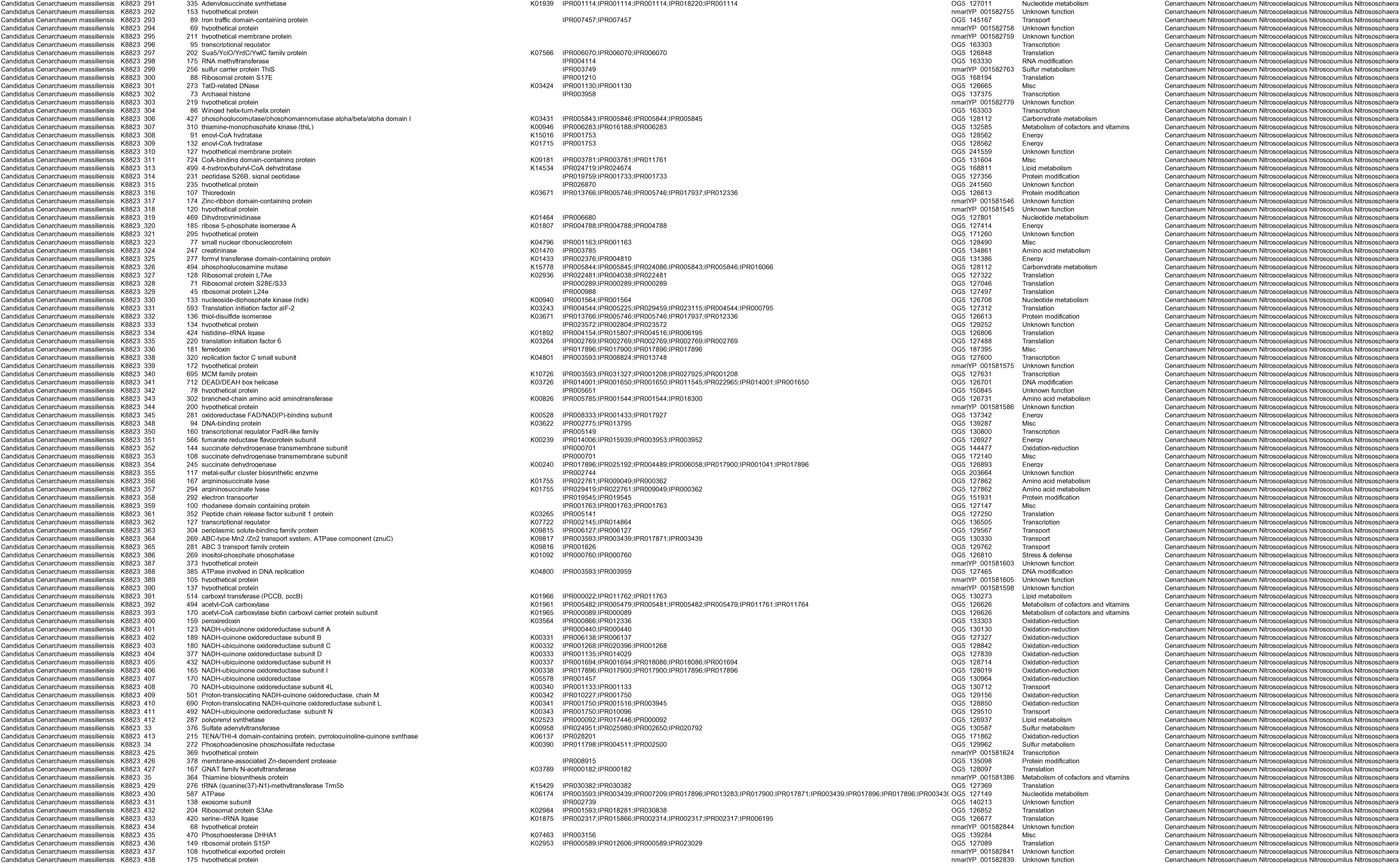
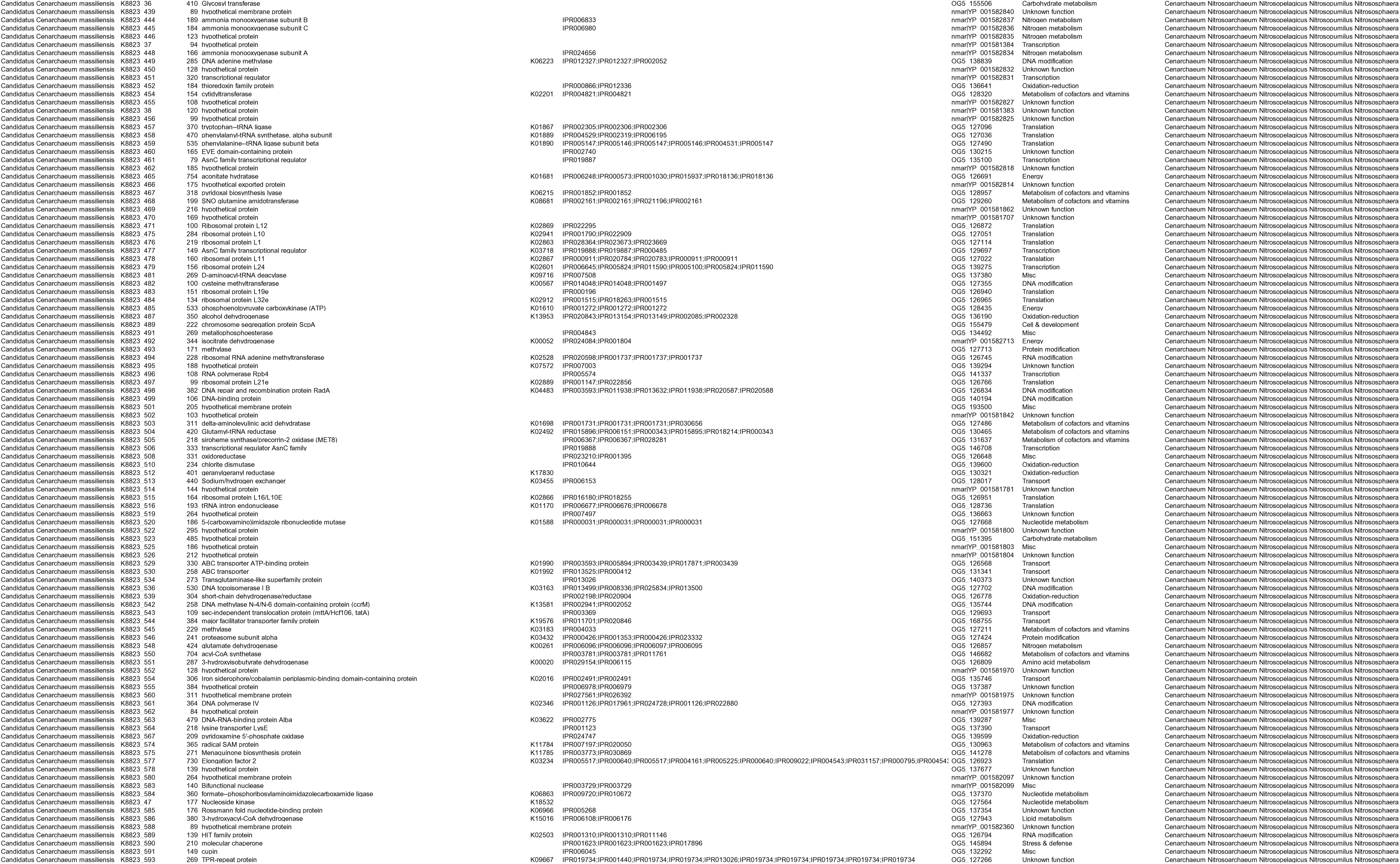
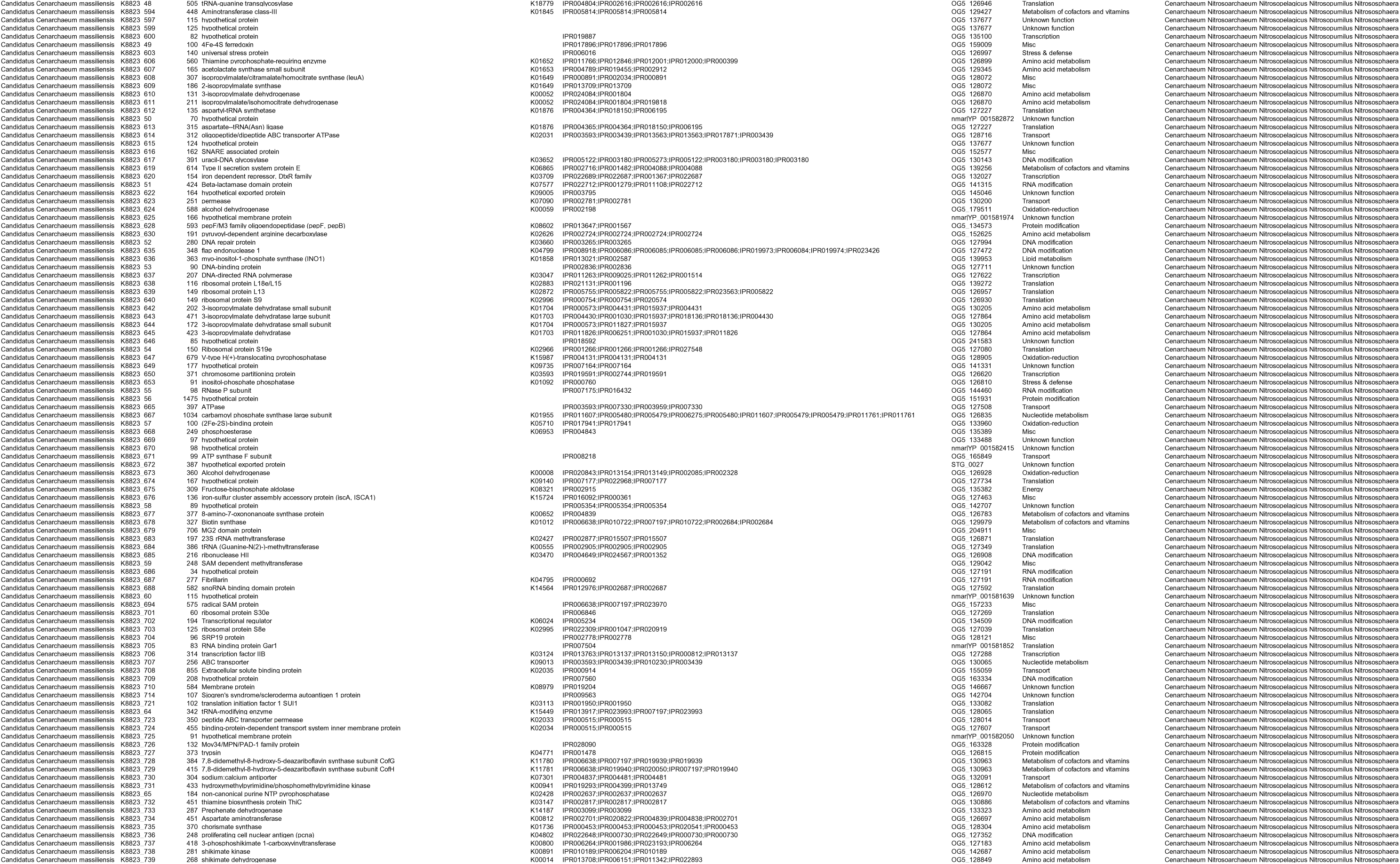
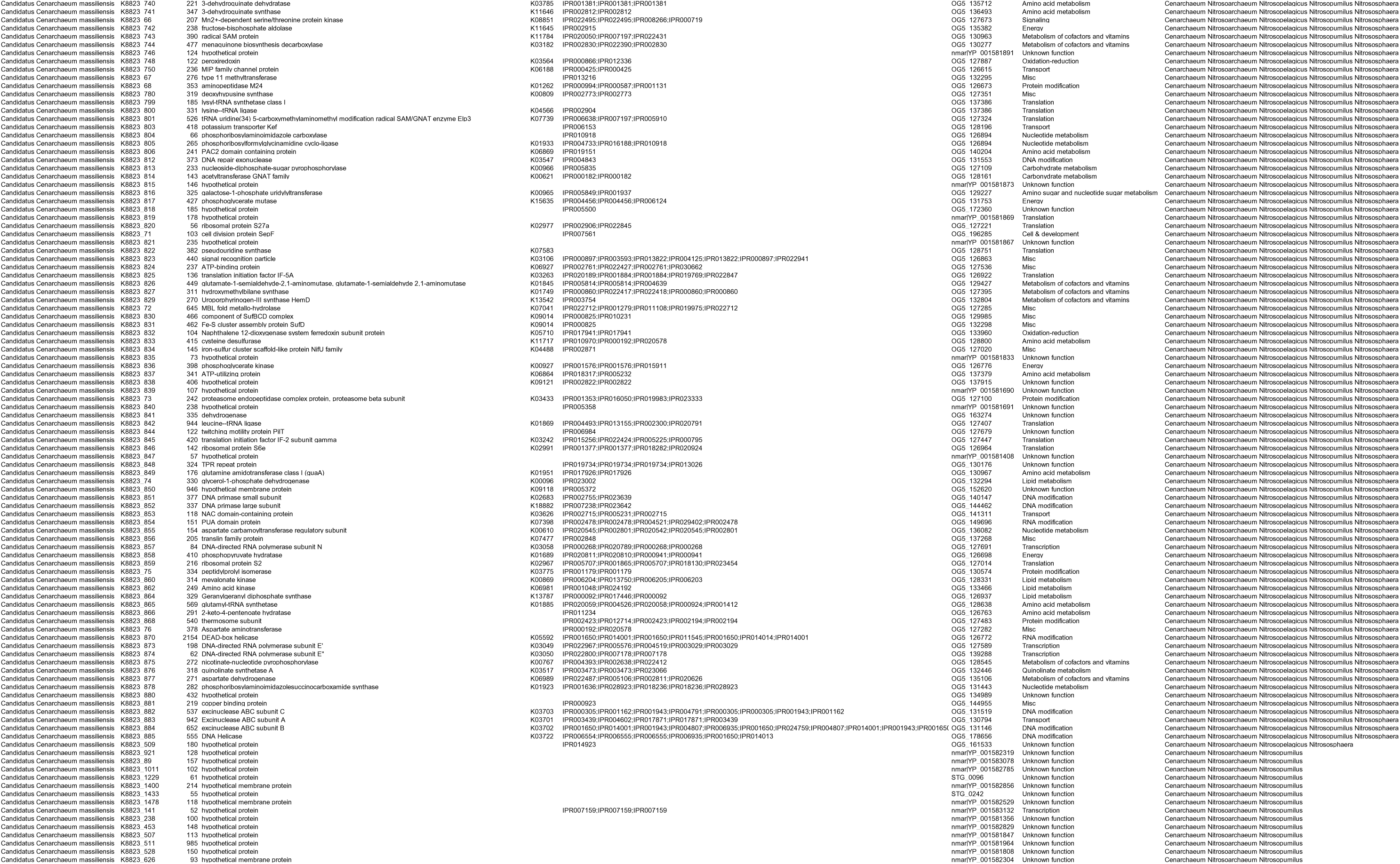
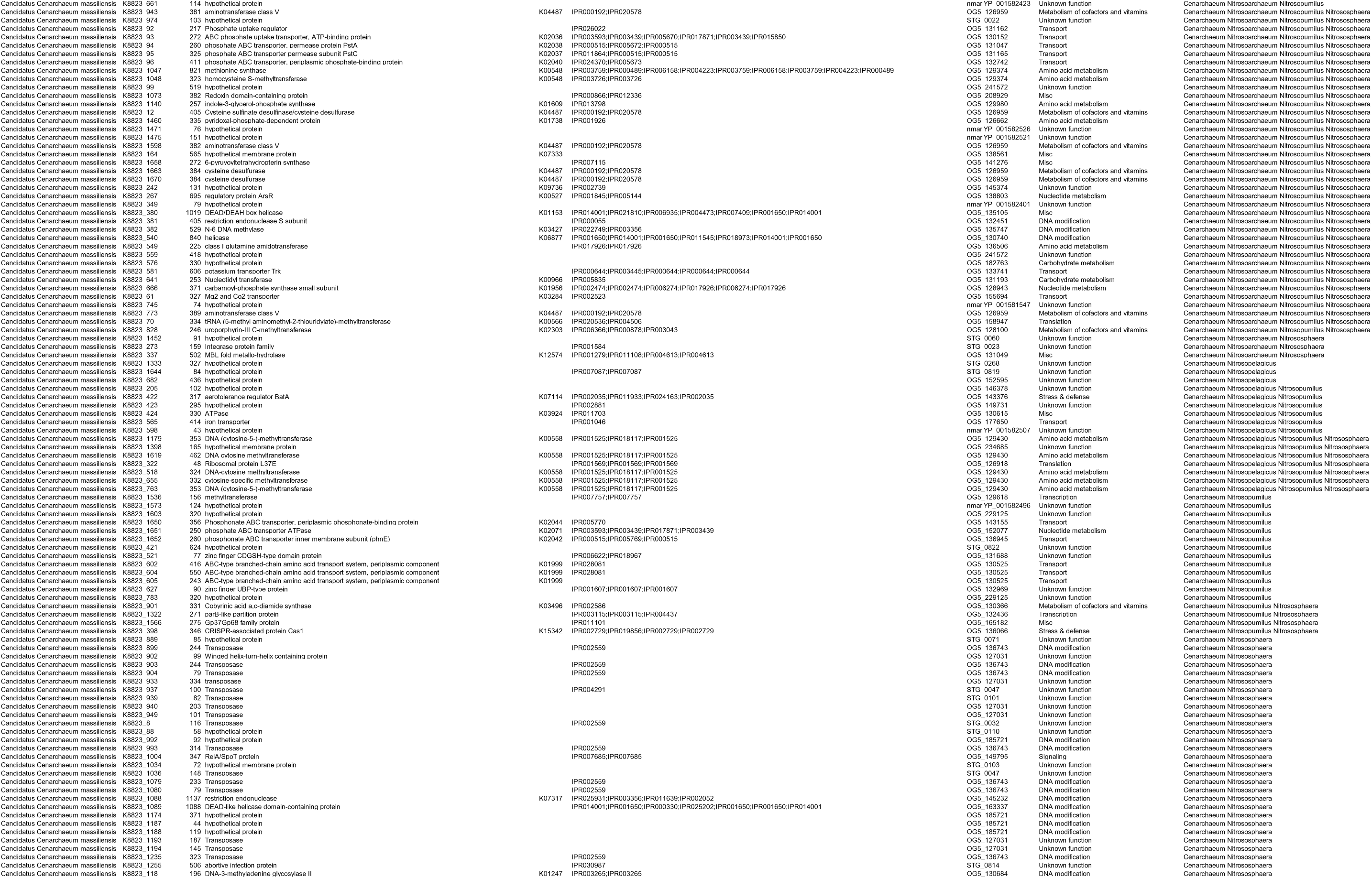
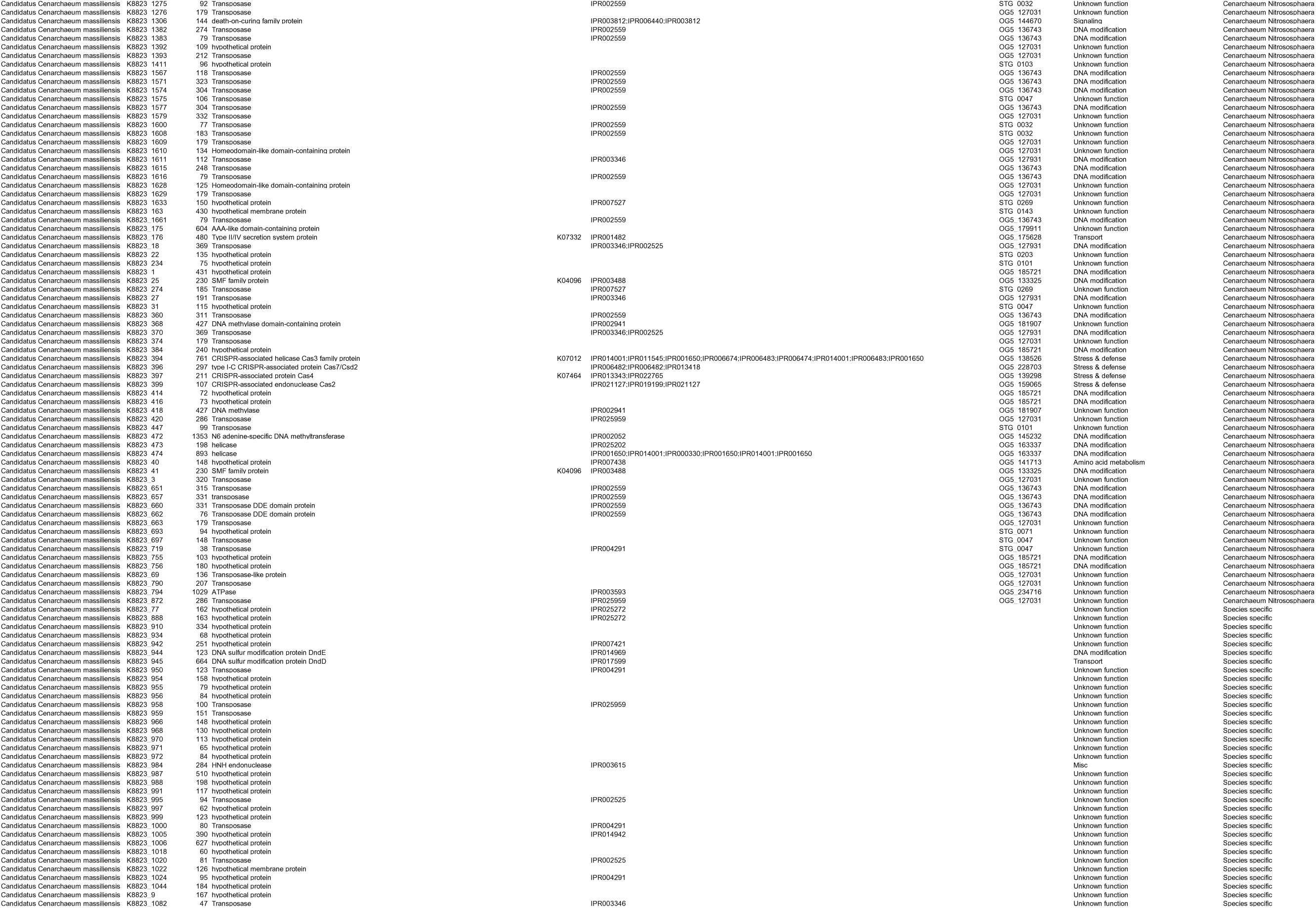
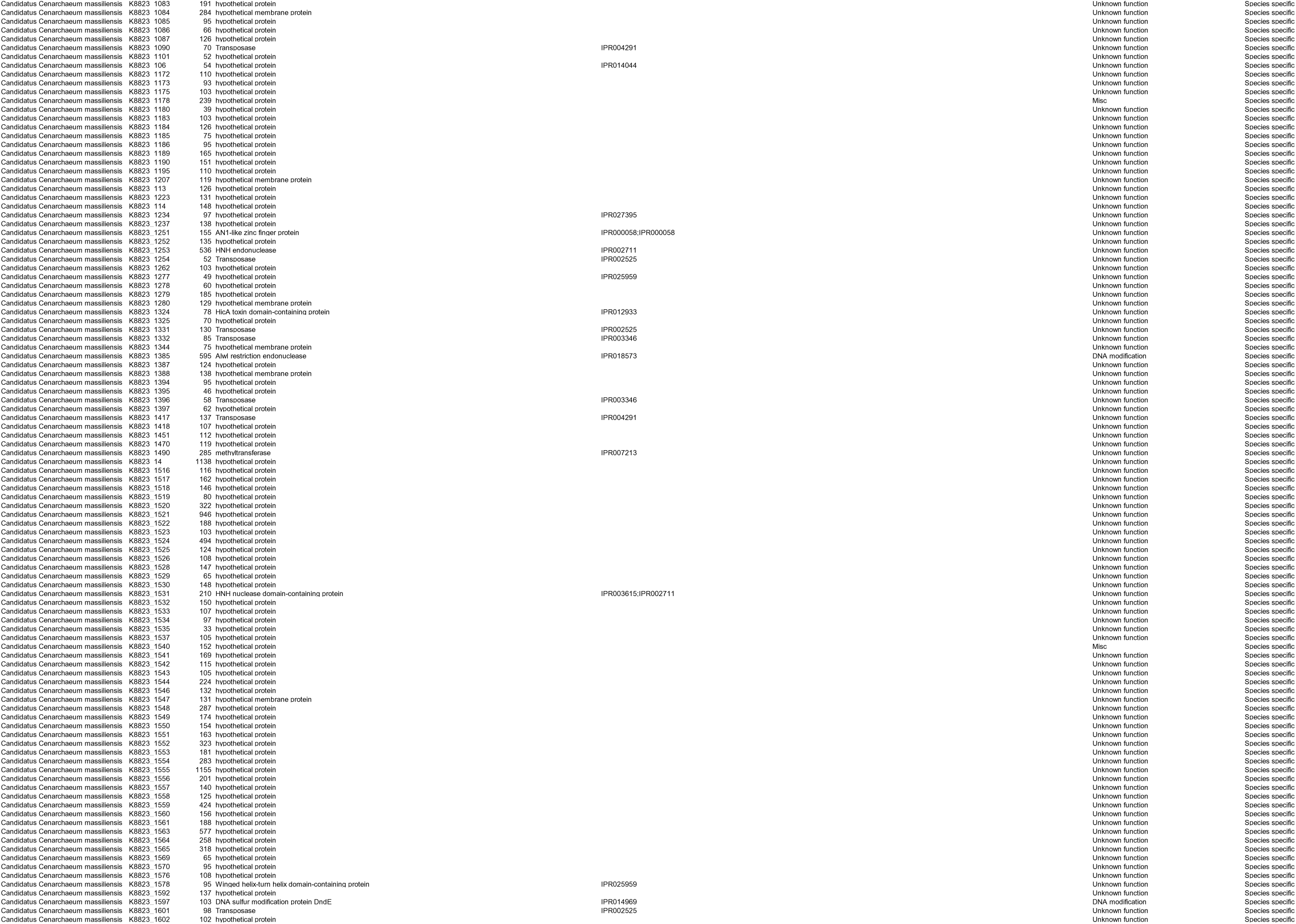
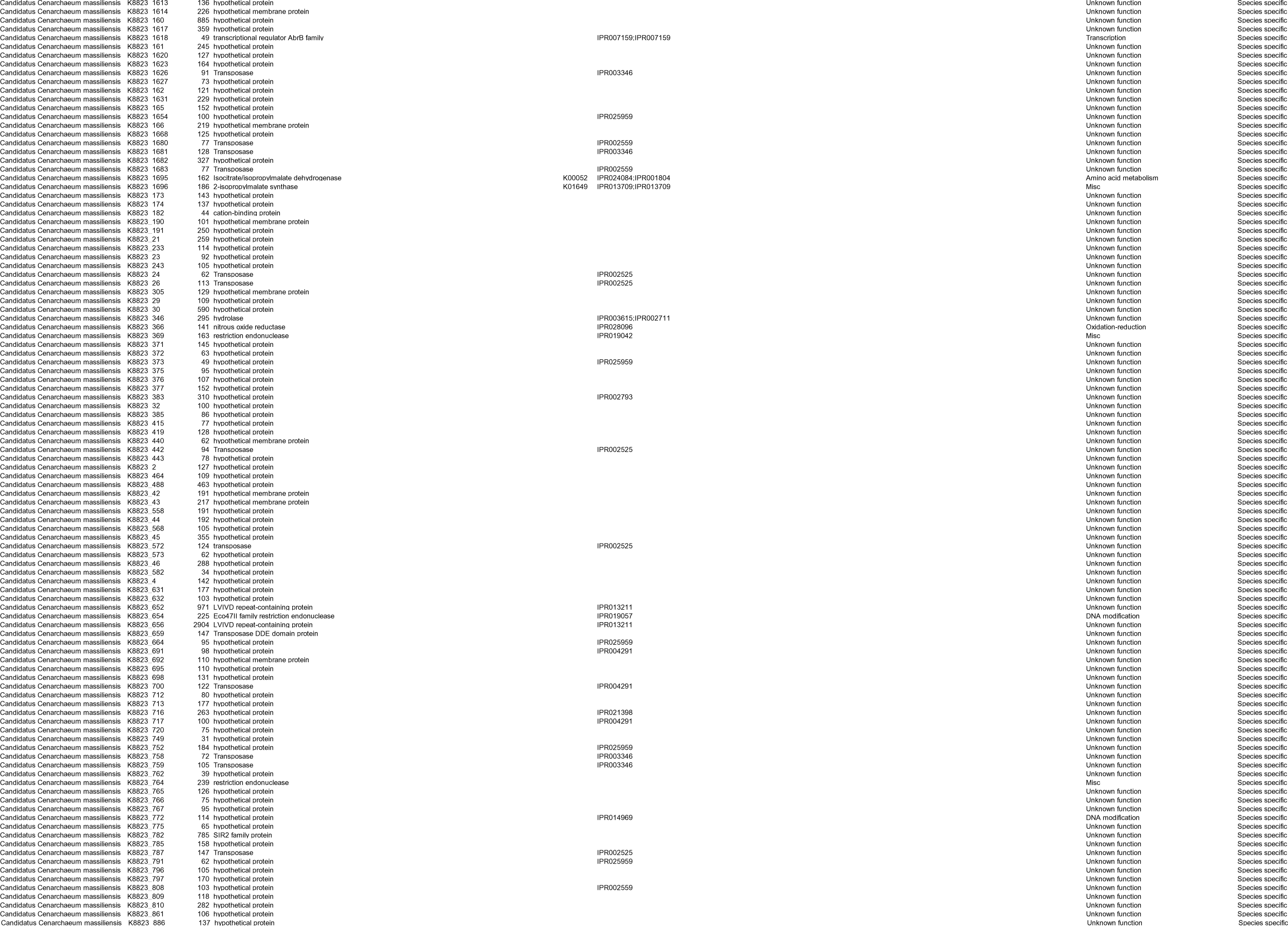
Annotation of the predicted protein-coding genes of Candidatus Cenarchaeum massilliensis.

**Supplementary figure 2:**
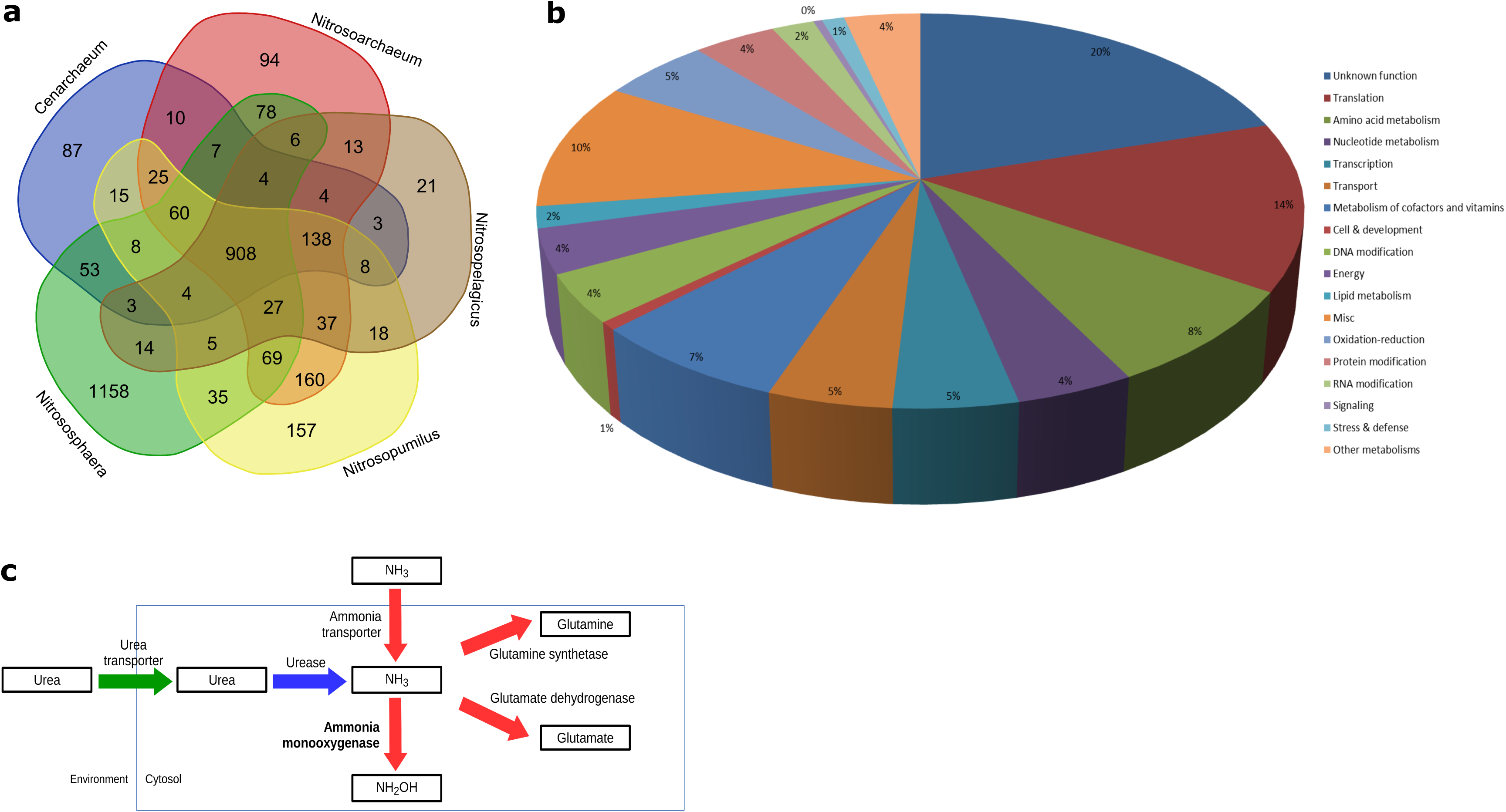
**Characterization of the genome of *Candidatus Cenarchaeum massiliensis***, the main symbiont of O. minuta, compared to other Thaumarchaeota species. **a** :Five-way-Venn-diagram illustrating orthologous genes clusters generated using OrthoMCL analysis on predicted proteins of *Cenarchaeum symbiosum* and *Cenarchaeum massiliensis* (blue circle), *Nitrosoarchaeum koreensis* and *Nitrosoarchaeum limnia*(red circle), *Nitrosopelagicus brevis* (brown circle), *Nitrosopumilus maritimus* and N*itrosopumilus koreensis*(yellow circle) and three soil Thaumarchaeota (*Nitrososphaera viennensis, Nitrososphaera gargensis* and *Nitrososphaera evergladensis*; green circle). Venn-diagram is generated using a webtool (http://bioinformatics.psb.ugent.be/webtools/Venn/). **b**: Functional classes of the 908 Thaumarchaeota conserved gene families. **c**: The nitrogen metabolism in Thaumarchaeota. Conserved steps in all Thaumarchaeota (red arrows), conserved step only in soil Thaumarchaeota and Cenarchaeum symbiosum (blue arrow) and conserved step only in soil Thaumarchaeota (green arrow) are shown.

**Supplementary figure 3:**
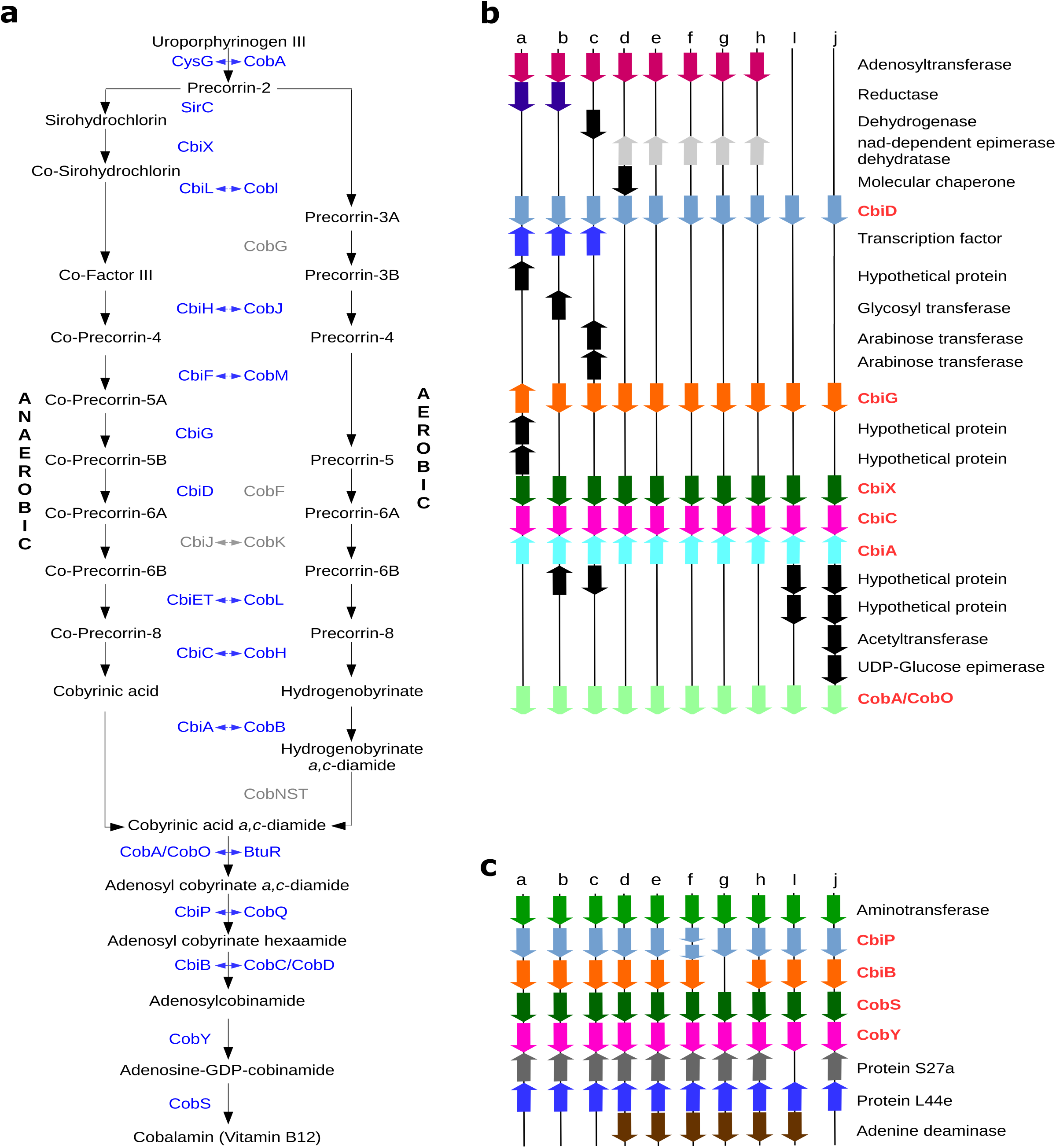
The cobalamin synthesis genes in Thaumarchaeota. **a:**Pathway of the cobalamin synthesis adapted from Doxey et al., 2014. Blue arrows indicate shared enzymatic steps between aerobic and anaerobic pathways. All enzymes of the anaerobic pathway are retrieved (indicated in blue) except CbiJ (indicated in grey). **b**:Microsynteny analysis of Thaumarchaeota genomic segments bearing six genes coding for early enzymatic steps of cobalamin synthesis. **c**:Microsynteny analysis of Thaumarchaeota genomic segments bearing four genes coding for late enzymatic steps of cobalamin synthesis. Segments “a” to “j” correspond to genomic segments from Nitrososphaera gargensis, Nitrososphaera viennensis, Nitrososphaera evergladensis,Nitrosopelagicus brevis, Nitrosoarchaeum koreensis, Nitrosoarchaeum limnia, Nitrosopumilus koreensis, Nitrosopumilus maritimus, Cenarchaeum symbiosumand Candidatus Cenarchaeum massiliensis, respectively. Arrows of the same color indicate orthologous genes and black arrows represent genes without orthology relationship in these regions from OrthoMCL analysis.

**Supplementary Figure 4:**
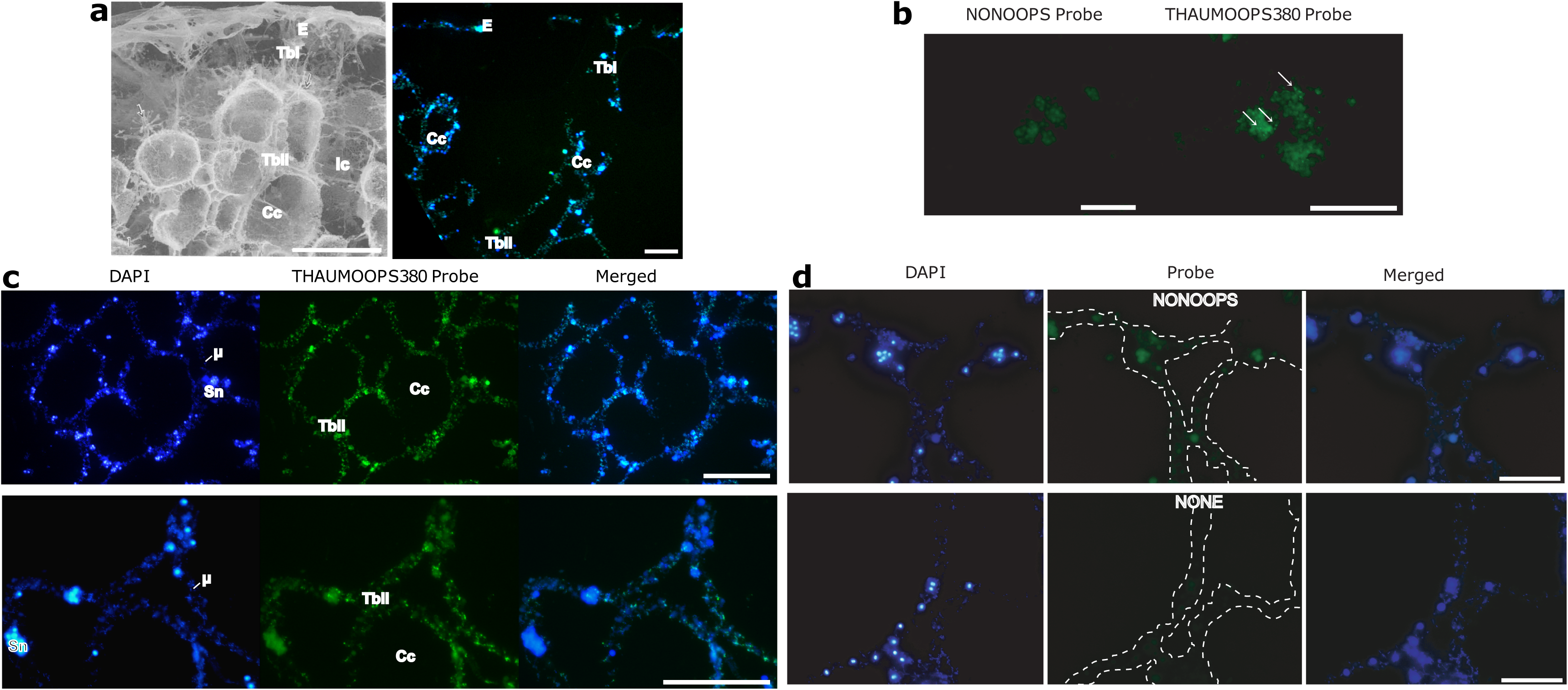
Localization by Card-FISH of the thaumarcheota *Candidatus Cenarchaeum massiliens* in the tissues of *Oopsacas minuta.* **a:** Confocal picture (on the right) showing its localization in the the different parts of the trabecular syncitium (Tb) : the Ectosome (E), the Choanochambers (CC), the inhalant canals (Ic), recognized by comparison with SEM pictures (on the left) (Boury-Esnault and Vacelet, 1994) rscale bars = 150 μm. **b:** Detection of *Ca. C. massiliensis* in embryos (right; THAUMOOPS380-HRP probe) and negative control (left, NONOOPS probe). **c:** Localization in the trabecular syncitium shown at higher magnification in two different individuals with the THAUMOOPS380-HRP probe (green) and conterstained with DAPI (blue), µ: microorganisms DNA, Sn: Sponge nuclei, scale bars = 70µm **d:** Card-FISH negative controls: with a random probe with no 100% match in the metagenome (NONOOPS-HRP), or no probe, evidencing that sponge cells have an endogenous peroxidase activity uncomplety abolished by 0.3% H2O2.

### Genomic characteristics of *Oopascas minuta*

**Supplementary Table 5a:** Genome features of the glass sponge *Oopsacas minuta*, compared to other sponge genomes from the three other classes.

| species | Hexactinellida |  | Demospongiae |  |  |  | Homoscleromorpha | Calcarea |
| --- | --- | --- | --- | --- | --- | --- | --- | --- |
|  | *O. minuta*<br>(genome mode) | *O. minuta*<br>(Augustus aa set) | *E. muelleri* (genome mode) | *E. muelleri*<br>(Augustus aa set) | *A. queenslandica* | *T. wilhelma* | *O. pearsei* | *S. ciliatum* |
| Total # of core genes queried | 303 | 303 | 303 | 303 | 303 | 303 | 303 | 303 |
| # of core genes detected | 282 (93.07%) | 278 (91.75%) | 243 (80.20%) | 254 (83.83%) | 274 (90.43%) | 258 (85.15%) | 207 (68.32%) | 240 (79.21%) |
| Complete + Partial | 285 (94.06%) | 284 (93.73%) | 254 (83.83%) | 273 (90.10%) | 281 (92.74%) | 275 (90.76%) | 237 (78.22%) | 259 (85.48%) |
| # of missing core genes | 18 (5.94%) | 19 (6.27%) | 49 (16.17%) | 30 (9.90%) | 22 (7.26%) | 28 (9.24%) | 66 (21.78%) | 44 (14.52%) |
| Average # of orthologues per core genes | 1.01 | 1.01 | 1.12 | 1.22 | 1.04 | 1.05 | 1 | 1.07 |
| % of detected core genes that have more than 1 ortholog | 1.06 | 0.72 | 10.7 | 19.6 | 4.38 | 4.26 | 0 | 6.67 |

**Supplementary Table 5b:**
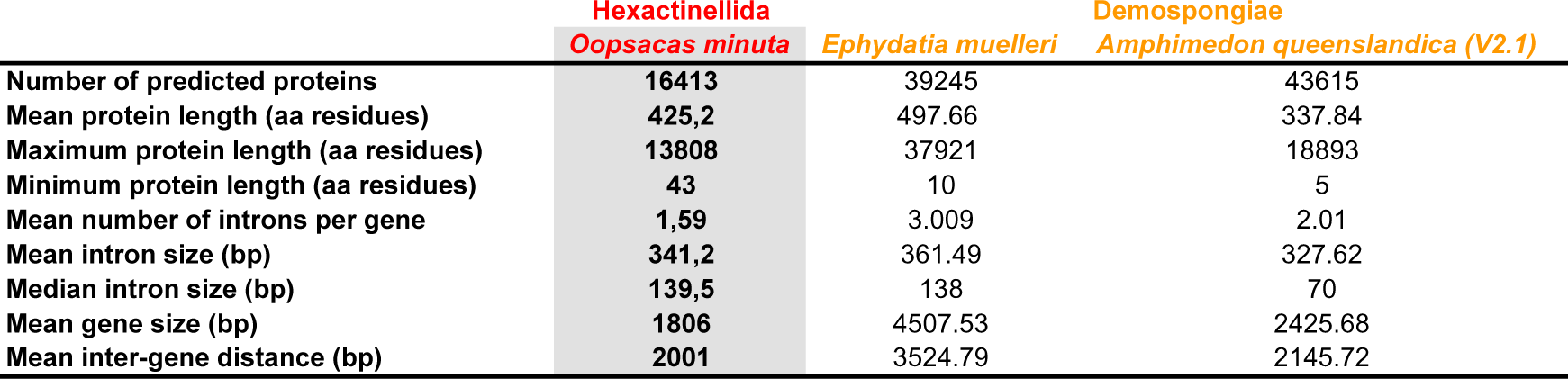
Mean size of coding and non-coding sequences in *Oopascas minuta* compared to two demosponges.

**Supplementary Table 6:** Respective number of predicted and annotated sequences in the genome of *Oopsacas minuta*.

| Categories |  | Number |
| --- | --- | --- |
| predicted genes |  | 16413 |
| Annotated Genes |  | 7737 |
| GO Terms | Biological | 4092 |
|  | Cellular | 2042 |
|  | Function | 6704 |
|  | Total | 12838 |

**Supplementary Figure 5:**
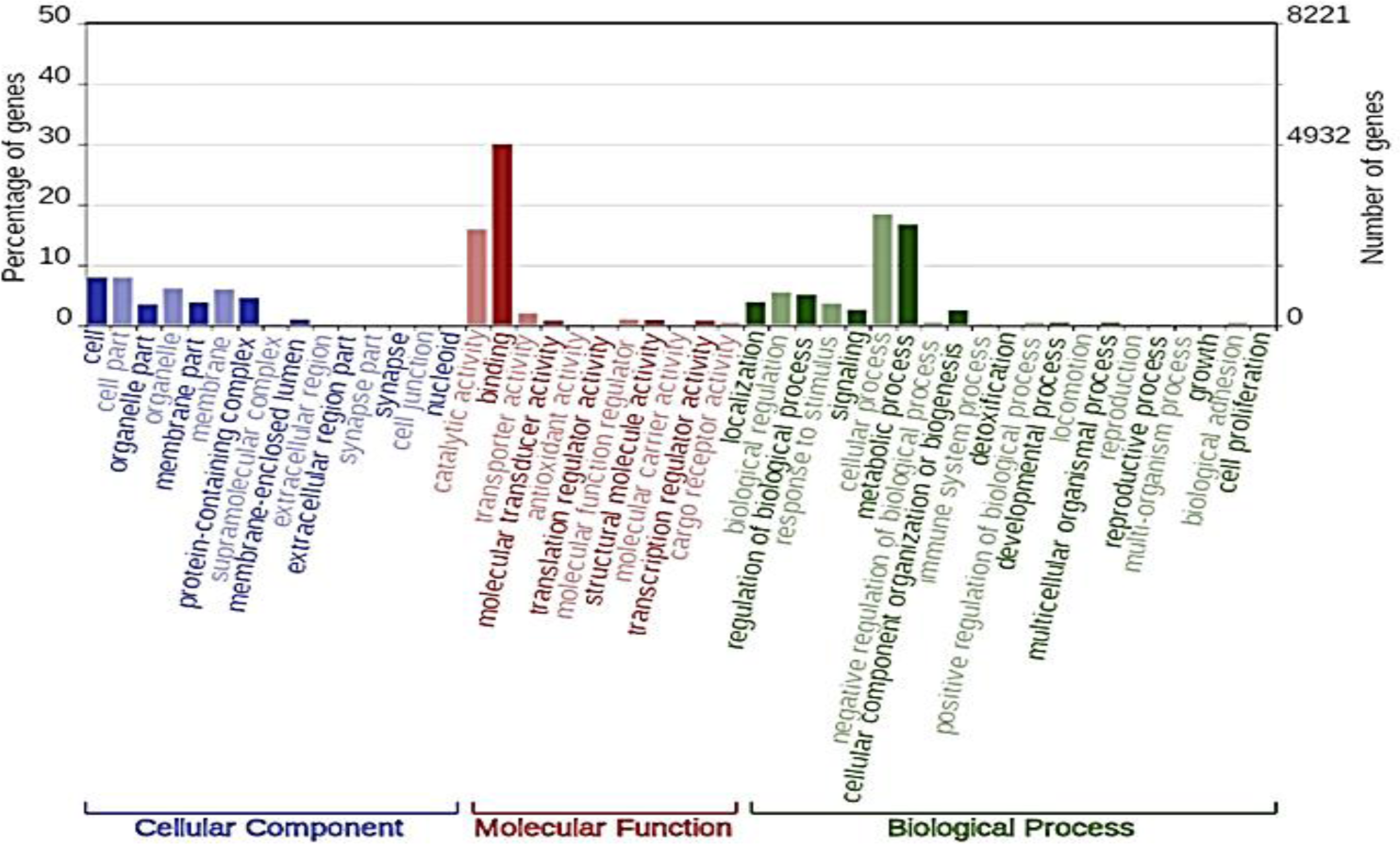
**Gene ontology at level two** based on the interproscan domain search over different databases.

### Survey and analysis of epithelial gene candidates

**Supplementary Table 7:** **Blast search of genes involved in epithelial functions in bilaterians** (only genes with significative hits are indicated)

| QUERY |  |  |  |  | REVERSE BLAST |
| --- | --- | --- | --- | --- | --- |
| Name of searched protein | Query ID | Accession number | Hit ID Oopsacas | e-value | Best Hit ID |
| Contactin | Contactin (*Aphrocalistes vastus*) | Ava_Contig_16553-1 | LOD99_12497 | 0 | Contactin 2 |
| | | | LOD99_4945 | $3.45 \times 10^{-32}$ | sidekick |
| | | | LOD99_2903 | $4.96 \times 10^{-20}$ | receptor-type tyrosine-protein phosphatase F/DSCAM/hemicentin |
| | | | LOD99_13407 | $6.41 \times 10^{-13}$ | DSCAM/sidekick |
| Integrin | Integrin alpha (*Aphrocalistes vastus*) | Ava_Contig_11100-1_and 11100-2 | LOD99_15309 | 0 | Integrin alpha 9 Amphimedon |
|  |  |  | LOD99_15308 | 0 | integrin alpha 9 Amphimedon |
| | | | LOD99_13917 | $7.61 \times 10^{-82}$ | integrin alpha -V (Xiphophorus couchianus) |
| | | | LOD99_9146 | $9.47 \times 10^{-70}$ | integrin alpha-PS2-like [Limulus polyphemus] |
| | | | LOD99_6533 | $7.55 \times 10^{-22}$ | integrin alpha-6 isoform X1 [Canis lupus familiaris] |
|  | Integrin beta (*Aphrocalistes vastus*) | Ava_Contig_13475-2_integrin-beta- | LOD99_8099 | 0 | integrin beta-7 [Microcaecilia unicolor] |
| | | | LOD99_2393 | $3.06 \times 10^{-69}$ | integrin beta-C subunit precursor [Strongylocentrotus purpuratus] |
| | | | LOD99_12882 | $2.35 \times 10^{-54}$ | integrin beta-1-like [Acropora millepora] |
| Nidogen | Nidogen (A. vastus) | Ava_Contig_19310_nidogen | **LOD99_13205** | $4.36 \times 10^{-118}$ | protogenin-like protein [Leptotrombidium deliense]/netrin |
| | | | LOD99_13214 | $3.17 \times 10^{-58}$ | receptor-type tyrosine-protein phosphatase epsilon-like [Amphimedon queenslandica] |
| Laminin | Laminin beta (A. vastus) | Ava_Contig_18402_Laminin-beta | **LOD99_7300** | 0 | laminin subunit gamma-1-like [Amphimedon queenslandica] |
| | | | **LOD99_12851** | $4.85 \times 10^{-118}$ | laminin subunit gamma-1-like [Acropora millepora] |
| | | | **LOD99_7636** | $1.90 \times 10^{-105}$ | laminin subunit alpha-like [Acanthaster planci] |
| | | | **LOD99_1235** | $1.92 \times 10^{-44}$ | |
| | | | LOD99_8477 | $1.71 \times 10^{-69}$ | Usherin (A. queenslandica) |

**Supplementary Figure 6a:**
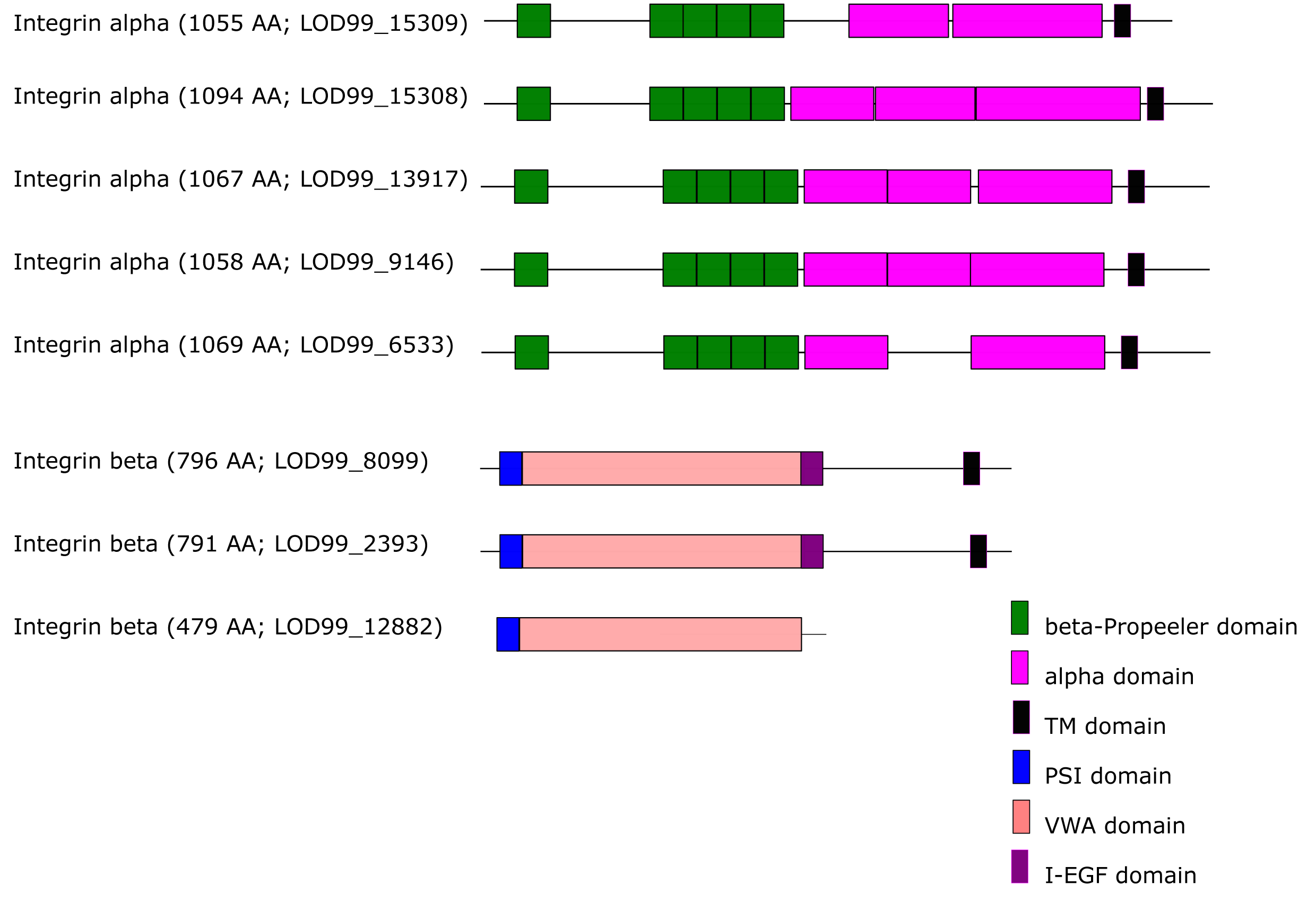
**Domain prediction of *Oopsacas* integrins** using Pfam. 5 predicted alpha-integrins of O. minuta have a domain structure similar to those observed in other metazoans: a globular head region in the extracellular domain consisting in five or six repeats of about 60 amino acids that fold into a five/six-bladed β propeller followed by a large integrin alpha-2 domain. The 3 proteins predicted as beta-integrins share an integrin plexin domain (PSI) in the Nt globular part, an integrin beta chain VWA domain, an integrin beta epidermal growth factor like domain 1 as reported in other metazoans.

**Supplementary Figure 6b:**
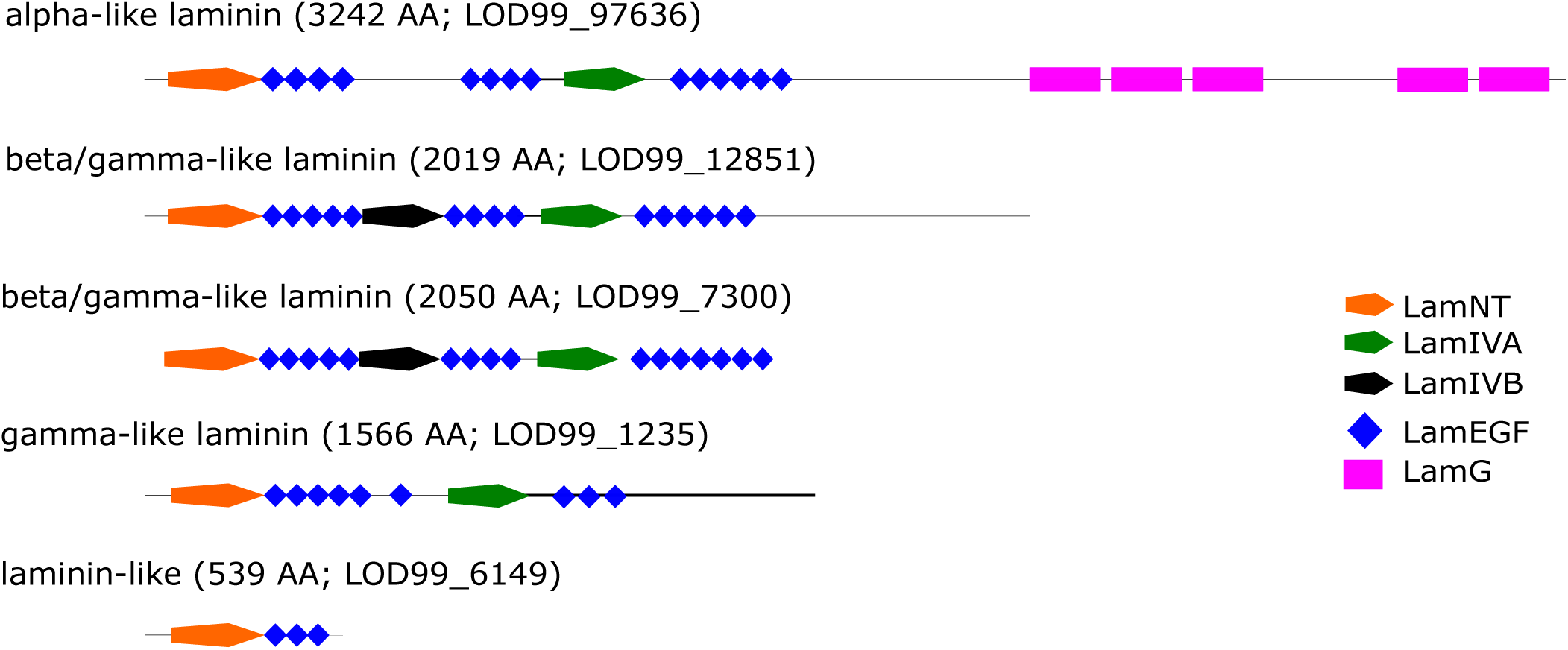
**Protein Domain prediction of *Oopsacas* laminins** using Pfam

### Survey of candidate genes for multiciliogenesis

**Supplementary Table 8:**
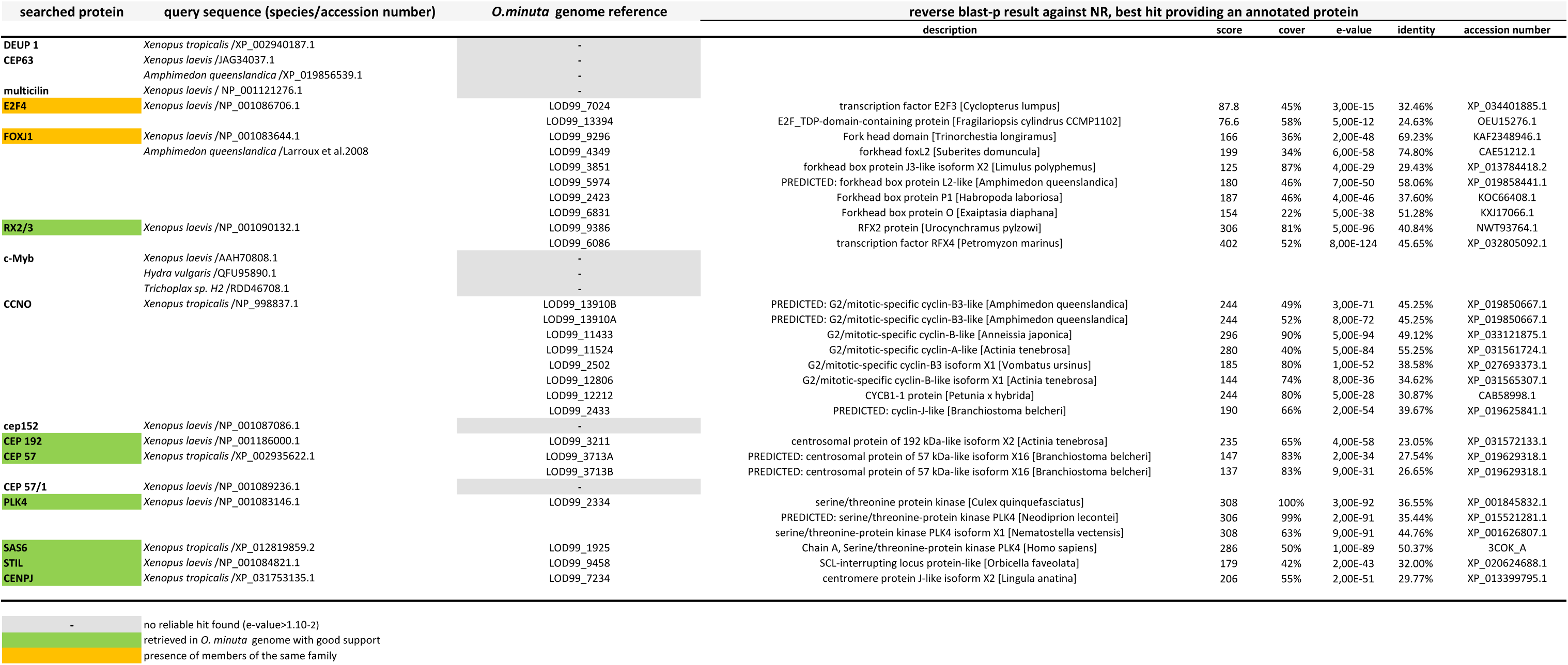
Summary of Blast search for candidate genes for multiciliogenesis against the predicted proteome of *Oopsacas minuta*, and reverse blast best hits against NR.

### Survey of members of key ancestral signaling pathways

**Supplementary Table 9:**
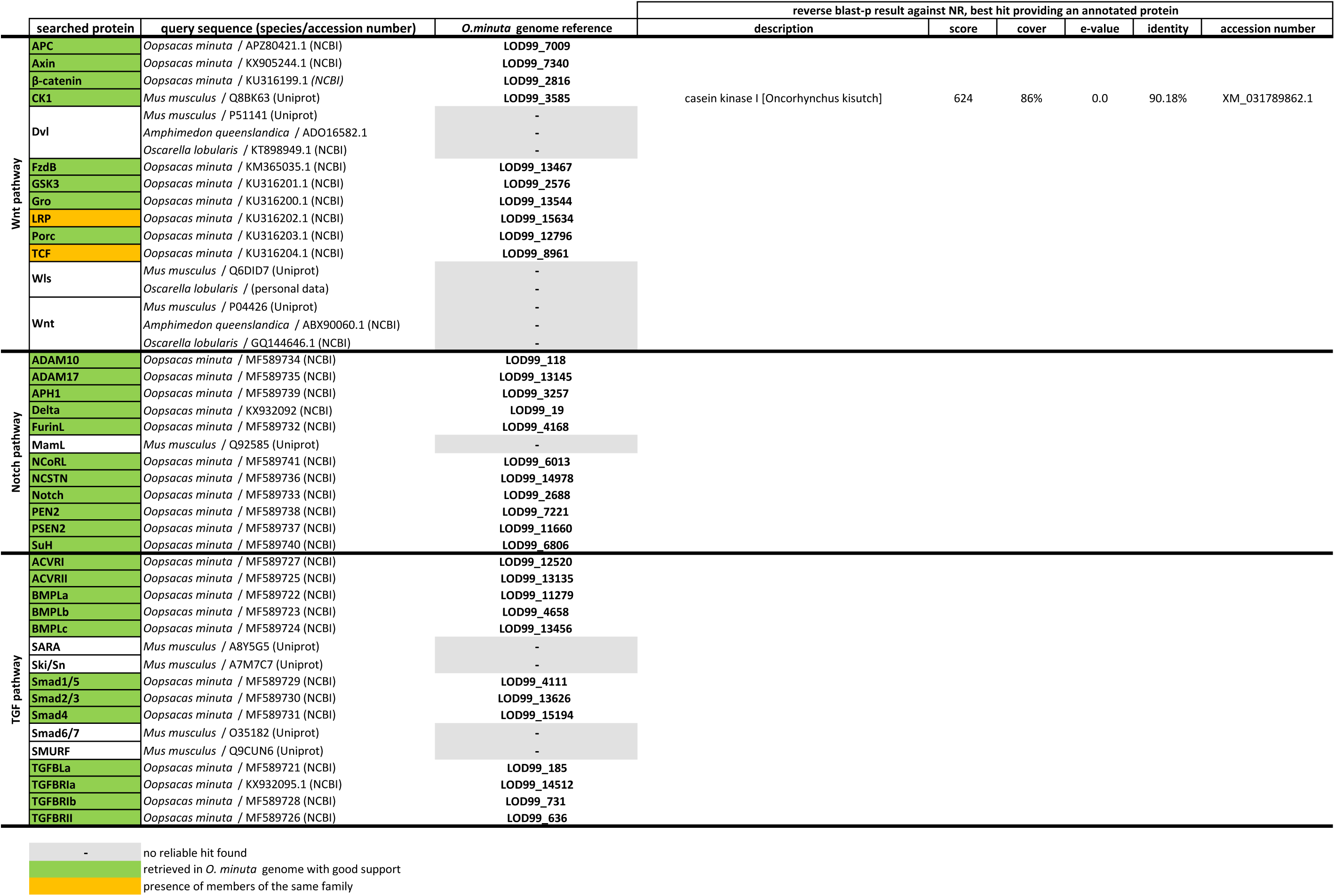
Blast Search of core genes involved in three main ancestral signalling pathways.

**Supplementary Table 10:**
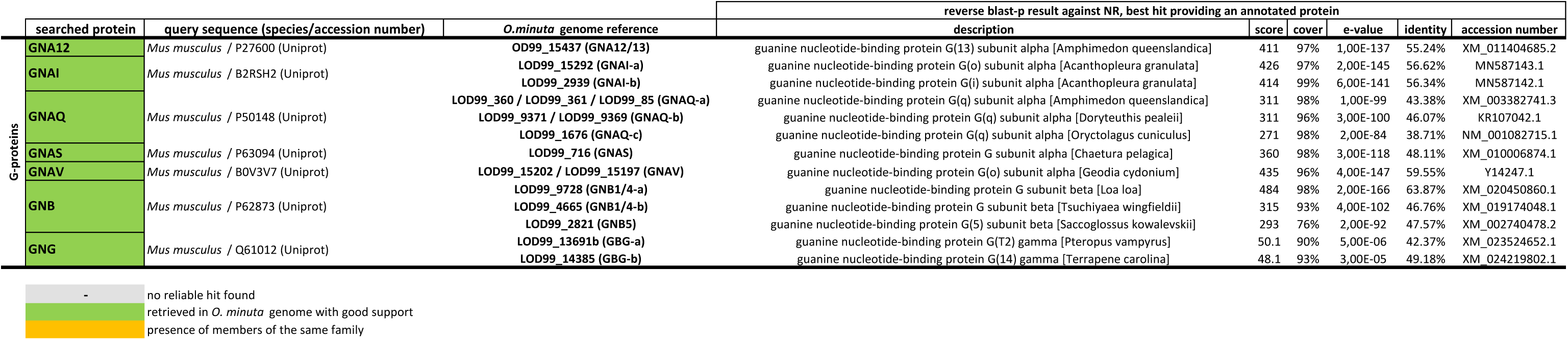
Blast search results of G-proteins against the predicted proteome of *Oopsacas minuta* and reverse blast best hit scores.

### Survey and analyses of transcription factors

**Supplementary Table 11:**
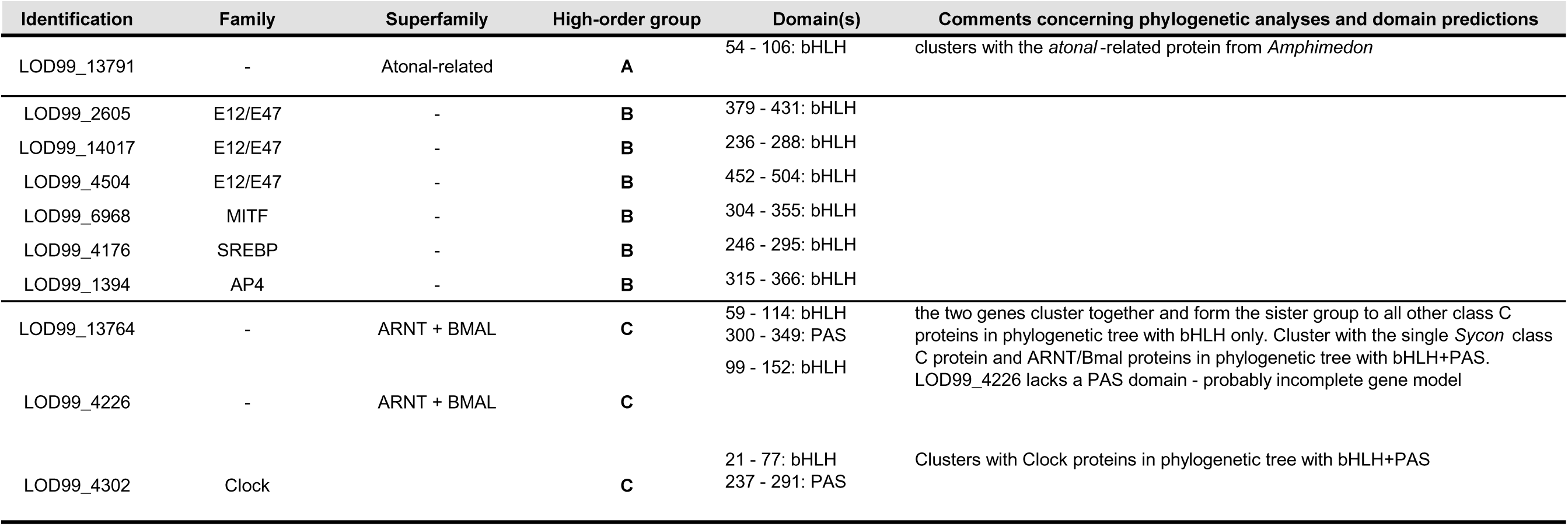
Summary of Blast search, protein domain and phylogenetic analyses concerning basic Helix Loop Helix (bHLH) transcription factors.

**Supplementary Table 12:**
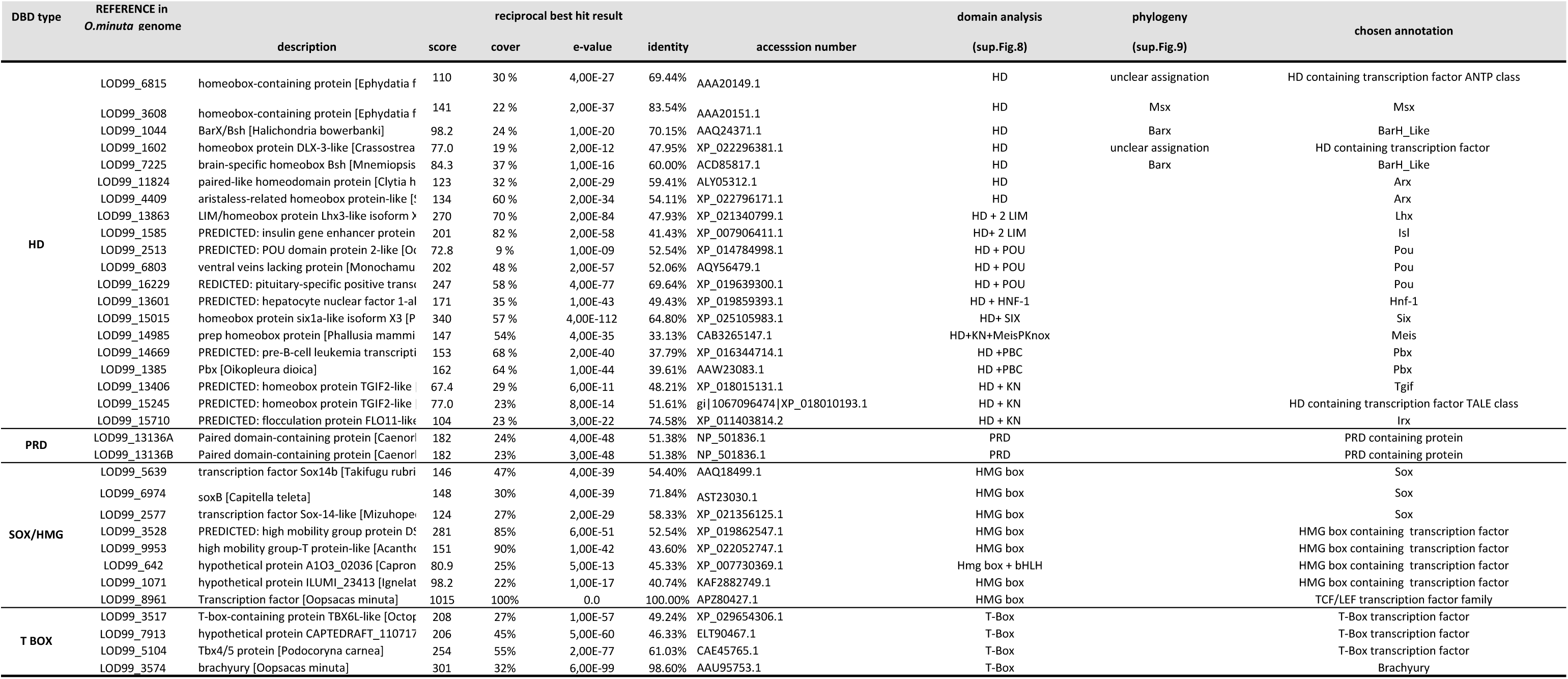
Summary of Blast search, domain and phylogenetic analyses performed of various transcription factor types with particular focus on HD transcription factors.

**Supplementary Figure 7:**
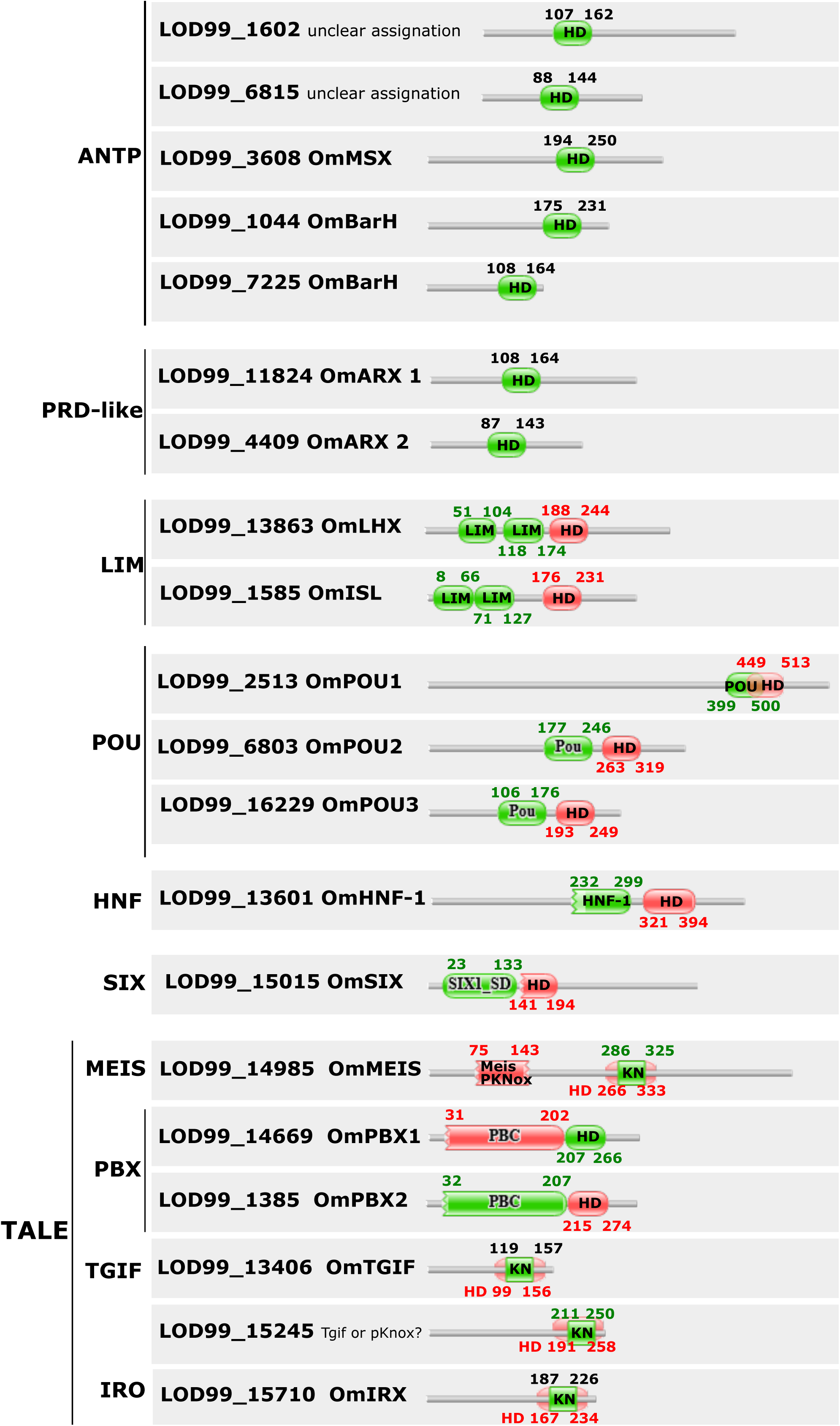
**Domain prediction of HD transcription factors** (by Pfam and checked with interproscan when needed)

**Supplementary Figure 8:**
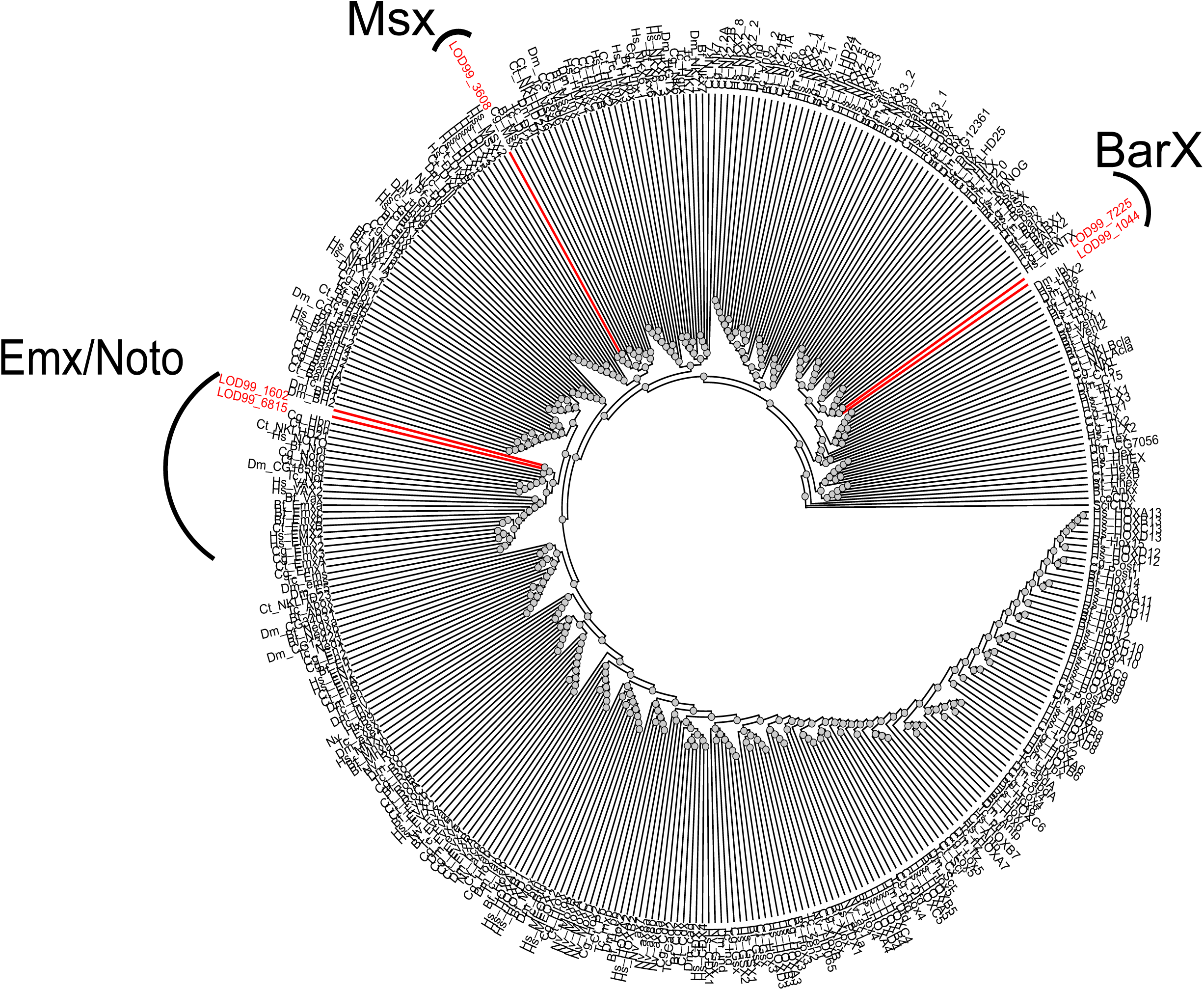
**Phylogenetic position of homeobox transcription factors of *O. minuta*** assigned to the Antennapedia class (in red). The *Oopsacas* HD sequences were integrated and aligned using MUSCLE in the dataset used by Pastrana *et al.*, 2019 gathering sequences from *Homo sapiens* (Hs), *Branchiostoma floridae* (Bf), *Drosophila melanogaster* (Dm), *Tribollium castaneum* (Tc), *Capitella teleta* (Ct), *Crassostrea gigas* (Cg), *Nematostella vectensis* (Nv).The bayesian phylogenetic analyses were performed on http://www.atgc-montpellier.fr (e-bayes option). The graphic representation was performed with Treeview.

### *Survey* and analyses *of genes involved in the bilaterian neurosensory system*

**Supplementary Table 13:**
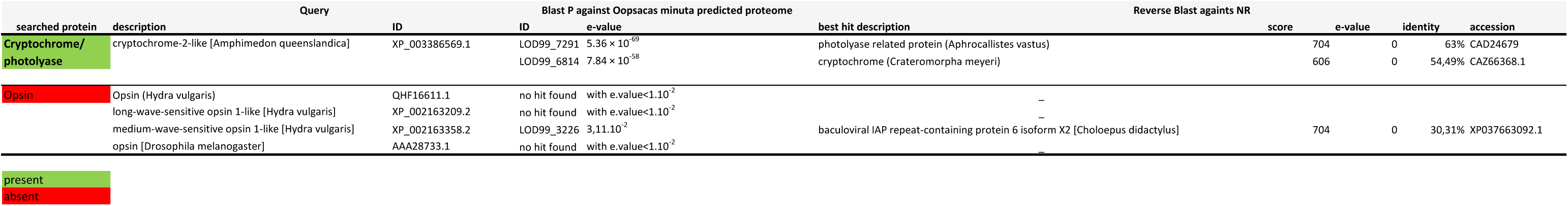
Survey of candidate genes for photoreception.

**Supplementary Table 14:**
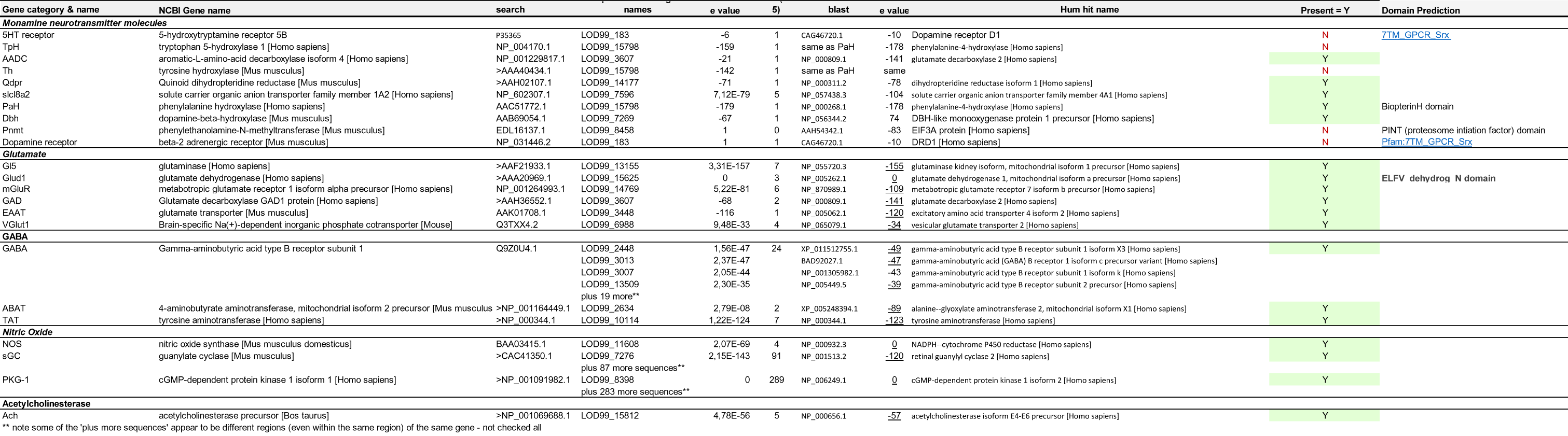
Blast search for genes encoding for proteins involved in chemical signaling against *Oopsacas* predicted proteome and reverse blast best hits againts NR.

**Supplementary Figure 9:**
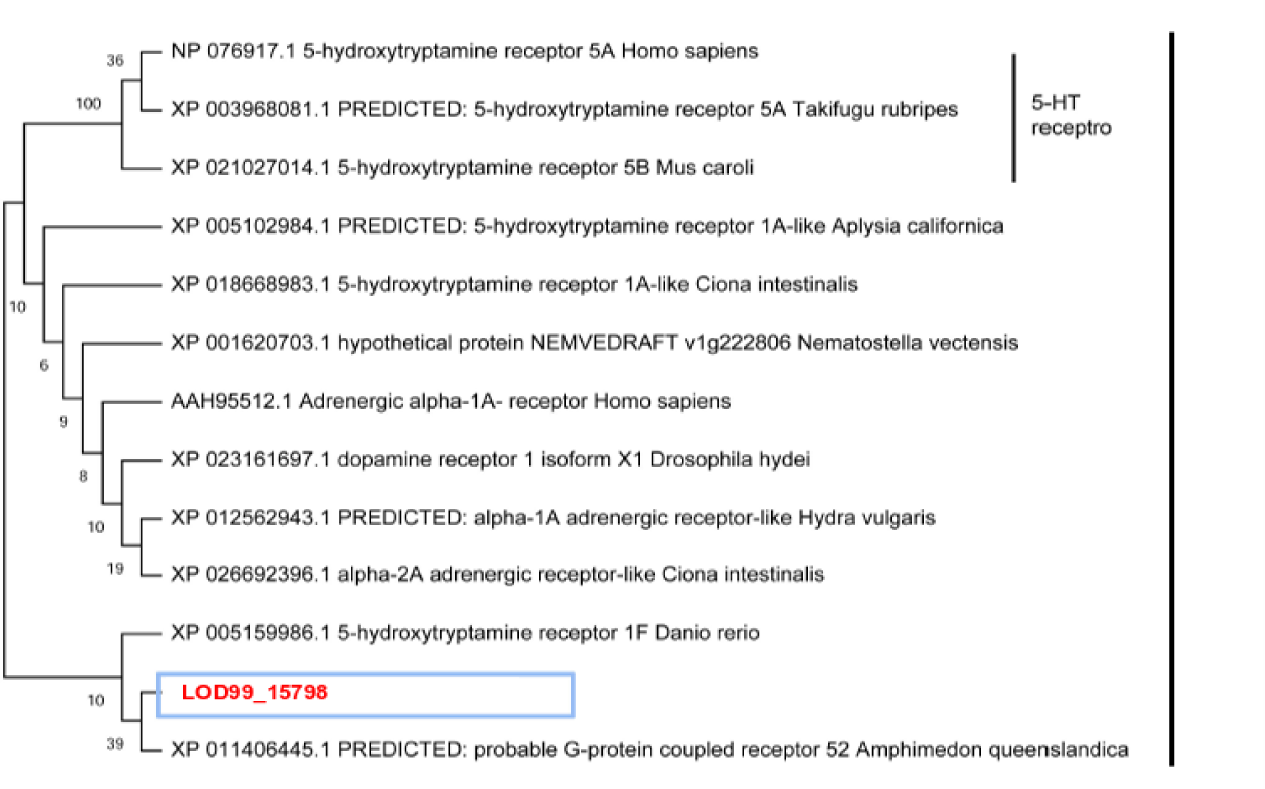
Phylogenetic position of the Oopsacas gene encoding for a monamine neurotransmitter.

**Supplementary Table 15:**
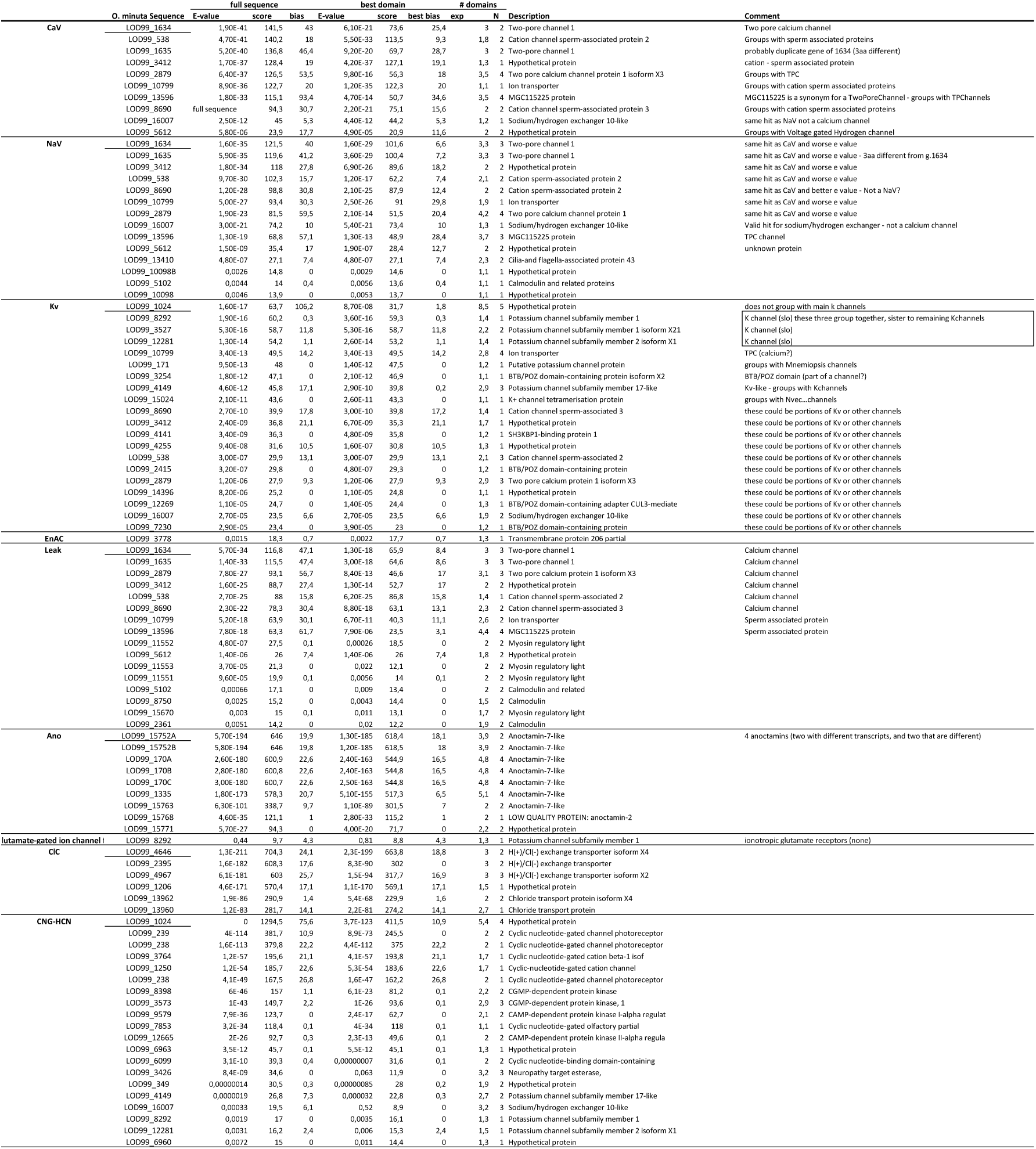
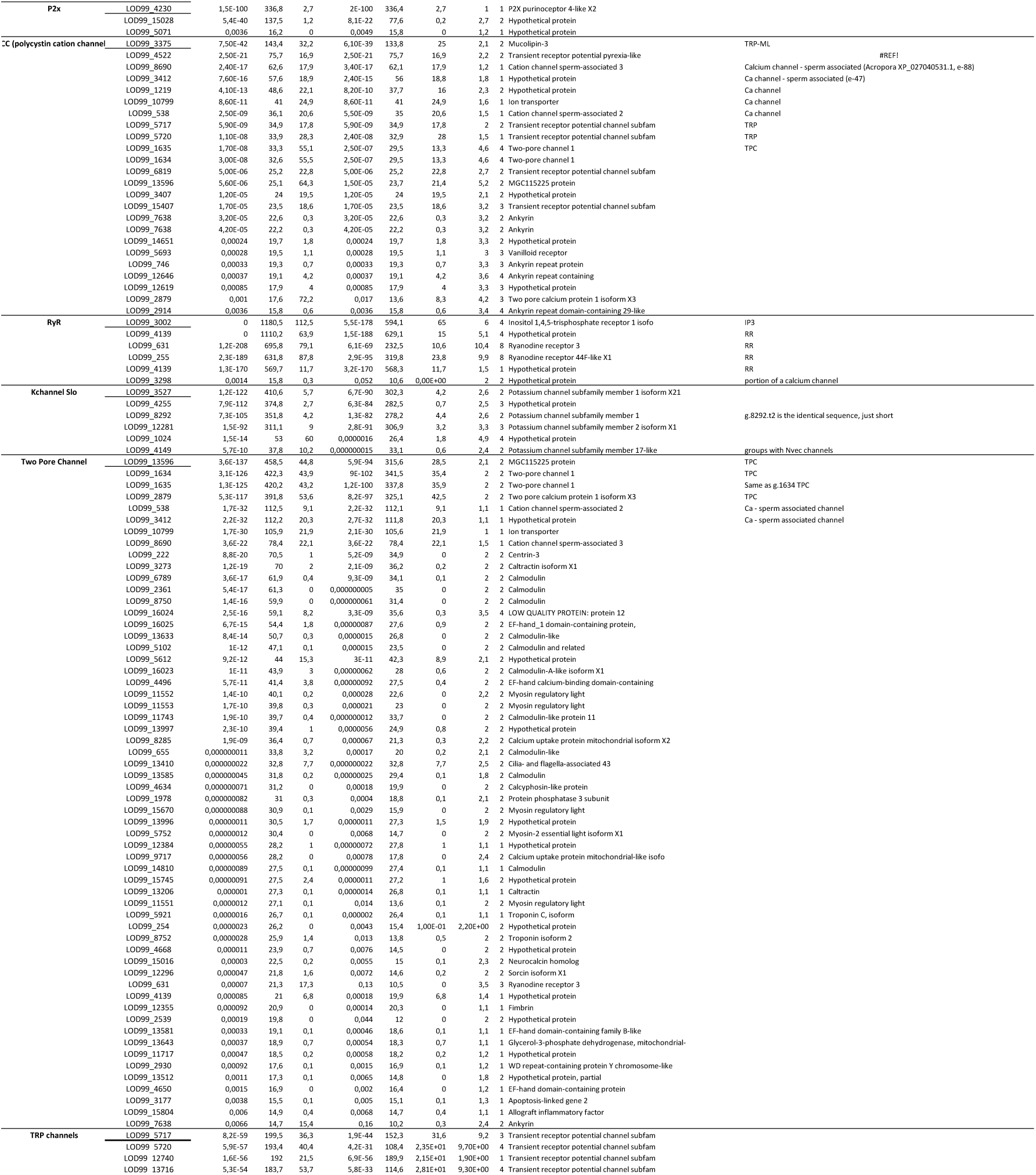
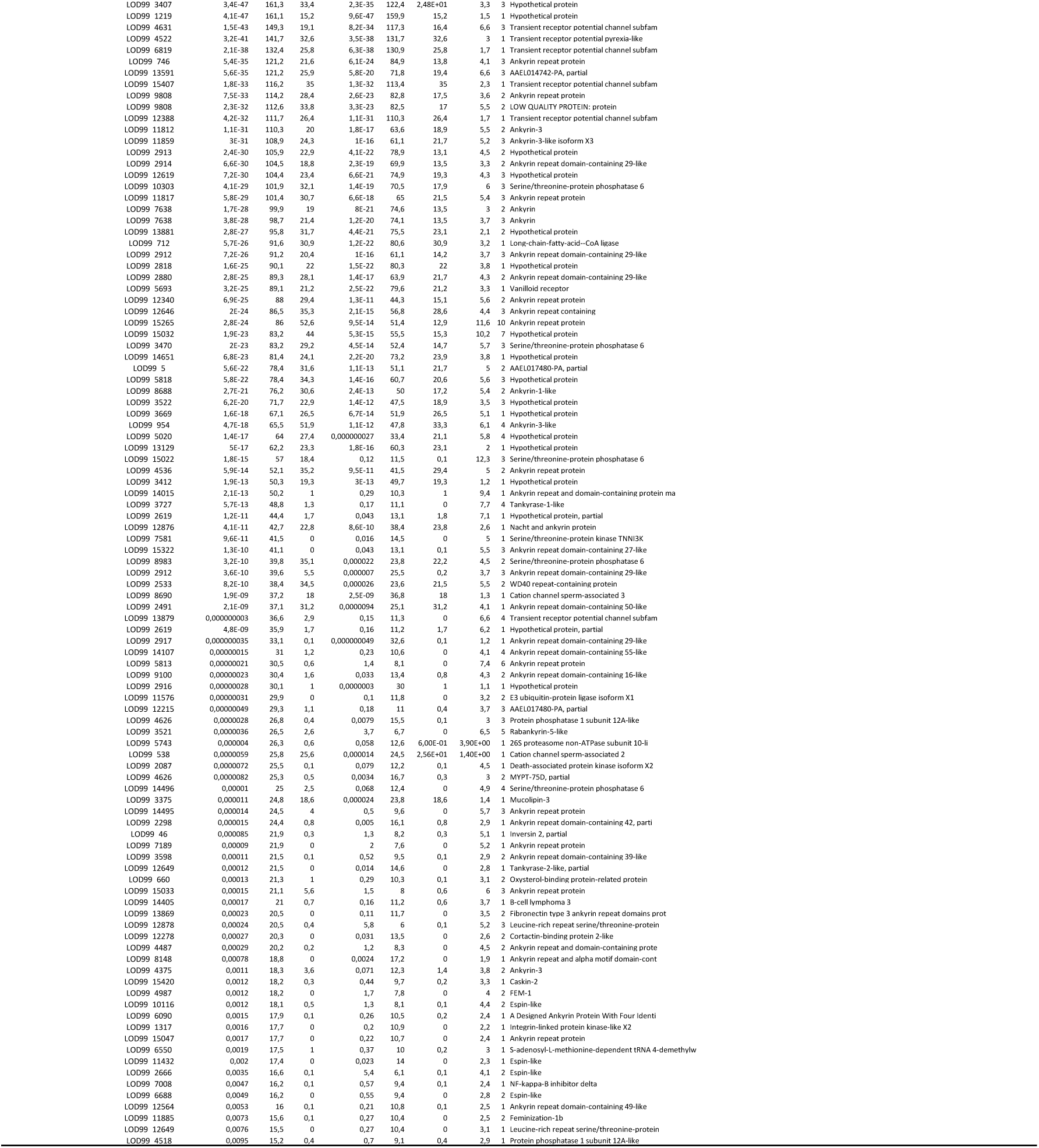
Blast results for voltage gated ion channels.

**Supplementary Figure 10:**
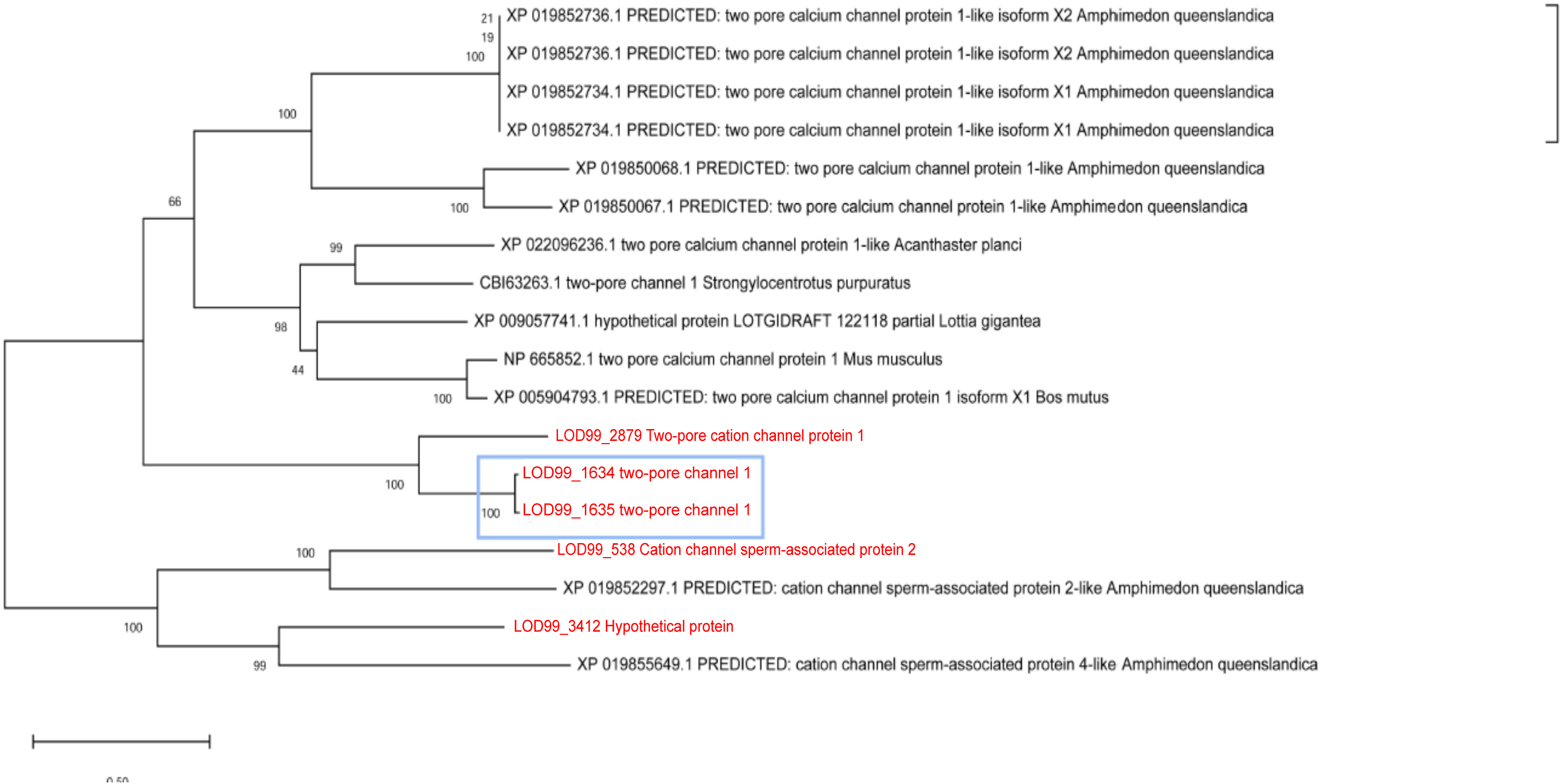
Phylogenetic position of Oopsacas genes encoding calcium channels.

**Supplementary Figure 11:**
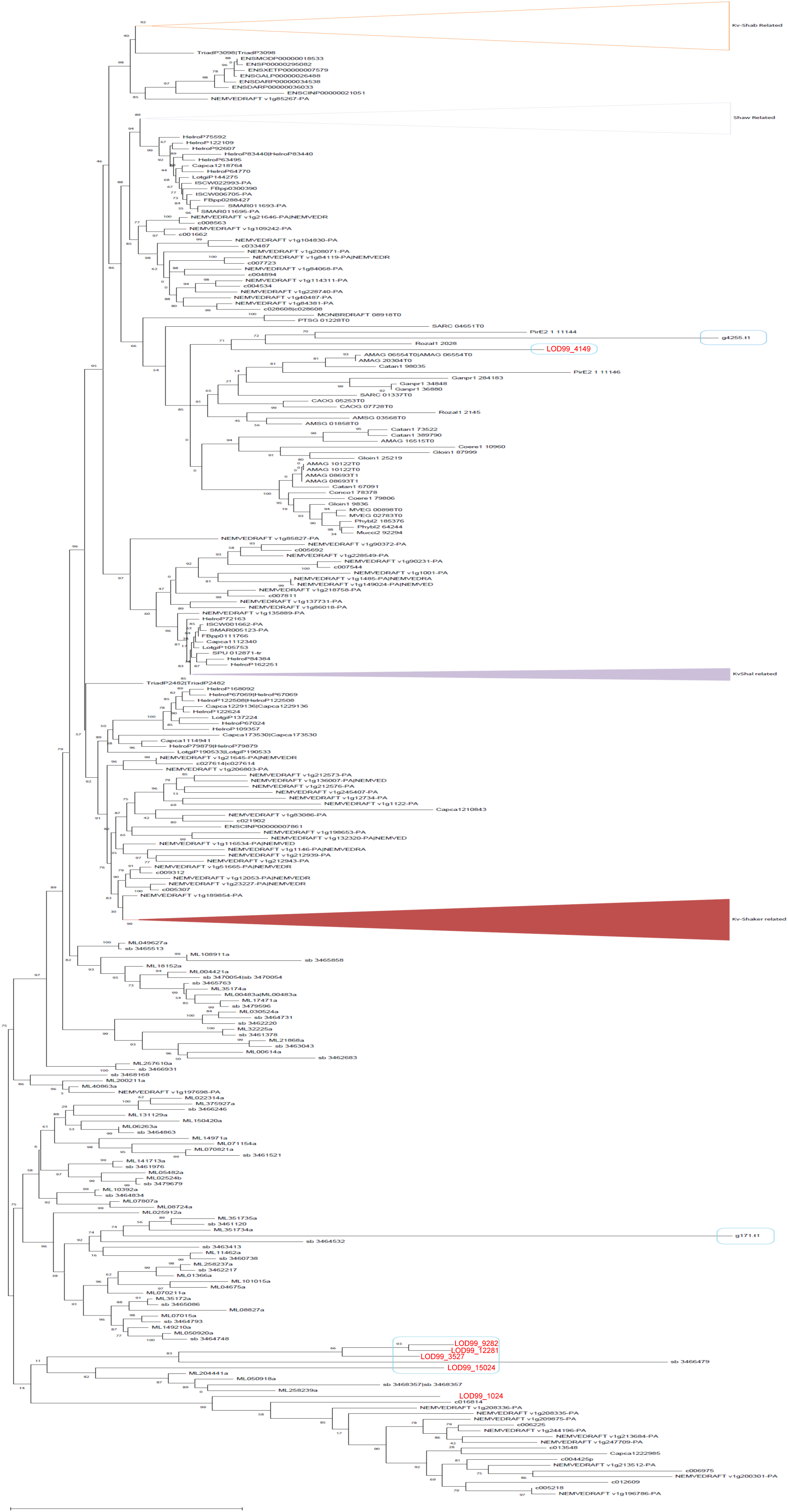
Phylogenetic position of *Oopsacas* Potassium channels.

**Supplementary Table 16:**
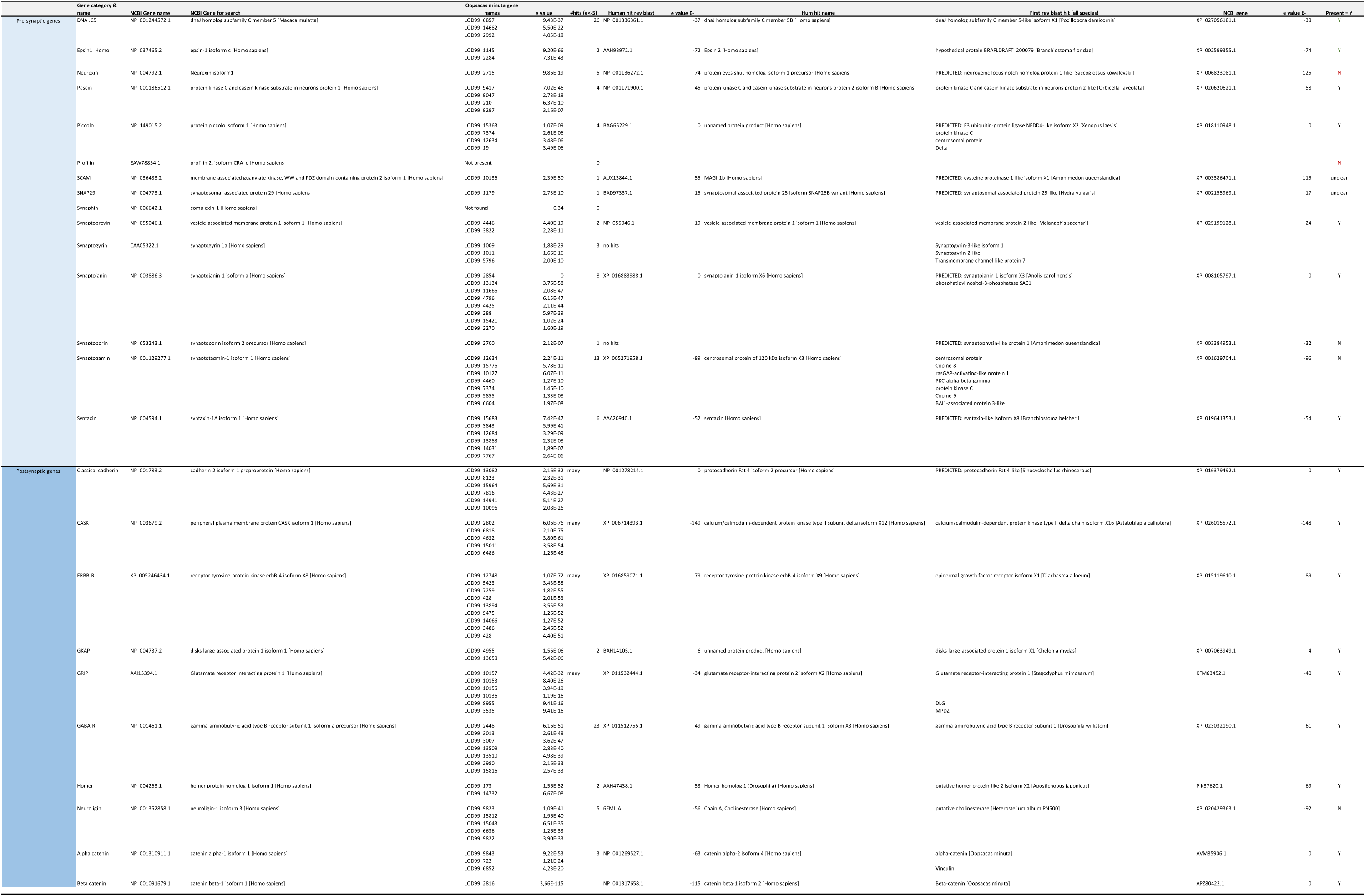
Blast search of genes involved in bilaterian synapses against the predicted proteome of *Oopascas minuta* and reverse blast best hits.

**Supplementary Table 17:**
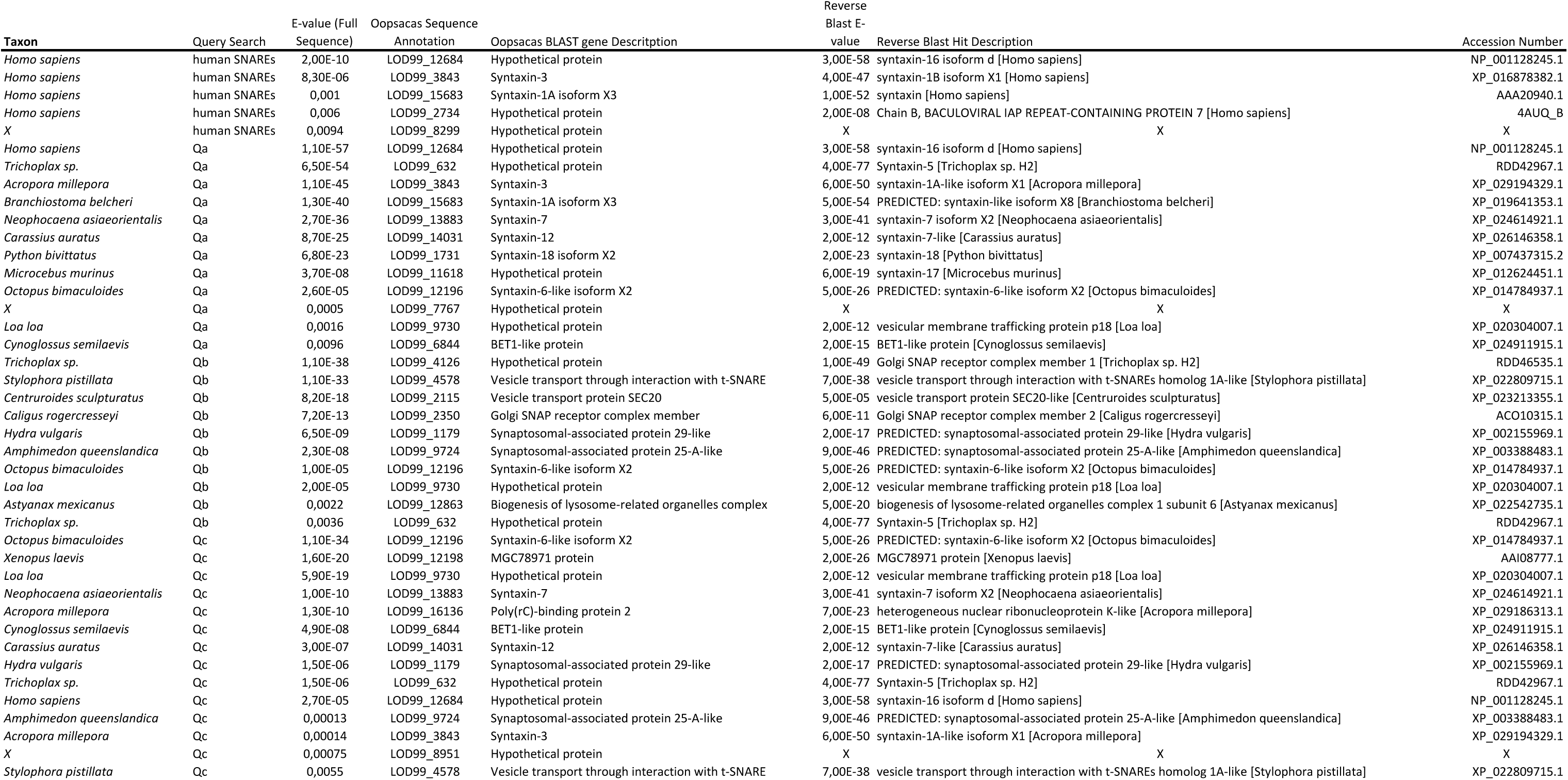
SNARES gene Blast search.

### Survey of genes involved in biosilification

**Supplementary Table 18:** Blast search of genes involved in silica biogenesis against Oopsacas predicted proteome, and reverse blast best hit identity.

| Name of protein | Query ID | Accession number | Hit ID Oosacras | evalue | Reverse Blast ID |
| --- | --- | --- | --- | --- | --- |
| Silicatein | silicatein-like protein [Aulosaccus sp. GV-2009] | ACU86972.1 | NONE |  |  |
| Silintaphin | silintaphin-1 [Suberites domuncula] | CAP16640.1 | NONE |  |  |
| galectin | galectin [Suberites domuncula] | CAD37940.1 | NONE |  |  |
| Cathepsin L | cathepsin L [Sympagella nux] | Own contig | LOD99_12702 | 2.52 × 10<sup>-102</sup> | Cathepsin L |
| Cathepsin L | cathepsin L [Sympagella nux] | Own contig | LOD99_4353 | 2.18 × 10<sup>-100</sup> | Cathepsin L |
| Cathepsin L | cathepsin L [Sympagella nux] | Own contig | LOD99_10364 | 5.08 × 10<sup>-92</sup> | Cathepsin L |
| Cathepsin L | cathepsin L [Sympagella nux] | Own contig | LOD99_11258 | 8.03 × 10<sup>-85</sup> | Cathepsin L |
| Cathepsin L | cathepsin L [Sympagella nux] | Own contig | LOD99_6637 | 3.02 × 10<sup>-84</sup> | Cathepsin L |
| Cathepsin L | cathepsin L [Sympagella nux] | Own contig | LOD99_6632 | 1.86 × 10<sup>-81</sup> | Cathepsin L |
| Cathepsin L | cathepsin L [Sympagella nux] | Own contig | LOD99_5135 | 4.93 × 10<sup>-73</sup> | Cathepsin L |
| Cathepsin L | cathepsin L [Sympagella nux] | Own contig | LOD99_7783 | 2.69 × 10<sup>-72</sup> | Cathepsin L |
| Cathepsin L | cathepsin L [Sympagella nux] | Own contig | LOD99_9107 | 1.42 × 10<sup>-71</sup> | Cathepsin L |
| Cathepsin L | cathepsin L [Sympagella nux] | Own contig | LOD99_29 | 3.92 × 10<sup>-53</sup> | Cathepsin L |
| Cathepsin L | cathepsin L [Sympagella nux] | Own contig | LOD99_14333 | 7.93 × 10<sup>-51</sup> | Cathepsin L |
| Cathepsin L | cathepsin L [Sympagella nux] | Own contig | LOD99_6910 | 1.12 × 10<sup>-45</sup> | Cathepsin L |
| Cathepsin L | cathepsin L [Sympagella nux] | Own contig | LOD99_16078 | 3.21 × 10<sup>-54</sup> | Cathepsin L |
| Cathepsin B | cathepsin B [Sympagella nux] | Own contig | LOD99_2071 | 3.21 × 10<sup>-51</sup> | Cathepsin B |
| Cathepsin B | cathepsin B [Sympagella nux] | Own contig | LOD99_8394 | 4.15 × 10<sup>-100</sup> | Cathepsin B |
| Cathepsin B | cathepsin B [Sympagella nux] | Own contig | LOD99_6927 | 1.35 × 10<sup>-64</sup> | Cathepsin B |
| Cathepsin B | cathepsin B [Sympagella nux] | Own contig | LOD99_2671 | 1.34 × 10<sup>-84</sup> | Cathepsin B |
| Cathepsin B | cathepsin B [Sympagella nux] | Own contig | LOD99_3637 | 2.26 × 10<sup>-100</sup> | Cathepsin B |
| NBCSA cotransporter | natriumbicarbonate silicic acid cotransporter [Suberites domuncula] | CAF32326.1 | LOD99_7310 | 2.52 × 10<sup>-144</sup> | anion exchange protein 2-like [Biomphalaria glabrata] |
| arsB transporter (LSi2) | PREDICTED: putative transporter arsB [Amphimedon queenslandica] | XP_003386169.1 | LOD99_7531 | 2.76 × 10<sup>-99</sup> | arsB transporter |
| arsB transporter (LSi2) | PREDICTED: putative transporter arsB [Amphimedon queenslandica] | XP_003386169.1 | LOD99_12653 | 1.84 × 10<sup>-94</sup> | arsB transporter |
| Glassin | glassin, partial [Euplectella curvstellata] | BAS21353.1 | LOD99_3750 | 1.32 × 10<sup>-54</sup> | Glassin, partial |
| Silicase | CA_SubDo [Suberites domuncula] | SFW10617.1 | LOD99_7560 | 2.57 × 10<sup>-21</sup> | Carbonic anhydrase 20 |
| Silicase | CA_SubDo [Suberites domuncula] | SFW10617.1 | LOD99_966 | 3.45 × 10<sup>-18</sup> | Carbonic anhydrase 1-like |
| Silicase | CA_SubDo [Suberites domuncula] | SFW10617.1 | LOD99_7562 | 1.96 × 10<sup>-15</sup> | Carbonic anhydrase 2-like |
| Silicase | CA_SubDo [Suberites domuncula] | SFW10617.1 | LOD99_7559 | 4.38 × 10<sup>-12</sup> | Alpha carbonic anhydrase 5 |
| Ferretin | ferretin [Amphimedon queenslandica] |  | LOD99_6480 | 5.72 × 10<sup>-70</sup> | Ferretin |
| Chitin synthase | chitin synthase CHS5611, partial [Spongilla lacustris] | AEI55441.1 | LOD99_13043 | 4.08 × 10<sup>-100</sup> | Chitin synthase 6 |

**Supplementary Table 19:**
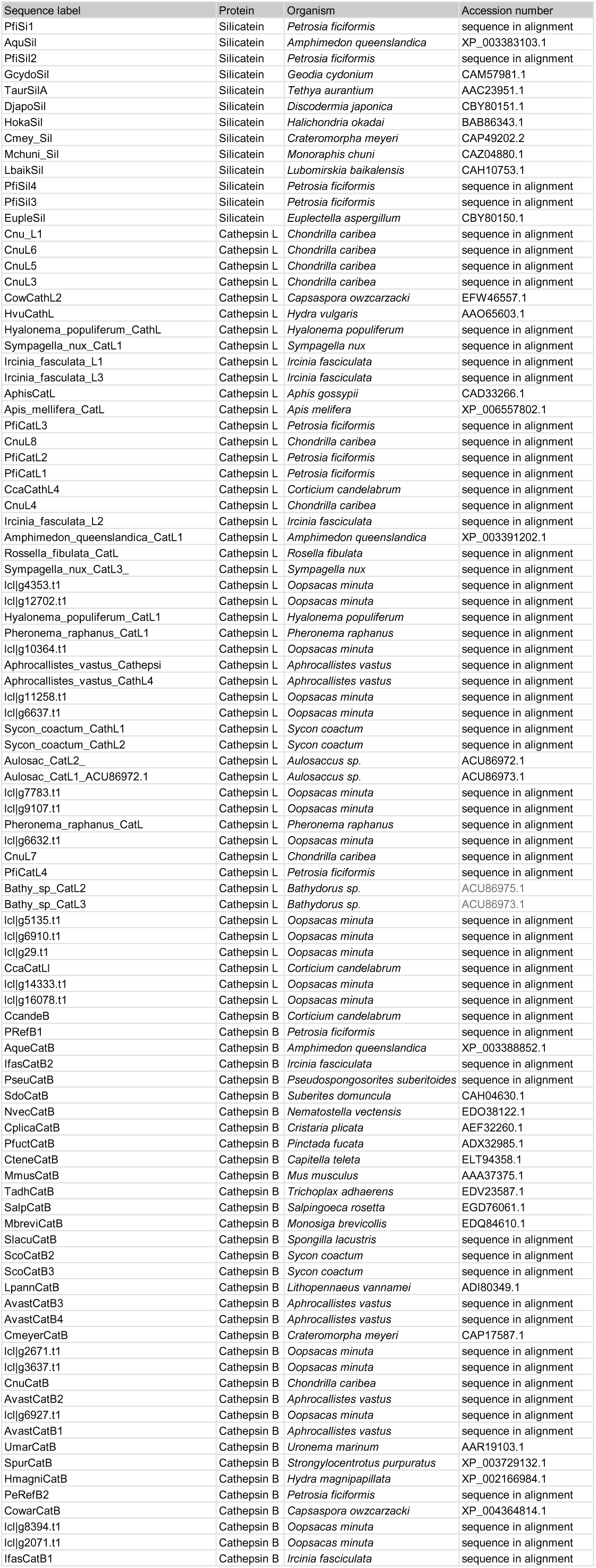
List of sequences used for phylogenetic analyses of silicatein and cathepsin (Figure 5)

### Presence /absence of Core Metazoan genes

**Supplementary Table 20:**
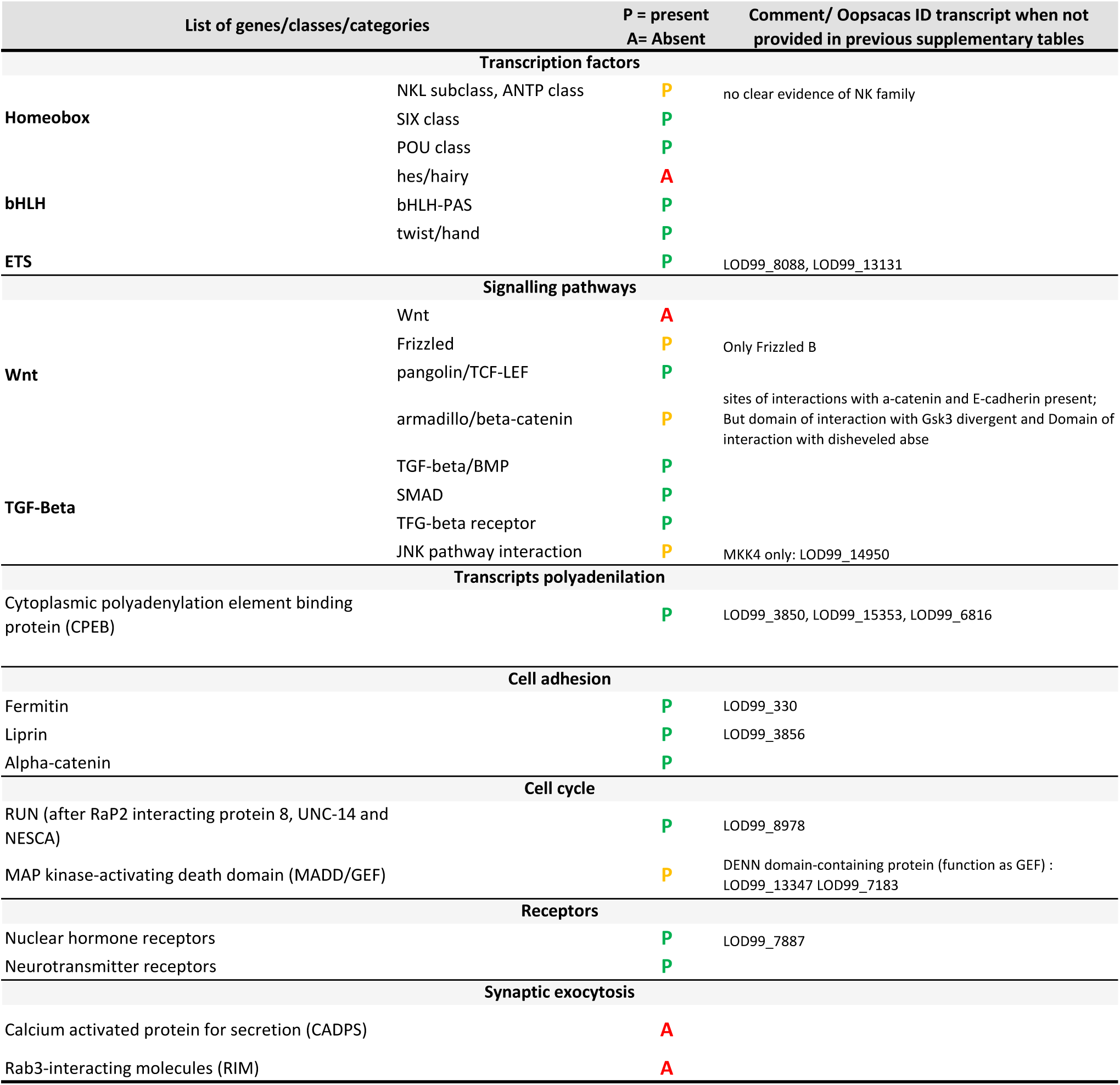
presence/absence of core metazoan innovations (as defined by Paps and Holland, 2018) in *Oopsacas minuta*.

## REFERENCES

1. Dunn CW, Ryan JF. The evolution of animal genomes. Curr Opin Genet Dev. 1 déc 2015;35:25-32.

2. Paps J, Holland PWH. Reconstruction of the ancestral metazoan genome reveals an increase in genomic novelty. Nat Commun. 30 avr 2018;9(1):1730.

3. Renard E, Leys SP, Wörheide G, Borchiellini C. Understanding Animal Evolution: The Added Value of Sponge Transcriptomics and Genomics. BioEssays [Internet]. 1 sept 2018 [cité 28 août 2018];40(9):e1700237(9). Disponible sur: https://onlinelibrary.wiley.com/doi/abs/10.1002/bies.201700237

4. de Mendoza A, Hatleberg WL, Pang K, Leininger S, Bogdanovic O, Pflueger J, et al. Convergent evolution of a vertebrate-like methylome in a marine sponge. Nat Ecol Evol. 2019;3(10):1464-73.

5. Erives A, Fritzsch B. A Screen for Gene Paralogies Delineating Evolutionary Branching Order of Early Metazoa. G3 GenesGenomesGenetics. 26 déc 2019;10(2):811-26.

6. Feuda R, Dohrmann M, Pett W, Philippe H, Rota-Stabelli O, Lartillot N, et al. Improved Modeling of Compositional Heterogeneity Supports Sponges as Sister to All Other Animals. Curr Biol [Internet]. 30 nov 2017 [cité 11 déc 2017];Dec 18;27(24):3864–3870.e4.(0). Disponible sur: http://www.cell.com/current-biology/abstract/S0960-9822(17)31453-7

7. Laumer CE, Fernández R, Lemer S, Combosch D, Kocot KM, Riesgo A, et al. Revisiting metazoan phylogeny with genomic sampling of all phyla. Proc Biol Sci. 10 2019;286(1906):20190831.

8. Redmond AK, McLysaght A. Evidence for sponges as sister to all other animals from partitioned phylogenomics with mixture models and recoding. Nat Commun. 19 mars 2021;12(1):1783.

9. Simion P, Philippe H, Baurain D, Jager M, Richter DJ, Di Franco A, et al. A Large and Consistent Phylogenomic Dataset Supports Sponges as the Sister Group to All Other Animals. Curr Biol CB. 3 avr 2017;27(7):958-67.

10. Whelan NV, Kocot KM, Moroz TP, Mukherjee K, Williams P, Paulay G, et al. Ctenophore relationships and their placement as the sister group to all other animals. Nat Ecol Evol. 9 oct 2017;1(11):1737-46.

11. Dunn CW, Leys SP, Haddock SHD. The hidden biology of sponges and ctenophores. Trends Ecol Evol. mai 2015;30(5):282-91.

12. Dohrmann M, Wörheide G. Dating early animal evolution using phylogenomic data. Sci Rep. 15 juin 2017;7(1):3599.

13. van Soest R, Boury-Esnault N, Hooper J, Rützler K, de Voogd N, Alvarez B, et al. World Porifera Database. Accessed at https://www.marinespecies.org/porifera on 2022-06-30 [Internet]. VLIZ; 2022 [cité 10 juin 2022]. Disponible sur: https://www.marinespecies.org/imis.php?dasid=546&doiid=359

14. Borchiellini C, Kassandra de PM, Amélie V, Rocher C, Ereskovsky A, Vacelet J, et al. Porifera (Sponges): Recent Knowledge and New Perspectives. In: eLS [Internet]. American Cancer Society; 2021 [cité 2 juill 2021]. p. 1-10. Disponible sur: https://onlinelibrary.wiley.com/doi/abs/10.1002/9780470015902.a0029283

15. Colgren J, Nichols SA. The significance of sponges for comparative studies of developmental evolution. WIREs Dev Biol. 2020;9(2):e359.

16. Degnan BM, Adamska M, Richards GS, Larroux C, Leininger S, Bergum B, et al. Porifera. In: Wanninger A, éditeur. Evolutionary Developmental Biology of Invertebrates 1: Introduction, Non-Bilateria, Acoelomorpha, Xenoturbellida, Chaetognatha [Internet]. Vienna: Springer; 2015 [cité 16 avr 2020]. p. 65-106. Disponible sur: https://doi.org/10.1007/978-3-7091-1862-7_4

17. Ereskovsky AV, Renard E, Borchiellini C. Cellular and molecular processes leading to embryo formation in sponges: evidences for high conservation of processes throughout animal evolution. Dev Genes Evol. 1 mars 2013;223(1-2):5-22.

18. Fahey B, Degnan BM. Origin of animal epithelia: insights from the sponge genome: Evolution of epithelia. Evol Dev. nov 2010;12(6):601-17.

19. Fernandez-Valverde SL, Calcino AD, Degnan BM. Deep developmental transcriptome sequencing uncovers numerous new genes and enhances gene annotation in the sponge *Amphimedon queenslandica*. BMC Genomics. 15 mai 2015;16: 387(1):387.

20. Srivastava M, Simakov O, Chapman J, Fahey B, Gauthier MEA, Mitros T, et al. The *Amphimedon queenslandica* genome and the evolution of animal complexity. Nature. 5 août 2010;466(7307):720-6.

21. Adamska M, Degnan BM, Green K, Zwafink C. What sponges can tell us about the evolution of developmental processes. Zool Jena Ger. févr 2011;114(1):1-10.

22. Gazave E, Lapébie P, Richards GS, Brunet F, Ereskovsky AV, Degnan BM, et al. Origin and evolution of the Notch signalling pathway: an overview from eukaryotic genomes. BMC Evol Biol. 13 oct 2009;9:249.

23. Riesgo A, Farrar N, Windsor PJ, Giribet G, Leys SP. The Analysis of Eight Transcriptomes from All Poriferan Classes Reveals Surprising Genetic Complexity in Sponges. Mol Biol Evol. 1 mai 2014;31(5):1102-20.

24. Schenkelaars Q, Fierro-Constain L, Renard E, Hill AL, Borchiellini C. Insights into Frizzled evolution and new perspectives. Evol Dev. avr 2015;17(2):160-9.

25. Schenkelaars Q, Fierro-Constain L, Renard E, Borchiellini C. Retracing the path of planar cell polarity. BMC Evol Biol [Internet]. 2 avr 2016;16:69. Disponible sur: http://www.ncbi.nlm.nih.gov/pmc/articles/PMC4818920/

26. Schenkelaars Q, Pratlong M, Kodjabachian L, Fierro-Constain L, Vacelet J, Le Bivic A, et al. Animal multicellularity and polarity without Wnt signaling. Sci Rep. 13 nov 2017;7(1):15383.

27. Francis WR, Eitel M, R SV, Adamski M, Haddock SH, Krebs S, et al. The Genome Of The Contractile Demosponge *Tethya wilhelma* And The Evolution Of Metazoan Neural Signalling Pathways. bioRxiv. 12 avr 2017;120998.

28. Kenny NJ, Francis WR, Rivera-Vicéns RE, Juravel K, de Mendoza A, Díez-Vives C, et al. Tracing animal genomic evolution with the chromosomal-level assembly of the freshwater sponge *Ephydatia muelleri*. Nat Commun. 27 juill 2020;11(1):3676.

29. 29. Ereskovsky AV, Richter DJ, Lavrov DV, Schippers KJ, Nichols SA. Transcriptome sequencing and delimitation of sympatric *Oscarella* species (*O. carmela* and *O. pearsei* sp. nov) from California, USA. PLoS ONE [Internet]. 11 sept 2017;12(9): e0183002(9). Disponible sur: https://www.ncbi.nlm.nih.gov/pmc/articles/PMC5593202/

30. Nichols SA, Roberts BW, Richter DJ, Fairclough SR, King N. Origin of metazoan cadherin diversity and the antiquity of the classical cadherin/β-catenin complex. Proc Natl Acad Sci U S A. 7 août 2012;109(32):13046-51.

31. Fortunato SAV, Adamski M, Ramos OM, Leininger S, Liu J, Ferrier DEK, et al. Calcisponges have a ParaHox gene and dynamic expression of dispersed NK homeobox genes. Nature. 30 oct 2014;514(7524):620-3.

32. Belahbib H, Renard E, Santini S, Jourda C, Claverie JM, Borchiellini C, et al. New genomic data and analyses challenge the traditional vision of animal epithelium evolution. BMC Genomics [Internet]. 24 mai 2018 [cité 28 août 2018];19, 393. Disponible sur: https://www.ncbi.nlm.nih.gov/pmc/articles/PMC5968619/

33. Fierro-Constaín L, Schenkelaars Q, Gazave E, Haguenauer A, Rocher C, Ereskovsky A, et al. The Conservation of the Germline Multipotency Program, from Sponges to Vertebrates: A Stepping Stone to Understanding the Somatic and Germline Origins. Genome Biol Evol. 1 mars 2017;9(3):474-88.

34. Boury-Esnault N, Vacelet J. Preliminary studies on the organization and development of a hexactinellid sponge from a Mediterranean cave, Oopsacas minuta. Sponges Time Space. 1994;407-16.

35. Vacelet J, Boury-Esnault N, Harmelin JG. Hexactinellid cave, a unique deep-sea habitat in the scuba zone. Deep Sea Res Part Oceanogr Res Pap. 1 juill 1994;41(7):965-73.

36. Leys SP, Mackie GO, Reiswig HM. The Biology of Glass Sponges. In: Advances in Marine Biology [Internet]. Academic Press; 2007 [cité 20 déc 2018]. p. 1-145. Disponible sur: http://www.sciencedirect.com/science/article/pii/S0065288106520012

37. Overstreet RM, Lotz JM. Host–Symbiont Relationships: Understanding the Change from Guest to Pest. Rasputin Eff Commensals Symbionts Become Parasit. 6 janv 2016;3:27-64.

38. Jourda C, Santini S, Rocher C, Le Bivic A, Claverie JM. Draft Genome Sequence of an Alphaproteobacterium Associated with the Mediterranean Sponge *Oscarella lobularis*. Genome Announc [Internet]. 3 sept 2015 [cité 24 mars 2020];3(3(5):e00977–15.). Disponible sur: https://www.ncbi.nlm.nih.gov/pmc/articles/PMC4559732/

39. Thomas T, Moitinho-Silva L, Lurgi M, Björk JR, Easson C, Astudillo-García C, et al. Diversity, structure and convergent evolution of the global sponge microbiome. Nat Commun [Internet]. 16 juin 2016;7. Disponible sur: https://www.ncbi.nlm.nih.gov/pmc/articles/PMC4912640/

40. Roué M, Quévrain E, Domart-Coulon I, Bourguet-Kondracki ML. Assessing calcareous sponges and their associated bacteria for the discovery of new bioactive natural products. Nat Prod Rep. 1 juill 2012;29(7):739-51.

41. Pascelli C, Laffy PW, Kupresanin M, Ravasi T, Webster NS. Morphological characterization of virus-like particles in coral reef sponges. PeerJ. 17 oct 2018;6:e5625.

42. Vacelet J, Gallissian MF. Virus-like particles in cells of the sponge *Verongia cavernicola* (demospongiae, dictyoceratida) and accompanying tissues changes. J Invertebr Pathol. 1 mars 1978;31(2):246-54.

43. Butina TV, Potapov SA, Belykh OI, Belikov SI. Genetic diversity of cyanophages of the myoviridae family as a constituent of the associated community of the Baikal sponge *Lubomirskia baicalensis*. Russ J Genet. 1 mars 2015;51(3):313-7.

44. Laffy PW, Wood-Charlson EM, Turaev D, Weynberg KD, Botté ES, van Oppen MJH, et al. HoloVir: A Workflow for Investigating the Diversity and Function of Viruses in Invertebrate Holobionts. Front Microbiol [Internet]. 9 juin 2016 [cité 2 sept 2020];7. Disponible sur: https://www.ncbi.nlm.nih.gov/pmc/articles/PMC4899465/

45. Laffy PW, Wood-Charlson EM, Turaev D, Jutz S, Pascelli C, Botté ES, et al. Reef invertebrate viromics: diversity, host specificity and functional capacity. Environ Microbiol. 2018;20(6):2125-41.

46. Waldron FM, Stone GN, Obbard DJ. Metagenomic sequencing suggests a diversity of RNA interference-like responses to viruses across multicellular eukaryotes. PLOS Genet. 30 juill 2018;14(7):e1007533.

47. Butina TV, Bukin YS, Khanaev IV, Kravtsova LS, Maikova OO, Tupikin AE, et al. Metagenomic analysis of viral communities in diseased Baikal sponge *Lubomirskia baikalensis*. Limnol Freshw Biol. 5 mars 2019;155-62.

48. Soffer N, Brandt ME, Correa AM, Smith TB, Thurber RV. Potential role of viruses in white plague coral disease. ISME J. févr 2014;8(2):271-83.

49. Dennis TPW, Flynn PJ, de Souza WM, Singer JB, Moreau CS, Wilson SJ, et al. Insights into Circovirus Host Range from the Genomic Fossil Record. J Virol [Internet]. 31 juill 2018 [cité 2 sept 2020];92(16). Disponible sur: https://www.ncbi.nlm.nih.gov/pmc/articles/PMC6069186/

50. Ishijima J, Iwabe N, Masuda Y, Watanabe Y, Matsuda Y. Sponge cytogenetics - mitotic chromosomes of ten species of freshwater sponge. Zoolog Sci. mai 2008;25(5):480-6.

51. Parks DH, Imelfort M, Skennerton CT, Hugenholtz P, Tyson GW. CheckM: assessing the quality of microbial genomes recovered from isolates, single cells, and metagenomes. Genome Res. juill 2015;25(7):1043-55.

52. Haber M, Burgsdorf I, Handley KM, Rubin-Blum M, Steindler L. Genomic Insights Into the Lifestyles of Thaumarchaeota Inside Sponges. Front Microbiol [Internet]. 2021 [cité 10 juin 2022];11. Disponible sur: https://www.frontiersin.org/article/10.3389/fmicb.2020.622824

53. Krupovic M, Makarova KS, Wolf YI, Medvedeva S, Prangishvili D, Forterre P, et al. Integrated mobile genetic elements in Thaumarchaeota. Environ Microbiol. juin 2019;21(6):2056-78.

54. Björk JR, Díez-Vives C, Astudillo-García C, Archie EA, Montoya JM. Vertical transmission of sponge microbiota is inconsistent and unfaithful. Nat Ecol Evol. août 2019;3(8):1172-83.

55. Sharp KH, Eam B, Faulkner DJ, Haygood MG. Vertical transmission of diverse microbes in the tropical sponge *Corticium sp*. Appl Environ Microbiol. janv 2007;73(2):622-9.

56. Webster NS, Taylor MW. Marine sponges and their microbial symbionts: love and other relationships. Environ Microbiol. 2012;14(2):335-46.

57. Martens JH, Barg H, Warren MJ, Jahn D. Microbial production of vitamin B12. Appl Microbiol Biotechnol. mars 2002;58(3):275-85.

58. Roth J, Lawrence J, Bobik T. COBALAMIN (COENZYME B12): Synthesis and Biological Significance. Annu Rev Microbiol. 1 oct 1996;50(1):137-81.

59. Hallam SJ, Konstantinidis KT, Putnam N, Schleper C, Watanabe Yichi, Sugahara J, et al. Genomic analysis of the uncultivated marine crenarchaeote *Cenarchaeum symbiosum*. Proc Natl Acad Sci U S A. 28 nov 2006;103(48):18296-301.

60. Jeffery NW, Jardine CB, Gregory TR. A first exploration of genome size diversity in sponges. Genome. août 2013;56(8):451-6.

61. Simão FA, Waterhouse RM, Ioannidis P, Kriventseva EV, Zdobnov EM. BUSCO: assessing genome assembly and annotation completeness with single-copy orthologs. Bioinforma Oxf Engl. 1 oct 2015;31(19):3210-2.

62. Dohrmann M, Janussen D, Reitner J, Collins AG, Wörheide G, Anderson F. Phylogeny and Evolution of Glass Sponges (Porifera, Hexactinellida). Syst Biol. 1 juin 2008;57(3):388-405.

63. Leys SP. The Choanosome of Hexactinellid Sponges. Invertebr Biol. 1999;118(3):221-35.

64. Leys SP. The Significance of Syncytial Tissues for the Position of the Hexactinellida in the Metazoa. Integr Comp Biol. 1 févr 2003;43(1):19-27.

65. Leys SP, Zaman AK, Boury-Esnault N. Three-dimensional fate mapping of larval tissues through metamorphosis in the glass sponge *Oopsacas minuta*. Invertebr Biol. 2016;135(3):259-72.

66. Abedin M, King N. Diverse evolutionary paths to cell adhesion. Trends Cell Biol. déc 2010;20(12):734-42.

67. Fidler AL, Darris CE, Chetyrkin SV, Pedchenko VK, Boudko SP, Brown KL, et al. Collagen IV and basement membrane at the evolutionary dawn of metazoan tissues. eLife. 18 avr 2017;Apr 18;6:e24176.

68. Leys SP, Riesgo A. Epithelia, an Evolutionary Novelty of Metazoans. J Exp Zoolog B Mol Dev Evol. 1 sept 2012;318(6):438-47.

69. Leys SP, Nichols SA, Adams EDM. Epithelia and integration in sponges. Integr Comp Biol. 1 août 2009;49(2):167-77.

70. Miller PW, Clarke DN, Weis WI, Lowe CJ, Nelson WJ. The Evolutionary Origin of Epithelial Cell-Cell Adhesion Mechanisms. Curr Top Membr. 2013;72:267-311.

71. Renard E, Le Bivic A, Borchiellini C. Origin and Evolution of Epithelial Cell Types [Internet]. Origin and Evolution of Metazoan Cell Types. Leys, S., & Hejnol, A. (Eds.). Origin and Evolution of Metazoan Cell Types (1st ed.). CRC Press.; 2021 [cité 23 avr 2021]. Disponible sur: https://www.taylorfrancis.com/chapters/edit/10.1201/b21831-5/origin-evolution-epithelial-cell-types-emmanuelle-renard-andre-le-bivic-carole-borchiellini

72. Tyler S. Epithelium—The Primary Building Block for Metazoan Complexity1. Integr Comp Biol. 2003;43(1):55-63.

73. Alexopoulos H, Böttger A, Fischer S, Levin A, Wolf A, Fujisawa T, et al. Evolution of gap junctions: the missing link? Curr Biol. 26 oct 2004;14(20):R879-80.

74. Clarke DN, Miller PW, Lowe CJ, Weis WI, Nelson WJ. Characterization of the Cadherin?Catenin Complex of the Sea Anemone *Nematostella vectensis* and Implications for the Evolution of Metazoan Cell?Cell Adhesion. Mol Biol Evol. août 2016;33(8):2016-29.

75. Ganot P, Zoccola D, Tambutté E, Voolstra CR, Aranda M, Allemand D, et al. Structural molecular components of septate junctions in cnidarians point to the origin of epithelial junctions in eukaryotes. Mol Biol Evol. 2015;32(1):44-62.

76. Takaku Y, Hwang JS, Wolf A, Böttger A, Shimizu H, David CN, et al. Innexin gap junctions in nerve cells coordinate spontaneous contractile behavior in *Hydra* polyps. Sci Rep [Internet]. 7 janv 2014 [cité 16 nov 2018];4. Disponible sur: https://www.ncbi.nlm.nih.gov/pmc/articles/PMC3882753/

77. Tucker RP, Adams JC. Chapter Eight - Adhesion Networks of Cnidarians: A Postgenomic View. In: Jeon KW, éditeur. International Review of Cell and Molecular Biology [Internet]. Academic Press; 2014 [cité 19 juill 2019]. p. 323-77. Disponible sur: http://www.sciencedirect.com/science/article/pii/B9780128000977000087

78. Leys SP, Cheung E, Boury-Esnault N. Embryogenesis in the glass sponge *Oopsacas minuta*: Formation of syncytia by fusion of blastomeres. Integr Comp Biol. 1 avr 2006;46(2):104-17.

79. Riesgo A, Farrar N, Windsor PJ, Giribet G, Leys SP. The Analysis of Eight Transcriptomes from All Poriferan Classes Reveals Surprising Genetic Complexity in Sponges. Mol Biol Evol. 1 mai 2014;31(5):1102-20.

80. Mackie GO, Singla CL. Studies on hexactinellid sponges - Studies on hexactinellid sponges. I. Histology of *Rhabdocalyptus dawsoni* (Lambe, 1873). Phil Trans R Soc Lond B. 5 juill 1983;301(1107):365-400.

81. De Pascalis C, Etienne-Manneville S. Single and collective cell migration: the mechanics of adhesions. Mol Biol Cell. 7 juill 2017;28(14):1833-46.

82. Magie CR, Martindale MQ. Cell-Cell Adhesion in the Cnidaria: Insights Into the Evolution of Tissue Morphogenesis. Biol Bull. 1 juin 2008;214(3):218-32.

83. Miller PW, Pokutta S, Mitchell JM, Chodaparambil JV, Clarke DN, Nelson WJ, et al. Analysis of a vinculin homolog in a sponge (phylum Porifera) reveals that vertebrate-like cell adhesions emerged early in animal evolution. J Biol Chem. 27 juill 2018;293(30):11674-86.

84. Walko G, Castañón MJ, Wiche G. Molecular architecture and function of the hemidesmosome. Cell Tissue Res. 2015;360(2):363-78.

85. Mitchell JM, Nichols SA. Diverse cell junctions with unique molecular composition in tissues of a sponge (Porifera). EvoDevo [Internet]. 29 oct 2019 [cité 25 mars 2020];10:26. Disponible sur: https://www.ncbi.nlm.nih.gov/pmc/articles/PMC6820919/

86. Nichols SA, Dirks W, Pearse JS, King N. Early evolution of animal cell signaling and adhesion genes. Proc Natl Acad Sci U S A. 15 août 2006;103(33):12451-6.

87. Fidler AL, Darris CE, Chetyrkin SV, Pedchenko VK, Boudko SP, Brown KL, et al. Collagen IV and basement membrane at the evolutionary dawn of metazoan tissues. eLife. 18 avr 2017;6.

88. Fidler AL, Boudko SP, Rokas A, Hudson BG. The triple helix of collagens – an ancient protein structure that enabled animal multicellularity and tissue evolution. J Cell Sci. 1 avr 2018;131(7):jcs203950.

89. Medwig TN, Matus DQ. Breaking down barriers: the evolution of cell invasion. Curr Opin Genet Dev. déc 2017;47:33-40.

90. Pozzi A, Yurchenco PD, Iozzo RV. The nature and biology of basement membranes. Matrix Biol J Int Soc Matrix Biol. janv 2017;57-58:1-11.

91. Sekiguchi R, Yamada KM. Chapter Four - Basement Membranes in Development and Disease. In: Litscher ES, Wassarman PM, éditeurs. Current Topics in Developmental Biology [Internet]. Academic Press; 2018 [cité 30 nov 2018]. p. 143-91. (Extracellular Matrix and Egg Coats; vol. 130). Disponible sur: http://www.sciencedirect.com/science/article/pii/S0070215318300371

92. Fahey B, Degnan BM. Origin and Evolution of Laminin Gene Family Diversity. Mol Biol Evol. 1 juill 2012;29(7):1823-36.

93. Whittaker CA, Bergeron KF, Whittle J, Brandhorst BP, Burke RD, Hynes RO. The echinoderm adhesome. Dev Biol. 1 déc 2006;300(1):252-66.

94. Ereskovsky A. The Comparative Embryology of Sponges. In 2010. p. 209-30.

95. Boute N, Exposito JY, Boury-Esnault N, Vacelet J, Noro N, Miyazaki K, et al. Type IV collagen in sponges, the missing link in basement membrane ubiquity*. Biol Cell. 1 janv 1996;88(1-2):37-44.

96. Srivastava M, Simakov O, Chapman J, Fahey B, Gauthier MEA, Mitros T, et al. The Amphimedon queenslandica genome and the evolution of animal complexity. Nature. 5 août 2010;466(7307):720-6.

97. Boutin C, Kodjabachian L. Biology of multiciliated cells. Curr Opin Genet Dev. 1 juin 2019;56:1-7.

98. Basquin C, Orfila AM, Azimzadeh J. Chapter 13 - The planarian *Schmidtea mediterranea* as a model for studying motile cilia and multiciliated cells. In: Basto R, Marshall WF, éditeurs. Methods in Cell Biology [Internet]. Academic Press; 2015 [cité 8 juill 2020]. p. 243-62. (Methods in Cilia & Flagella; vol. 127). Disponible sur: http://www.sciencedirect.com/science/article/pii/S0091679X1500028X

99. Rink JC, Vu HTK, Sánchez Alvarado A. The maintenance and regeneration of the planarian excretory system are regulated by EGFR signaling. Dev Camb Engl. 1 sept 2011;138(17):3769-80.

100. Rompolas P, Azimzadeh J, Marshall WF, King SM. Chapter Twelve - Analysis of Ciliary Assembly and Function in Planaria. In: Marshall WF, éditeur. Methods in Enzymology [Internet]. Academic Press; 2013 [cité 8 juill 2020]. p. 245-64. (Cilia, Part B; vol. 525). Disponible sur: http://www.sciencedirect.com/science/article/pii/B9780123979445000122

101. Tamm S, Tamm SL. Development of macrociliary cells in Beroe. I. Actin bundles and centriole migration. J Cell Sci. 1 janv 1988;89(1):67-80.

102. Thi-Kim Vu H, Rink JC, McKinney SA, McClain M, Lakshmanaperumal N, Alexander R, et al. Stem cells and fluid flow drive cyst formation in an invertebrate excretory organ. eLife [Internet]. 2015 [cité 8 juill 2020];4. Disponible sur: https://www.ncbi.nlm.nih.gov/pmc/articles/PMC4500094/

103. Tyler S. Development of cilia in embryos of the turbellarian *Macrostomum*. Hydrobiologia. 1 oct 1981;84(1):231-9.

104. Hsu J, Sage J. Novel functions for the transcription factor E2F4 in development and disease. Cell Cycle. 1 déc 2016;15(23):3183-90.

105. Arquint C, Nigg EA. The PLK4–STIL–SAS-6 module at the core of centriole duplication. Biochem Soc Trans. 15 oct 2016;44(5):1253-63.

106. Azimzadeh J, Wong ML, Downhour DM, Alvarado AS, Marshall WF. Centrosome Loss in the Evolution of Planarians. Science. 27 janv 2012;335(6067):461-3.

107. BOURY-ESNAULT N, EFREMOVA S, BÉZAC C, VACELET J. Reproduction of a hexactinellid sponge: first description of gastrulation by cellular delamination in the Porifera. Invertebr Reprod Dev. 1 juill 1999;35(3):187-201.

108. Riesgo A, Taylor C, Leys SP. Reproduction in a carnivorous sponge: the significance of the absence of an aquiferous system to the sponge body plan. Evol Dev. 2007;9(6):618-31.

109. Vij S, Rink JC, Ho HK, Babu D, Eitel M, Narasimhan V, et al. Evolutionarily Ancient Association of the FoxJ1 Transcription Factor with the Motile Ciliogenic Program. PLOS Genet. 8 nov 2012;8(11):e1003019.

110. Mendoza A de, Sebé-Pedrós A, Šestak MS, Matejčić M, Torruella G, Domazet-Lošo T, et al. Transcription factor evolution in eukaryotes and the assembly of the regulatory toolkit in multicellular lineages. Proc Natl Acad Sci. 10 déc 2013;110(50):E4858-66.

111. Biesalski HK, Doepner G, Tzimas G, Gamulin V, Schröder HC, Batel R, et al. Modulation of myb gene expression in sponges by retinoic acid. Oncogene. sept 1992;7(9):1765-74.

112. Nabais C, Peneda C, Bettencourt-Dias M. Evolution of centriole assembly. Curr Biol. 18 mai 2020;30(10):R494-502.

113. Adamska M. Developmental Signalling and Emergence of Animal Multicellularity. In: Ruiz-Trillo I, Nedelcu AM, éditeurs. Evolutionary Transitions to Multicellular Life: Principles and mechanisms [Internet]. Dordrecht: Springer Netherlands; 2015 [cité 8 juill 2020]. p. 425-50. (Advances in Marine Genomics). Disponible sur: https://doi.org/10.1007/978-94-017-9642-2_20g

114. Babonis LS, Martindale MQ. Phylogenetic evidence for the modular evolution of metazoan signalling pathways. Philos Trans R Soc B Biol Sci [Internet]. 5 févr 2017 [cité 30 nov 2018];372(1713):20150477.(1713). Disponible sur: https://www.ncbi.nlm.nih.gov/pmc/articles/PMC5182411/

115. Brunet T, King N. The origin of animal multicellularity and cell differentiation. Dev Cell. 23 oct 2017;43(2):124-40.

116. Ispolatov I, Ackermann M, Doebeli M. Division of labour and the evolution of multicellularity. Proc Biol Sci. 7 mai 2012;279(1734):1768-76.

117. Niklas KJ. The evolutionary-developmental origins of multicellularity. Am J Bot. 1 janv 2014;101(1):6-25.

118. Richards GS, Degnan BM. The dawn of developmental signaling in the metazoa. Cold Spring Harb Symp Quant Biol. 2009;74:81-90.

119. Sebé-Pedrós A, de Mendoza A. Transcription Factors and the Origin of Animal Multicellularity. In: Ruiz-Trillo I, Nedelcu AM, éditeurs. Evolutionary Transitions to Multicellular Life: Principles and mechanisms [Internet]. Dordrecht: Springer Netherlands; 2015 [cité 8 juill 2020]. p. 379-94. (Advances in Marine Genomics). Disponible sur: https://doi.org/10.1007/978-94-017-9642-2_18

120. Windsor Reid PJ, Matveev E, McClymont A, Posfai D, Hill AL, Leys SP. Wnt signaling and polarity in freshwater sponges. BMC Evol Biol. 2 févr 2018;18(1):12.

121. Chang ES, Neuhof M, Rubinstein ND, Diamant A, Philippe H, Huchon D, et al. Genomic insights into the evolutionary origin of Myxozoa within Cnidaria. Proc Natl Acad Sci U S A. 1 déc 2015;112(48):14912-7.

122. Loh KM, van Amerongen R, Nusse R. Generating Cellular Diversity and Spatial Form: Wnt Signaling and the Evolution of Multicellular Animals. Dev Cell. 26 sept 2016;38(6):643-55.

123. Oka Y, Saraiva LR, Kwan YY, Korsching SI. The fifth class of Galpha proteins. Proc Natl Acad Sci U S A. 3 févr 2009;106(5):1484-9.

124. Lokits AD, Indrischek H, Meiler J, Hamm HE, Stadler PF. Tracing the evolution of the heterotrimeric G protein α subunit in Metazoa. BMC Evol Biol [Internet]. 11 avr 2018 [cité 2 juill 2020];18. Disponible sur: https://www.ncbi.nlm.nih.gov/pmc/articles/PMC5896119/

125. Bastiani CA, Gharib S, Simon MI, Sternberg PW. *Caenorhabditis elegans* Galphaq regulates egg-laying behavior via a PLCbeta-independent and serotonin-dependent signaling pathway and likely functions both in the nervous system and in muscle. Genetics. déc 2003;165(4):1805-22.

126. Katanayeva N, Kopein D, Portmann R, Hess D, Katanaev VL. Competing activities of heterotrimeric G proteins in Drosophila wing maturation. PloS One. 23 août 2010;5(8):e12331.

127. Macias-Muñoz A, Murad R, Mortazavi A. Molecular evolution and expression of opsin genes in *Hydra vulgaris*. BMC Genomics [Internet]. 17 déc 2019 [cité 2 juill 2020];20. Disponible sur: https://www.ncbi.nlm.nih.gov/pmc/articles/PMC6918707/

128. Wright SC, Cañizal MCA, Benkel T, Simon K, Le Gouill C, Matricon P, et al. FZD5 is a Gαq-coupled receptor that exhibits the functional hallmarks of prototypical GPCRs. Sci Signal. 4 déc 2018;11(559):eaar5536.

129. Jones S. An overview of the basic helix-loop-helix proteins. Genome Biol. 2004;5(6):226.

130. Fortunato SAV, Vervoort M, Adamski M, Adamska M. Conservation and divergence of bHLH genes in the calcisponge Sycon ciliatum. EvoDevo. 2016;7:23.

131. Richards GS, Simionato E, Perron M, Adamska M, Vervoort M, Degnan BM. Sponge genes provide new insight into the evolutionary origin of the neurogenic circuit. Curr Biol CB. 5 août 2008;18(15):1156-61.

132. Simionato E, Ledent V, Richards G, Thomas-Chollier M, Kerner P, Coornaert D, et al. Origin and diversification of the basic helix-loop-helix gene family in metazoans: insights from comparative genomics. BMC Evol Biol. 2 mars 2007;7(1):33.

133. Bürglin TR, Affolter M. Homeodomain proteins: an update. Chromosoma. 2016;125:497-521.

134. Ferrier DEK. Evolution of Homeobox Gene Clusters in Animals: The Giga-Cluster and Primary vs. Secondary Clustering. Front Ecol Evol [Internet]. 2016 [cité 8 juill 2020];4. Disponible sur: https://www.frontiersin.org/articles/10.3389/fevo.2016.00036/full

135. Garcia-Fernàndez J. The genesis and evolution of homeobox gene clusters. Nat Rev Genet. déc 2005;6(12):881-92.

136. Larroux C, Fahey B, Degnan SM, Adamski M, Rokhsar DS, Degnan BM. The NK Homeobox Gene Cluster Predates the Origin of Hox Genes. Curr Biol. 17 avr 2007;17(8):706-10.

137. Thomas-Chollier M, Martinez P. Origin of Metazoan Patterning Systems and the Role of ANTP-Class Homeobox Genes. In: eLS [Internet]. American Cancer Society; 2016 [cité 5 juill 2021]. p.1-10. Disponible sur: https://onlinelibrary.wiley.com/doi/abs/10.1002/9780470015902.a0022852.pub2

138. Fortunato SA, Leininger S, Adamska M. Evolution of the Pax-Six-Eya-Dach network: the calcisponge case study. EvoDevo. 23 juin 2014;5(1):23.

139. Ryan JF, Pang K, Mullikin JC, Martindale MQ, Baxevanis AD, NISC Comparative Sequencing Program. The homeodomain complement of the ctenophore Mnemiopsis leidyi suggests that Ctenophora and Porifera diverged prior to the ParaHoxozoa. EvoDevo. 4 oct 2010;1(1):9.

140. Elliott GRD, Leys SP. Coordinated contractions effectively expel water from the aquiferous system of a freshwater sponge. J Exp Biol. 1 nov 2007;210(21):3736-48.

141. Leys SP. Elements of a « nervous system » in sponges. J Exp Biol. 15 févr 2015;218(4):581-91.

142. Leys SP, Meech RW. Physiology of coordination in sponges. Can J Zool. 2006;84(2):288-306.

143. Leys SP, Mah JL, McGill PR, Hamonic L, De Leo FC, Kahn AS. Sponge Behavior and the Chemical Basis of Responses: A Post-Genomic View. Integr Comp Biol. 01 2019;59(4):751-64.

144. Mah JL, Leys SP. Think like a sponge: The genetic signal of sensory cells in sponges. Dev Biol. 01 2017;431(1):93-100.

145. Maldonado M. The ecology of the sponge larva. Can J Zool. 2006;84(2):175-94.

146. Nickel M. Evolutionary emergence of synaptic nervous systems: what can we learn from the non-synaptic, nerveless Porifera? Invertebr Biol. 2010;129(1):1-16.

147. Renard E, Vacelet J, Gazave E, Lapébie P, Borchiellini C, Ereskovsky AV. Origin of the neuro-sensory system: new and expected insights from sponges. Integr Zool. sept 2009;4(3):294-308.

148. Simpson TL. The cell biology of sponges. New York: Springer-Verlag; 1984. xix, 662 p.

149. Leys SP, Degnan BM. Cytological basis of photoresponsive behavior in a sponge larva. Biol Bull. déc 2001;201(3):323-38.

150. Rivera AS, Ozturk N, Fahey B, Plachetzki DC, Degnan BM, Sancar A, et al. Blue-light-receptive cryptochrome is expressed in a sponge eye lacking neurons and opsin. J Exp Biol. 15 avr 2012;215(8):1278-86.

151. Leys SP, Mackie GO. Electrical recording from a glass sponge. Nature. mai 1997;387(6628):29-30.

152. Leys SP, Mackie GO, Meech RW. Impulse conduction in a sponge. J Exp Biol. mai 1999;202 (Pt 9):1139-50.

153. Leys SP. Comparative study of spiculogenesis in demosponge and hexactinellid larvae. Microsc Res Tech. 2003;62(4):300-11.

154. Shimizu K, Amano T, Bari MR, Weaver JC, Arima J, Mori N. Glassin, a histidine-rich protein from the siliceous skeletal system of the marine sponge *Euplectella*, directs silica polycondensation. Proc Natl Acad Sci U S A. 15 sept 2015;112(37):11449-54.

155. Shimizu K, Kobayashi H, Nishi M, Tsukahara M, Bito T, Arima J. Exploration of Genes Associated with Sponge Silicon Biomineralization in the Whole Genome Sequence of the Hexactinellid *Euplectella curvistellata*. In: Endo K, Kogure T, Nagasawa H, éditeurs. Biomineralization. Singapore: Springer; 2018. p. 147-53.

156. Wang X, Schloßmacher U, Wiens M, Batel R, Schröder HC, Müller WEG. Silicateins, silicatein interactors and cellular interplay in sponge skeletogenesis: formation of glass fiber-like spicules. FEBS J. 2012;279(10):1721-36.

157. Woodland W. Memoirs: Studies in Spicule Formation: VIII.--Some Observations on the Scieroblastic Development of Hexactinellid and other Siliceous Sponge Spicules. J Cell Sci. 1 janv 1908;s2-52(205):139-57.

158. Riesgo A, Maldonado M, López-Legentil S, Giribet G. A Proposal for the Evolution of Cathepsin and Silicatein in Sponges. J Mol Evol. 1 juin 2015;80(5):278-91.

159. Müller WEG, Eckert C, Kropf K, Wang X, Schloßmacher U, Seckert C, et al. Formation of giant spicules in the deep-sea hexactinellid *Monorhaphis chuni* (Schulze 1904): electron-microscopic and biochemical studies. Cell Tissue Res. 1 août 2007;329(2):363-78.

160. Müller WEG, Schlossmacher U, Wang X, Boreiko A, Brandt D, Wolf SE, et al. Poly(silicate)-metabolizing silicatein in siliceous spicules and silicasomes of demosponges comprises dual enzymatic activities (silica polymerase and silica esterase). FEBS J. janv 2008;275(2):362-70.

161. Müller WEG, Wang X, Kropf K, Boreiko A, Schlossmacher U, Brandt D, et al. Silicatein expression in the hexactinellid *Crateromorpha meyeri*: the lead marker gene restricted to siliceous sponges. Cell Tissue Res. août 2008;333(2):339-51.

162. Müller WEG, Wang X, Burghard Z, Bill J, Krasko A, Boreiko A, et al. Bio-sintering processes in hexactinellid sponges: fusion of bio-silica in giant basal spicules from Monorhaphis chuni. J Struct Biol. déc 2009;168(3):548-61.

163. Veremeichik GN, Shkryl YN, Bulgakov VP, Shedko SV, Kozhemyako VB, Kovalchuk SN, et al. Occurrence of a silicatein gene in glass sponges (Hexactinellida: Porifera). Mar Biotechnol N Y N. août 2011;13(4):810-9.

164. Schröder HC, Perović-Ottstadt S, Rothenberger M, Wiens M, Schwertner H, Batel R, et al. Silica transport in the demosponge *Suberites domuncula:* fluorescence emission analysis using the PDMPO probe and cloning of a potential transporter. Biochem J. 1 août 2004;381(Pt 3):665-73.

165. Maldonado M, López-Acosta M, Beazley L, Kenchington E, Koutsouveli V, Riesgo A. Cooperation between passive and active silicon transporters clarifies the ecophysiology and evolution of biosilicification in sponges. Sci Adv. juill 2020;6(28):eaba9322.

166. Marron AO, Ratcliffe S, Wheeler GL, Goldstein RE, King N, Not F, et al. The Evolution of Silicon Transport in Eukaryotes. Mol Biol Evol. 2016;33(12):3226-48.

167. Smith G, Manzano-Marín A, Reyes-Prieto M, Antunes CSR, Ashworth V, Goselle ON, et al. Human Follicular Mites: Ectoparasites Becoming Symbionts. Mol Biol Evol. 1 juin 2022;39(6):msac125.

168. Yahalomi D, Atkinson SD, Neuhof M, Chang ES, Philippe H, Cartwright P, et al. A cnidarian parasite of salmon (Myxozoa: Henneguya) lacks a mitochondrial genome. Proc Natl Acad Sci U S A. 10 mars 2020;117(10):5358-63.

169. Burke M, Scholl EH, Bird DM, Schaff JE, Colman SD, Crowell R, et al. The plant parasite *Pratylenchus coffeae* carries a minimal nematode genome. Nematology. 2015;17(6):621-37.

170. Rocher C, Vernale A, Fierro-Constain L, Sejourne N, Chenesseau S, Marschal C, et al. The buds of *Oscarella lobularis* (Porifera): a new convenient model for sponge cell and developmental biology. bioRxiv. 23 juin 2020;2020.06.23.167296.

171. Koren S, Walenz BP, Berlin K, Miller JR, Bergman NH, Phillippy AM. Canu: scalable and accurate long-read assembly via adaptive k-mer weighting and repeat separation. Genome Res. mai 2017;27(5):722-36.

172. Langmead B, Wilks C, Antonescu V, Charles R. Scaling read aligners to hundreds of threads on general-purpose processors. Bioinforma Oxf Engl. 01 2019;35(3):421-32.

173. Bankevich A, Nurk S, Antipov D, Gurevich AA, Dvorkin M, Kulikov AS, et al. SPAdes: a new genome assembly algorithm and its applications to single-cell sequencing. J Comput Biol J Comput Mol Cell Biol. mai 2012;19(5):455-77.

174. Zhu W, Lomsadze A, Borodovsky M. Ab initio gene identification in metagenomic sequences. Nucleic Acids Res. juill 2010;38(12):e132.

175. Altschul SF, Gish W, Miller W, Myers EW, Lipman DJ. Basic local alignment search tool. J Mol Biol. 5 oct 1990;215(3):403-10.

176. Boetzer M, Henkel CV, Jansen HJ, Butler D, Pirovano W. Scaffolding pre-assembled contigs using SSPACE. Bioinforma Oxf Engl. 15 févr 2011;27(4):578-9.

177. Walker BJ, Abeel T, Shea T, Priest M, Abouelliel A, Sakthikumar S, et al. Pilon: an integrated tool for comprehensive microbial variant detection and genome assembly improvement. PloS One. 2014;9(11):e112963.

178. Nadalin F, Vezzi F, Policriti A. GapFiller: a de novo assembly approach to fill the gap within paired reads. BMC Bioinformatics. 2012;13 Suppl 14:S8.

179. Quevillon E, Silventoinen V, Pillai S, Harte N, Mulder N, Apweiler R, et al. InterProScan: protein domains identifier. Nucleic Acids Res. 1 juill 2005;33(Web Server issue):W116–120.

180. Marchler-Bauer A, Lu S, Anderson JB, Chitsaz F, Derbyshire MK, DeWeese-Scott C, et al. CDD: a Conserved Domain Database for the functional annotation of proteins. Nucleic Acids Res. 1 janv 2011;39(suppl_1):D225-9.

181. Käll L, Krogh A, Sonnhammer ELL. A combined transmembrane topology and signal peptide prediction method. J Mol Biol. 14 mai 2004;338(5):1027-36.

182. Eddy SR. Accelerated Profile HMM Searches. PLOS Comput Biol. 20 oct 2011;7(10):e1002195.

183. Chan PP, Lowe TM. tRNAscan-SE: Searching for tRNA Genes in Genomic Sequences. Methods Mol Biol Clifton NJ. 2019;1962:1-14.

184. Lowe TM, Chan PP. tRNAscan-SE On-line: integrating search and context for analysis of transfer RNA genes. Nucleic Acids Res. 08 2016;44(W1):W54–57.

185. Kim D, Pertea G, Trapnell C, Pimentel H, Kelley R, Salzberg SL. TopHat2: accurate alignment of transcriptomes in the presence of insertions, deletions and gene fusions. Genome Biol. 25 avr 2013;14(4):R36.

186. Hoff KJ, Lomsadze A, Borodovsky M, Stanke M. Whole-Genome Annotation with BRAKER. Methods Mol Biol Clifton NJ. 2019;1962:65-95.

187. Stanke M, Diekhans M, Baertsch R, Haussler D. Using native and syntenically mapped cDNA alignments to improve de novo gene finding. Bioinforma Oxf Engl. 1 mars 2008;24(5):637-44.

188. Glöckner FO, Fuchs BM, Amann R. Bacterioplankton compositions of lakes and oceans: a first comparison based on fluorescence in situ hybridization. Appl Environ Microbiol. août 1999;65(8):3721-6.

189. Teira E, Reinthaler T, Pernthaler A, Pernthaler J, Herndl GJ. Combining Catalyzed Reporter Deposition-Fluorescence In Situ Hybridization and Microautoradiography To Detect Substrate Utilization by Bacteria and Archaea in the Deep Ocean. Appl Environ Microbiol. juill 2004;70(7) :4411-4.

